# Fluorinated trehalose analogues for cell surface engineering and imaging of *Mycobacterium tuberculosis*

**DOI:** 10.1101/2024.01.30.577379

**Authors:** Collette S. Guy, James A. Gott, Jonathan Ramírez-Cárdenas, Christopher de Wolf, Christopher M. Furze, Geoff West, Juan C. Muñoz-García, Jesus Angulo, Elizabeth Fullam

**Author notes:** To whom correspondence should be addressed: Elizabeth Fullam, School of Life Sciences, University of Warwick, Coventry, CV4 7AL, United Kingdom.

## Abstract

The sensitive, rapid and accurate diagnosis of *Mycobacterium tuberculosis* (*Mtb*) infection is a central challenge in controlling the global tuberculosis (TB) pandemic. Yet the detection of mycobacteria is often made difficult by the low sensitivity of current diagnostic tools, with over 3.6 million TB cases missed each year. To overcome these limitations there is an urgent need for next-generation TB diagnostic technologies. Here we report the use of a discrete panel of native ^19^F-trehalose (F-Tre) analogues to label and directly visualise *Mtb* by exploiting the uptake of fluorine-modified trehalose analogues *via* the mycobacterial trehalose LpqY-SugABC ATP-binding cassette (ABC) importer. We discovered the extent of modified F-Tre uptake correlates with LpqY substrate recognition and characterisation of the interacting sites by saturation transfer difference NMR coupled with molecular dynamics provides a unique glimpse into the molecular basis of fluorine-modified trehalose import in *Mtb*. Lipid profiling demonstrated that F-Tre analogues modified at positions 2, 3 and 6 are incorporated into mycobacterial cell-surface trehalose-containing glycolipids. This rapid one-step labelling approach facilitates the direct visualisation of F-Tre-labelled *Mtb* by focused ion beam (FIB) secondary ion mass spectrometry (SIMS), enabling pathogen specific detection. Collectively, our findings highlight that F-Tre analogues have potential as tools to probe and unravel *Mtb* biology and can be exploited to detect and image TB.

## Introduction

*Mycobacterium tuberculosis* (*Mtb*), the causative agent of tuberculosis (TB), continues to threaten humanity and remains a major health challenge, having claimed over 1.3 million lives in 2022.^1^ Efforts to control the global TB pandemic have been jeopardised by the emergence of drug-resistant *Mtb* strains that are more complicated to treat and, in some instances, incurable.^1, 2^ Early diagnosis is crucial for the global management of TB since the timely intervention with antitubercular treatment regimens has the potential to prevent transmission, limit the emergence and spread of antibiotic resistant strains and save lives, yet each year over million individuals do not receive treatment compounding the TB burden^1, 3, 4^. The current World Health Organisation approved TB diagnostics are limited to three main methods: microscopy, culture and nucleic acid amplification^3, 5^; however, the low-cost microscopy-based tests used most widely in low-resource settings have limitations, which include low sensitivity^6^. Clearly, new molecular probes for improved, rapid, sensitive and accurate TB diagnostic technologies are needed.

As the architecture of the *Mtb* cell envelope is distinct from other species, comprising an interconnected peptidoglycan-arabinogalactan-mycolic acid (mAGP) core interspersed with a diverse array of mycobacterial specific ‘free’ lipids and glycolipids^7-9^, it provides a unique opportunity to develop *Mtb* pathogen specific reporter probes through utilisation of the specialised mycobacterial cell-wall assembly machinery for the direct incorporation of a detection moiety into the mycobacterial cell-envelope. Indeed, an array of fluorescent-based probes have been developed to monitor *Mtb* metabolism and image cell-envelope components, which have provided new insights into fundamental aspects of *Mtb* biology, including enzyme function, metabolism, cell wall biosynthesis and cell division^10-15^. Since the *Mtb* cell envelope is highly abundant in trehalose-containing glycolipids that include the mycolic acid esters: trehalose monomycolates (TMM) and trehalose dimycolates (TDM), the acyltrehaloses comprising di- and polyacyltrehaloses and sulfoglycolipids, not typically found in other bacterial species^7-9^, recent attention has focused on the use of labelled trehalose derivatives for the visualisation of *Mtb*^10-12, 15-17^. Incorporation of trehalose into mycobacterial cell-envelope glycolipids can proceed *via* two interlinked pathways: either through extracellular synthesis catalysed by the antigen 85 complex (Ag85)^18^ and/or trehalose import^19^ followed by biosynthesis of the TMM glycolipid within the cytoplasm (Fig. 1). The Ag85 complex, which include the secreted proteins AgA, AgB and AgC, is responsible for both the conjugation of extracellular mycolic acids to the arabinogalactan (AG) cell-wall core and the incorporation of a mycolic acid from one TMM into another TMM molecule to generate TDM^18^. Alternatively the ‘free’ trehalose moiety, released concomitantly during Ag85 mediated cell-wall biosynthesis, can be recycled *via* the LpqY-SugABC ATP-binding cassette transporter^19^ and used as a precursor to yield cytoplasmic TMM, which is subsequently shuttled across the membrane by the MmpL3 exporter for insertion into the mycobacterial cell-envelope^20, 21^. Recent studies identified that both pathways can be hijacked to incorporate modified trehalose analogues into mycobacterial cell-envelope trehalose mycolates. The Ag85 complex is known to be particularly promiscuous for a wide panel of extensively modified trehalose substrates^12^ whereas the LpqY-SugABC importer only tolerates minor modifications^11, 16, 22^. Recent structural and functional analysis of the LpqY substrate binding lipoprotein has afforded molecular level insights into the rules governing the precise substrate selectivity for this transporter enabling the design of reporter mimics to hijack this pathway^22^.

**Fig. 1.**
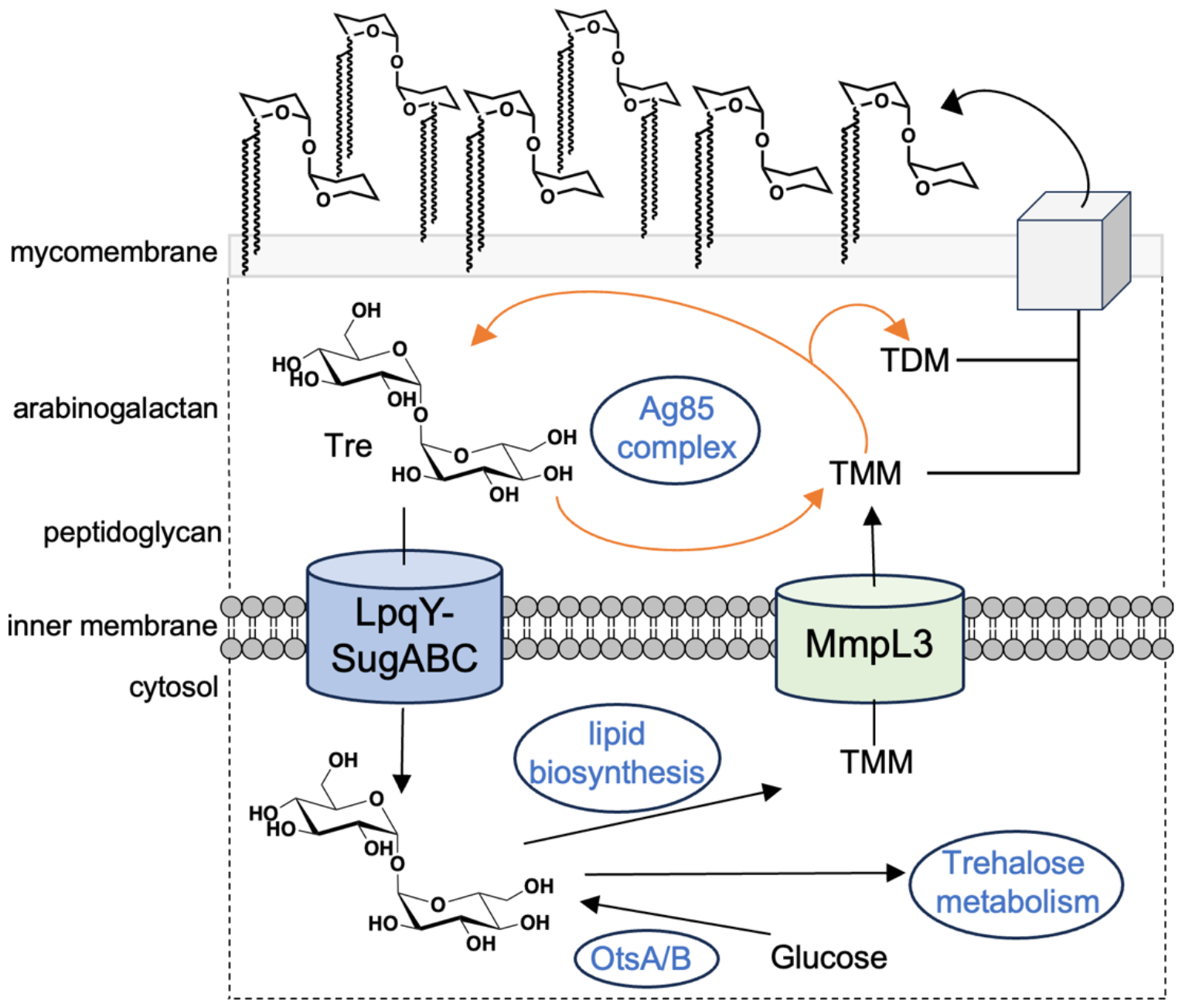
Schematic illustrating the pathways involved in trehalose processing in mycobacteria. TMM: trehalose monomycolate; TDM: trehalose dimycolate

To date, trehalose probes possessing ‘clickable’ azide and alkyne reactive handles have been used extensively for imaging of mycobacteria^11, 12, 16, 23, 24^, however this approach requires extensive wash steps and a secondary, bulky fluorophore label for the direct visualisation of the cells, limiting their use. An ^18^F trehalose-modified radioprobe can be used to label the non-pathogenic model *Mycobacterium smegmatis* organism^17, 25^. However, this direct approach is challenging as fluorine-18 has a short half-life (∼109 minutes) and it is not possible to infer labelling of pathogenic *Mtb* because of the significant differences in nutrient uptake and processing between these diverse mycobacterial species^26-28^. To address these limitations we rationalised we could leverage the *Mtb* LpqY-SugABC transporter to develop and deliver native ^19^F-modified trehalose reporter probes as a direct route to specifically visualise mycobacteria by focused ion beam (FIB) secondary ion mass spectrometry (SIMS). We show here that a discrete panel of unnatural fluorine modified trehalose analogues are indeed tolerated by the *Mtb* LpqY-SugABC transporter and reveal the molecular architecture dictating their specific recognition. Subsequent characterisation demonstrated that this transport promiscuity can be harnessed to facilitate the uptake of fluorinated trehalose probes into live *Mtb* cells, which are subsequently incorporated into extracellular mycobacterial trehalose containing mycolic acids, enabling pathogen specific detection. Taken together, these results highlight the possibility of exploiting native fluorinated-trehalose derivatives to be used not only as tools for unravelling *Mtb* biology but also as new molecular probes for the direct visualisation and detection of *Mtb*.

## Results and Discussion

### Panel of fluorinated trehalose analogues

To explore the potential of fluorinated trehalose for *Mtb* detection and evaluate how selected modifications influence assimilation and cell-labelling, we synthesised a panel of fluorinated trehalose analogues with systematic modifications at either the 2-, 3-, 4-, or 6-position on one glucose subunit with retention of the natural stereochemistry (**2**-**5**). An established chemoenzymatic route^29^ was employed to access 2-fluoro-2-deoxy-trehalose (2F-Tre) (**2**), 3-fluoro-3-deoxy-trehalose (3F-Tre) (**3**) and 6-fluoro-6-deoxy-trehalose (6F-Tre) (**5**) and a modified five-step synthetic route ^17, 30^ (Scheme S1) afforded 4-fluoro-4-deoxy-trehalose (4F-Tre) (**4**), giving the focused panel shown in Fig. 2 (details provided in ESI, Schemes S1, SI Figs S13-S31 and characterisation data).

**Fig. 2.**
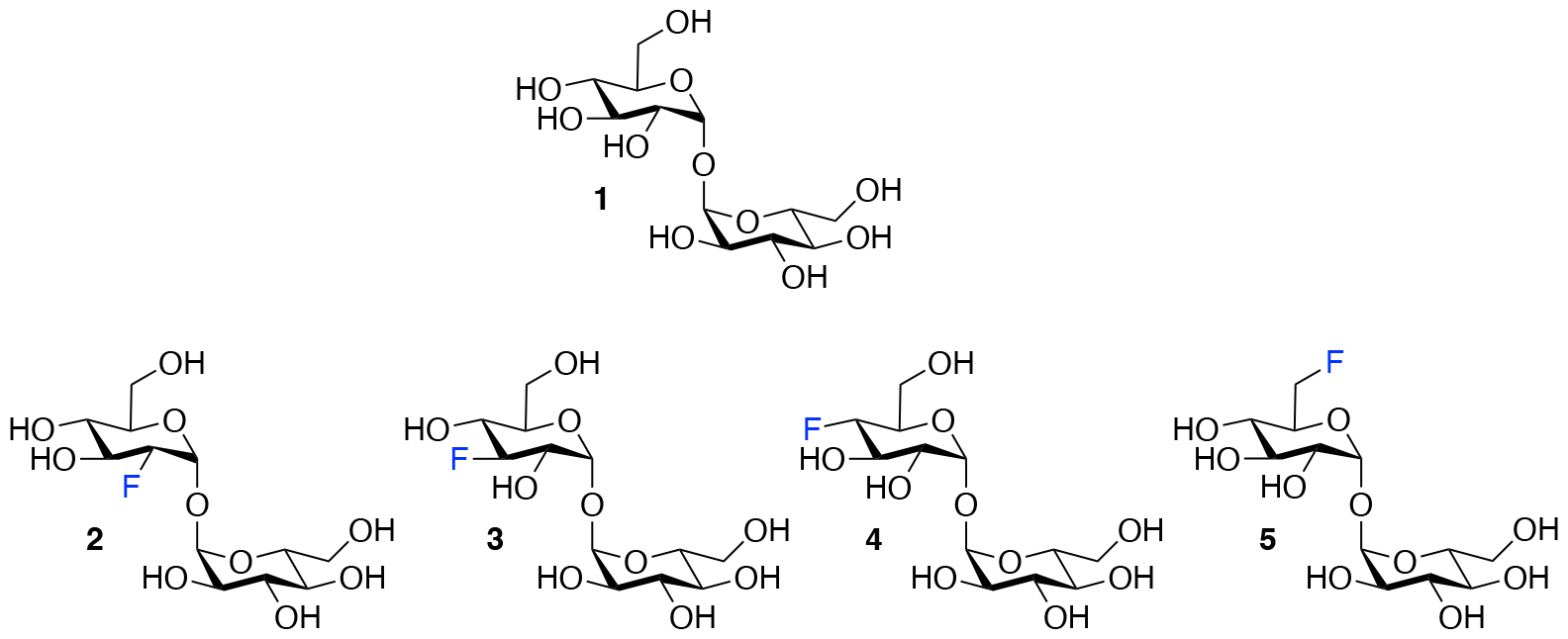
Panel of fluorinated trehalose analogues.

### LpqY substrate specificity for F-Tre analogues

As the mycobacterial specific LpqY-SugABC transporter is solely responsible for the import of trehalose^11, 19, 22^, our first goal was to establish if the introduction, and position, of a fluorine moiety impacts on substrate recognition. We assessed the binding affinity of each fluorinated analogue (**2-5**) to LpqY, the substrate-binding domain that dictates substrate specificity and hence, subsequent transport, by microscale thermophoresis (MST). Highest binding affinities (*K*_*d*_) were observed for 3F-Tre (110.8 ± 7.1 μM) and 6F-Tre (90.3 ± 0.9 μM) with comparable affinities to unmodified trehalose (109.5 ± 3.2 μM) whereas 2F-Tre and 4F-Tre bind ∼8- and ∼16-fold more weakly, with 4F-Tre having affinity in the millimolar range (Table 1, Fig. S1). Altogether, these findings indicate that the position of the fluorine modification is key to LpqY recognition with replacement at the 3- or 6-positions preferred. This is in direct contrast to previous LpqY-azide-trehalose binding analyses (Table 1) ^22^ that identified preferential binding of azide-trehalose modified at the 4- > 6- ≥ 3- > 2-positions, albeit with ∼2-4 weaker affinity than the corresponding fluorinated versions, highlighting the importance of both the substituent type and position in transporter-substrate tolerance and recognition.

**Table 1.** Binding affinity analysis of LpqY to F-Tre analogues.

| Substrate | $K_d$ (μM) | Reference |
| --- | --- | --- |
| trehalose | $109.5 \pm 3.2$ | This study |
| 2-fluoro-2-deoxy-trehalose (2F-Tre) | $770.4 \pm 23.2$ | This study |
| 2-azido-2-deoxy-trehalose (2Az-Tre) | $1915.6 \pm 7.7$ | <sup>22</sup> |
| 3-fluoro-3-deoxy-trehalose (3F-Tre) | $110.8 \pm 7.1$ | This study |
| 3-azido-3-deoxy-trehalose (3Az-Tre) | $474.9 \pm 17.2$ | <sup>22</sup> |
| 4-fluoro-4-deoxy-trehalose (4F-Tre) | $1483.7 \pm 75.6$ | This study |
| 4-azido-4-deoxy-trehalose (4Az-Tre) | $246.6 \pm 7.1$ | <sup>22</sup> |
| 6-fluoro-6-deoxy-trehalose (6F-Tre) | $90.3 \pm 0.9$ | This study |
| 6-azido-6-deoxy-trehalose (6Az-Tre) | $442.7 \pm 5.3$ | <sup>22</sup> |

Mean ± standard error mean is from at least three independent experiments. The substrate structures are shown in Fig. 2. The binding affinity data is shown in Fig S1.

### Molecular basis of F-Tre analogue recognition

Having established that LpqY has the capacity to recognise fluorinated trehalose analogues we sought to determine the molecular rules of recognition in solution. Saturation transfer difference (STD) NMR confirmed binding for each fluorinated trehalose analogue, verifying that switching a hydroxyl group to fluorine at all possible positions is not detrimental for recognition by the mycobacterial trehalose transporter (Fig. 3). The STD NMR intensities differed depending on the temperature of the experiment (Fig. S2-3 and Tables S1-4). More intense signal intensities were observed at low temperature (5 °C) for 2F-Tre and 4F-Tre, whereas a higher temperature of 30 °C was required for STD NMR observation of 3F-Tre and 6F-Tre, especially in the case of the 3-fluoro derivative where the particularly low signal-to-noise ratio meant no STD NMR analysis was possible at 5 °C. These results indicate a much stronger affinity for 3F-Tre and 6F-Tre, in comparison with the 2- and 4-fluoro derivatives, consistent with our binding affinity data (Table 1). STD binding epitope maps are unique for each F-Tre analogue (Fig. 3) and indicate that protons from both the unmodified and modified carbohydrate rings make distinct close contacts to LpqY relevant for recognition, with a significant overall decrease in the relative STD intensities at the modified fluorinated glucose ring (Fig. 3).

**Fig. 3.**
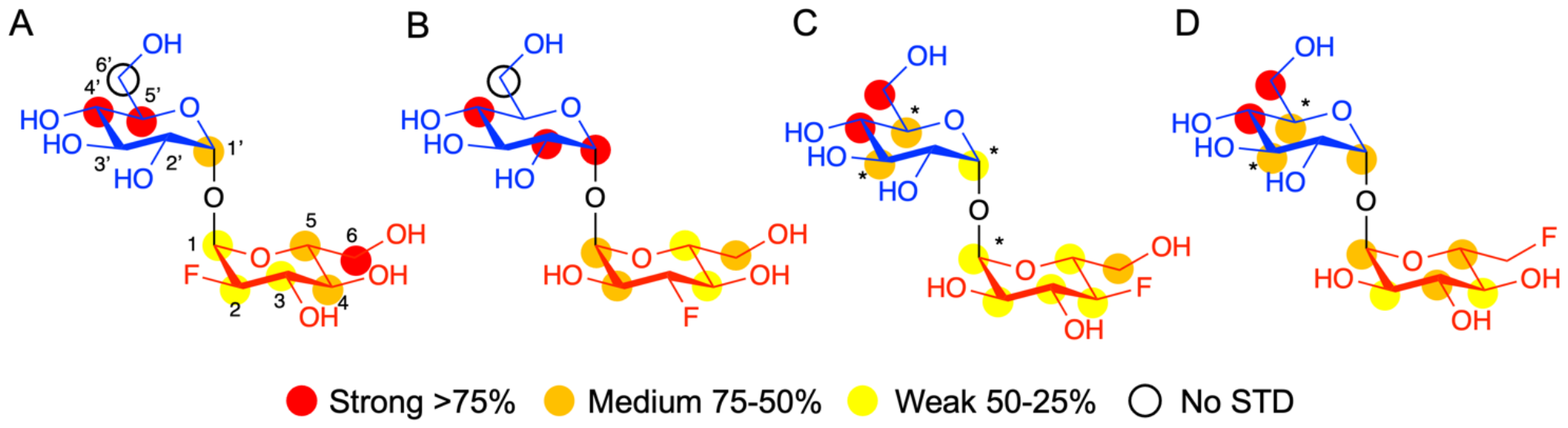
STD NMR binding epitope mapping of F-Tre analogues (2-5) to LpqY. Binding epitope map of (A) 2F-Tre, (B) 3F-Tre, (C) 4F-Tre and (D) 6F-Tre. The coloured spheres represent normalized STD NMR intensities. Protein saturation was achieved by irradiation at 0.53 ppm for (A) and (C), whereas irradiation was at 0.84 ppm for (B) and (D). The coloured spheres represent normalized STD NMR intensities. STD responses are only indicated for protons that could accurately be measured. Asterisks indicate averaged STD values due to overlapping. Glc-1 *(blue)*, Glc-2 *(red)*, STD NMR, saturation transfer difference NMR.

Comparative DEEP-STD NMR analyses^31^, which assesses the type of protein residue that makes contacts with the fluorinated trehalose analogues (Fig. S4), in combination with molecular dynamics simulations over 100 ns (SI Fig S5) revealed the relative orientations of the four ligands in the LpqY binding pocket (Fig. 4). Each of the F-Tre analogues bind in similar relative orientations and sit in an analogous position to trehalose (PDB 7APE)^22^ (Fig S6). The fluorinated-modified glucose ring is located towards the entrance of the substrate binding cavity, similar to the orientation we observed previously for 6Az-Tre ^22^, indicating that when the second glucose ring is modified it is preferentially accommodated within an expanded binding pocket at the channel entrance rather than buried at the base of the binding cleft (Fig. 4). Because the sidechains of Asn25, Glu26, Gln59 and Asn134 make hydrogen bond contacts with the modified glucose ring it is clear this interaction network plays an important role in controlling fluoro-modified trehalose specificity. Indeed, the significantly reduced affinity of 4F-Tre could be attributed to weakened interactions through the loss of a hydrogen-bond partner with Glu26 when a fluorine group is substituted at the 4-position.

**Fig. 4.**
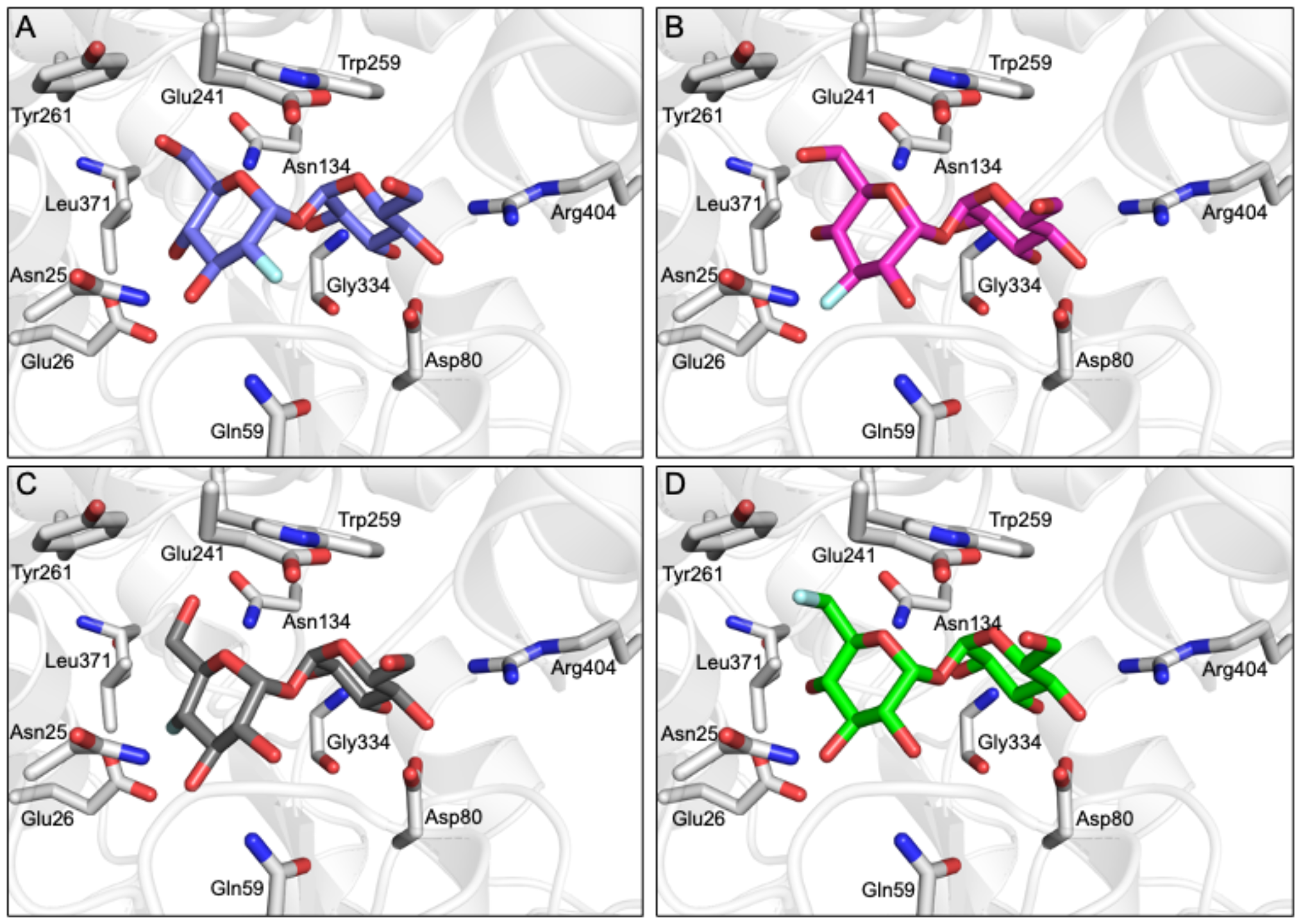
F-Tre binding to LpqY. Model of the mode of binding of F-Tre analogues based on experimentally derived DEEP-STD NMR derived interactions and molecular dynamic simulations. (A) 2F-Tre (blue carbon atoms), (B) 3F-Tre (magenta carbon atoms), (C) 4F-Tre (dark grey carbon atoms), and (D) 6F-Tre (green carbon atoms). Cartoon representation of LpqY in grey (PDB 7APE), trehalose and binding site residues are represented as sticks (oxygen: red; fluorine: pale blue).

### *Mtb* imports 2F-, 3F- and 6F-Tre analogues *via* the LpqY-SugABC transporter

Having confirmed that the fluorinated trehalose analogues bind to LpqY, and since not all substrates recognised by periplasmic binding domains are translocated by the cognate ABC transporter, we sought to evaluate whether these derivatives are taken up by live *Mtb* cells by ion chromatography. Treatment of *Mtb* cells to each fluorinated analogue led to a significant dose-dependent increase in the accumulation of 2F-Tre, 3F-Tre and 6F-Tre in *Mtb* cells, with the assimilation of micromolar levels even after exposure to the lowest concentration of the probe (25 μM) (Fig. 5). No uptake of 4F-Tre was detected, a finding consistent with its poor binding affinity to LpqY. The uptake pattern matches the entry of these analogues into non-pathogenic *M. smegmatis* cells, suggesting overlapping fluorine-modified trehalose analogue recognition profiles between these bacilli^17^. These findings, together with the determined F-Tre analogue affinities for LpqY, emphasise the importance of the LpqY substrate binding domain in controlling which substrates are translocated.

**Fig. 5.**
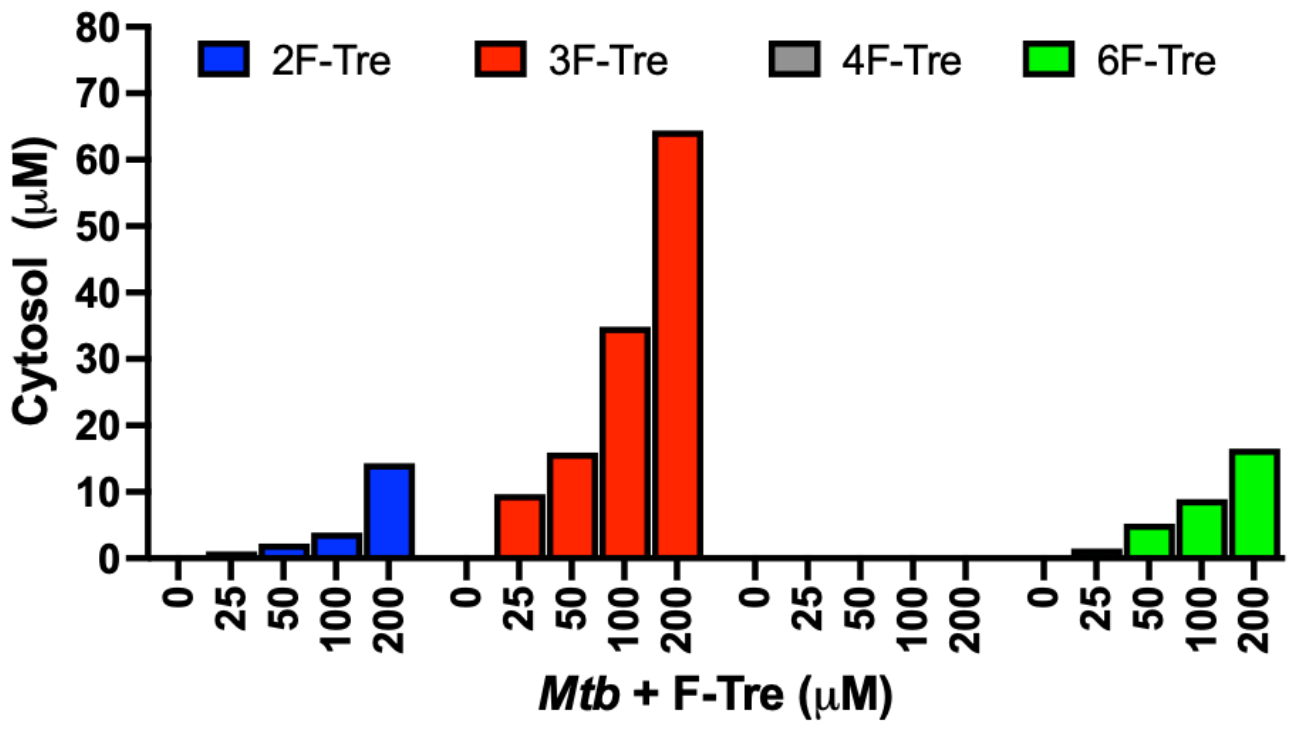
Ion chromatography analysis of F-Tre labelled *Mycobacterium tuberculosis*. *Mtb* was cultured in the presence of F-Tre analogues **2**-**5** (0 – 200 μM) and the cytosolic extracts analysed by high performance anion exchange chromatography with pulsed amperometric detection (HPAEC-PAD), and F-Tre concentration quantified. The ion-chromatography traces are shown in Fig S7.

As *Mtb* efficiently imports the 2-, 3- and 6-fluorinated trehalose derivatives we then asked how rapidly each analogue is accumulated. We evaluated uptake in cytosolic extracts and calculated accumulation rates of 0.80 μM.h^-1^ (2F-Tre) 0.94 μM.h^-1^ (3F-Tre) and 0.82 μM.h^-1^ (6F-Tre). Rapid uptake of each F-Tre derivative was observed at 30 mins and at 8 hours ∼ 20% increased accumulation of 3F-Tre was detected compared to 2F- and 6F-Tre (Fig 6, Fig S8, Table S5). No saturation of uptake was observed for each F-Tre analogue **2, 3** and **5**, indicating that an equilibrium between trehalose import and export has not yet been reached at the 8-hour timepoint, which may be consistent with extremely slow growth rate of the bacillus. Altogether, these observations establish that 2F-, 3F- and 6F-trehalose are rapidly transported, and accumulate, in live *Mtb* cells.

**Fig. 6.**
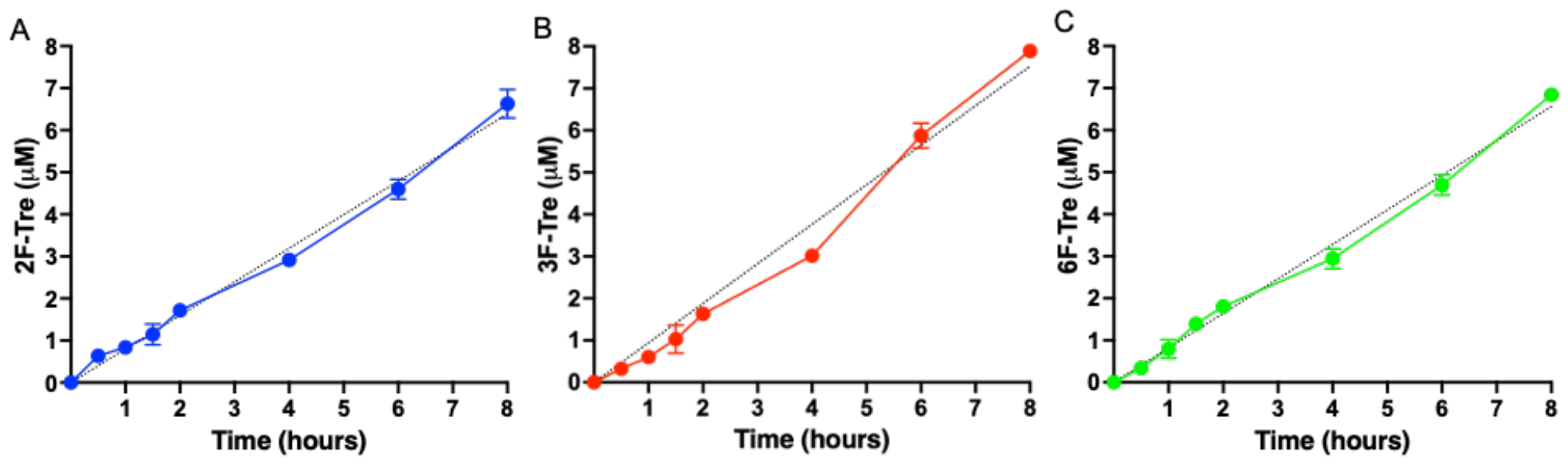
Time dependent uptake of F-Tre analogues in *Mycobacterium tuberculosis*. *Mtb* was cultured in the presence of (A) 2F-Tre, (B) 3F-Tre and (C) 6F-Tre (100 μM) and the cytosolic extracts sampled at the time points indicated and analysed by high performance anion exchange chromatography with pulsed amperometric detection (HPAEC-PAD). Error bars denote the standard deviation from duplicate experiments. Simple linear regression fit is shown with a black dotted line (A) y=80x; (B) y= 0.94x; (C) y=0.82x. Representative ion-chromatography traces are shown in Fig S8, and the concentration values are in Table S5.

We then asked whether the F-Tre analogues are imported *via* LpqY-SugABC transporter or *via* an alternative route and evaluated uptake in a *Mycobacterium bovis* BCG strain lacking the trehalose transporter *ΔlpqY-sugABC)* ^11^, as a model for *Mtb*. In agreement with our *Mtb* data 2F-, 3F- and 6F-Tre accumulate in wild-type *M. bovis* BCG cytosolic extracts and not 4F-Tre (Fig. 7). Notably, uptake of each F-Tre derivative was completely abolished in the *ΔlpqY-sugABC* mutant and transport restored in the corresponding complemented strain. Taken together, as the mutant strain is unable to utilise F-Tre, these data clearly show that uptake of fluorinated trehalose analogues occurs solely *via* the trehalose LpqY-SugABC import system.

**Fig. 7.**
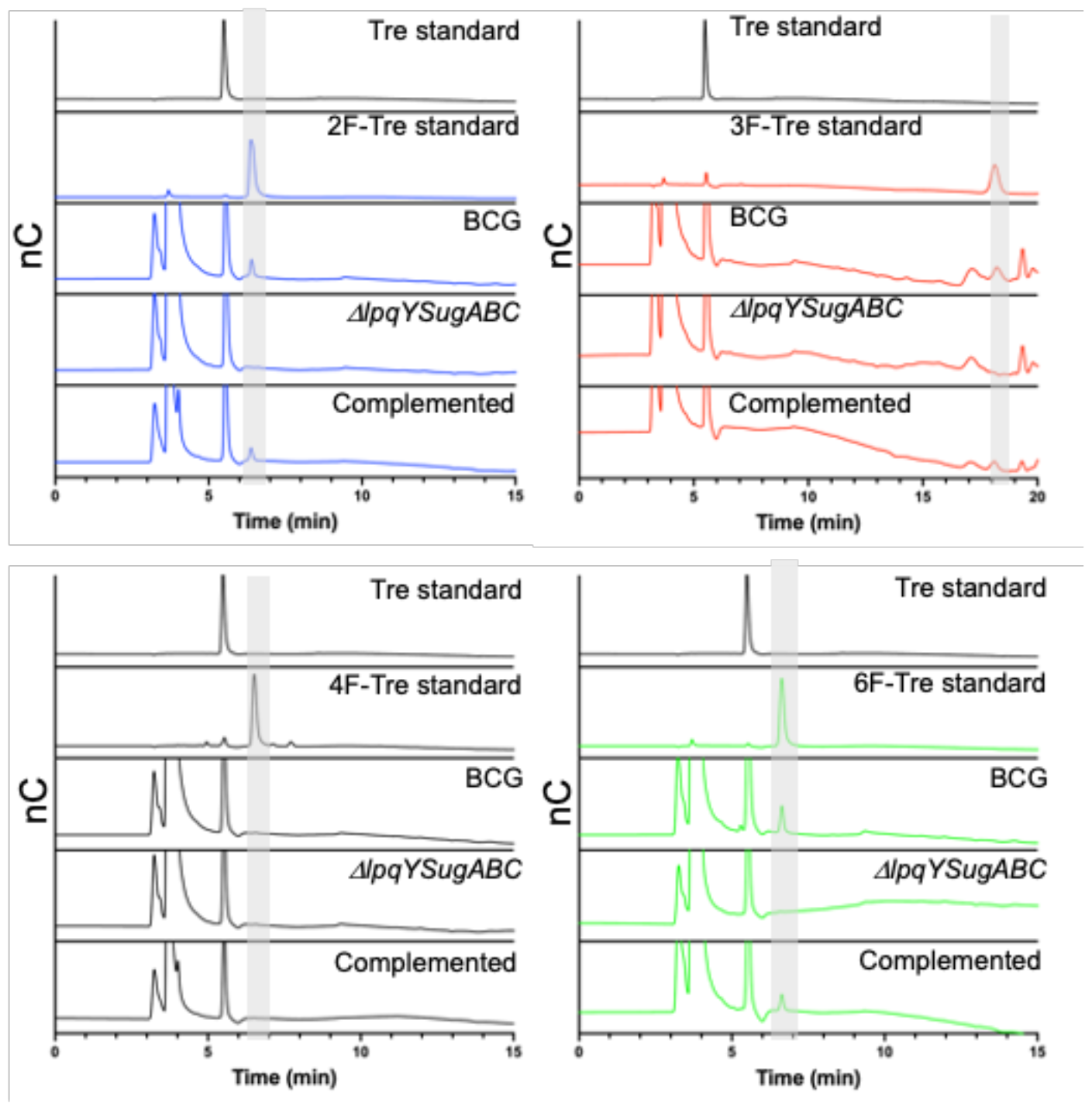
Comparison of F-Tre uptake in *Mycobacterium tuberculosis, Mycobacterium bovis* BCG and LpqY-SugABC mutant strains. Wild-type *Mycobacterium tuberculosis, Mycobacterium bovis* BCG and the *ΔlpqY-sugABC Mycobacterium bovis* BCG mutant and complemented strain *Mycobacterium bovis* BCG *ΔlpqY-sugABC::lpqY-SugABC* were cultured in the presence of F-Tre analogues **2, 3, 4** and **5** (100 μM) and the cytosolic extracts analysed by high performance anion exchange chromatography with pulsed amperometric detection (HPAEC-PAD).

### *Mtb* incorporates 2F-, 3F- and 6F-Tre analogues into trehalose-containing glycolipids

To determine the fate of F-Tre analogues following import across the mycobacterial cell-envelope *Mtb* and *M. bovis* BCG were exposed to each F-Tre analogue (**2**-**5**), alongside trehalose and untreated cells, and the extracted lipids evaluated by thin layer chromatography (TLC) and ion chromatography (Fig. 8 and Fig. S9). Treatment of *Mtb* and *M. bovis* BCG with 2F-, 3F- and 6F-Tre afforded new spots in the lipid extracts that were not present in the control samples (Fig. 8). No new spots were observed with 4F-Tre addition, consistent with its lack of uptake. Two additional lower polarity bands than TMM are present for the 2F- and 3-F samples and one for 6F-Tre (Fig. 8), indicating their incorporation into TMM (Fig. 8).This labelling pattern is consistent with two possible fluorinated-TMM regiochemistries for 2F- and 3F-Tre, where the fluorine is either on the same glucose moiety as the mycolic acid or the other subunit (Fig. 8). Since the TMM 6-position is blocked through mycolic attachment, only one 6-F trehalose isomer is possible, as observed (Fig. 8). The presence of one additional lower polarity TDM band for 2F- and 3F-Tre suggests these analogues are also incorporated into the TDMs. As expected, when mycobacteria are labelled with 6F-Tre no additional band in the TDM region is observed, since both 6-positions are inaccessible due to the attachment of mycolic acids on both glucose rings preventing the production of 6-fluorinated TDM.

**Fig. 8.**
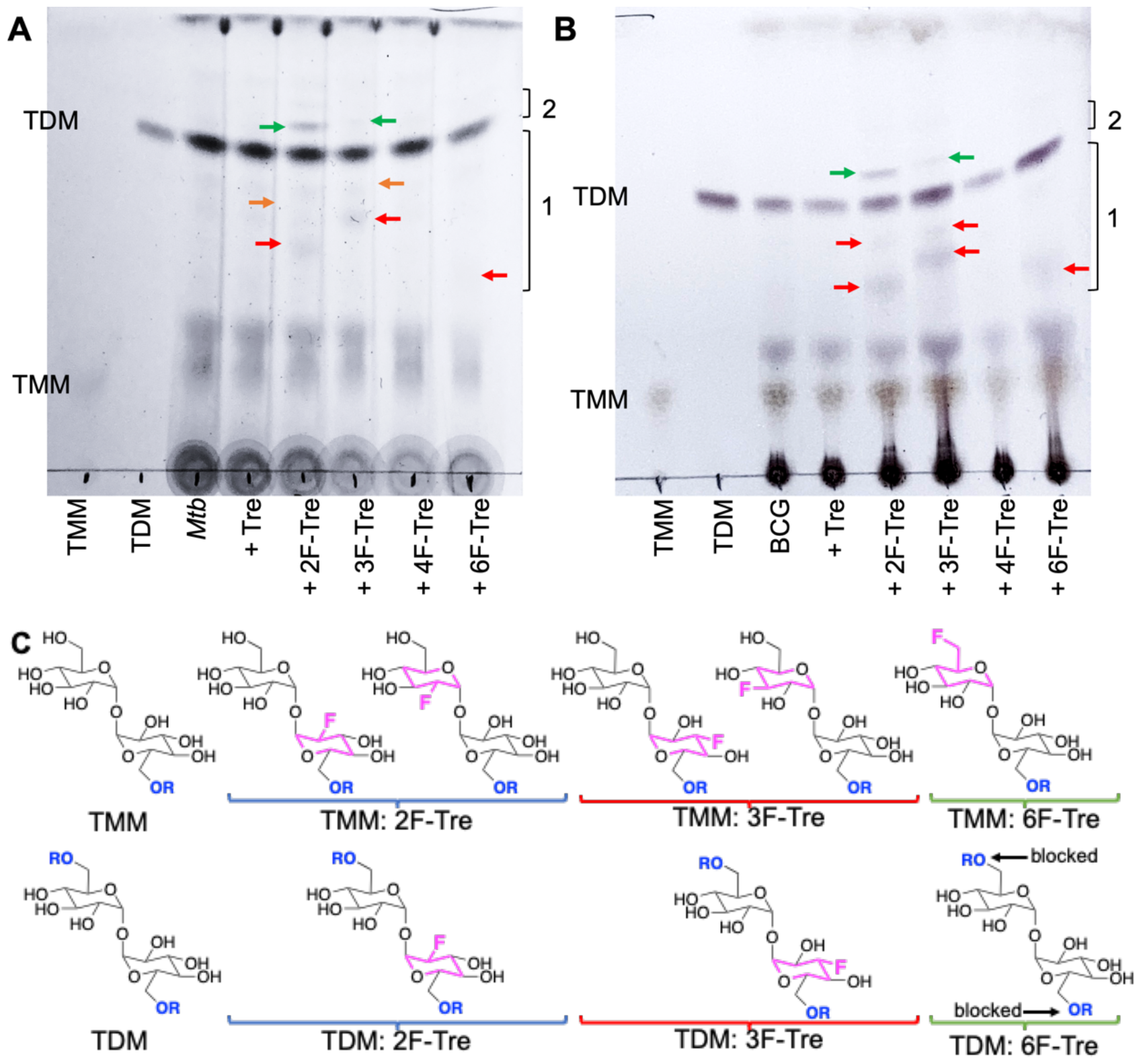
Thin layer chromatography analysis of *Mtb* and *M. bovis* BCG lipids after labelling with F-Tre analogues. A) *Mtb* extracted apolar lipids. B) *M. bovis* BCG extracted apolar lipids. C) Structures illustrating potential F-Tre TMM and TDM modifications. Red arrows indicate new spots with intermediate polarity compared to TMM and TDM standards in region 1; green arrows indicated new spots which are less polar than TDM in region 2; orange arrows indicate new spots with intermediate polarity compared to TMM and TDM standards in region 1 that are visible directly on the TLC plate but not as clearly visible on the image. The image has been contrast adjusted to improve the visualization of spots. TMM: trehalose monomycolate standard, TDM: trehalose dimycolate standard.

Altogether, these experiments demonstrate that, unless the 4-position is modified, mycobacteria have the capacity to take up fluorinated trehalose analogues and incorporate these modified sugars into cell-surface trehalose mycolates. It is notable that 4F-Tre is not found in any trehalose glycolipid in either *Mtb* or *M. bovis* BCG as this differs from the situation found for the analogous azido-trehalose derivatives which are all incorporated into mycobacterial glycolipids, albeit *via* different routes, depending on the mycobacterial species.^11, 17^ For example, in *M. bovis* BCG, 3- and 4-azido trehalose are not recycled *via* LpqY-SugABC but, instead, incorporated into TMM and TDM by the extracellular Ag85 complex^11^. Yet this does not occur for any F-Tre analogue. Thus, these findings reveal distinct mechanistic pathways for trehalose analogue recognition and use, which depend on both the type and position of modification and the mycobacterial species, highlighting an evolutionary divergence of trehalose transport across mycobacterial species.

### *Mtb* does not readily metabolise 2F-, 3F- and 6F-Tre analogues

Next, to determine if the fluorinated-trehalose derivatives are processed further we analysed the same cytosolic extracts of *Mtb* exposed to **2, 3**, and **5** for the presence of fluorinated-glucose break-down products: 2-fluoro-2-deoxy-glucose (2F-Glc), 3-fluoro-3-deoxy-glucose (3F-Glc) and 6-fluoro-6-deoxy-glucose (6F-Glc) respectively (Fig. S10). No cytosolic 3F- or 6F-glucose were found, indicating that the cleavage of 3F- and 6F-Tre does not occur. This suggests that intact 3F- and 6F-Tre are predominantly processed by pathways that ultimately incorporate these derivatives into cell-surface trehalose glycolipids. In contrast, *Mtb* labelled with 2F-Tre revealed the presence of low levels of 2F-Glc, indicating some degradation occurs, likely catalysed by the *Mtb* trehalase ^32^. It is possible that the 2F-Glc metabolite is then diverted into alternative glucose utilisation pathways and not solely incorporated into cell-surface trehalose opening up potential off-target labelling routes.

### F-Tre analogues specifically label mycobacteria

It is important that a diagnostic probe is specific for mycobacteria, therefore we evaluated fluorinated-trehalose assimilation in a panel of bacteria including the ESKAPE pathogens *Enterobacter cloacae, Staphylococcus aureus, Klebsiella pneumoniae, Acinetobacter baumannii, Pseudomonas aeruginosa* and *Enterococcus faecalis* as well as the representative Gram-negative *Escherichia coli* and Gram-positive *Bacillus subtilis* species. *Acinetobacter baumannii, Klebsiella pneumoniae, Pseudomonas aeruginosa* and *Staphylococcus aureus* are particularly relevant as lung pathogens. To test whether trehalose is important for these organisms we assessed natural cytosolic trehalose levels in these bacteria. Detectable levels of intracellular trehalose were only observed for *A. baumannii* and *E. cloacae*, which were ∼6-fold and ∼11-fold lower compared to *Mtb* (Fig. 9A) that contains especially high levels of this disaccharide ^33^. The addition of exogenous trehalose did not lead to an increase in the intracellular trehalose level in any organism (Fig. 9B), suggesting these bacterial species lack the homologous import machinery to assimilate trehalose and instead likely synthesise trehalose *de novo* from glucose-6-phosphate and UDP-glucose *via* an OtsA/B system.^34^

**Fig. 9.**
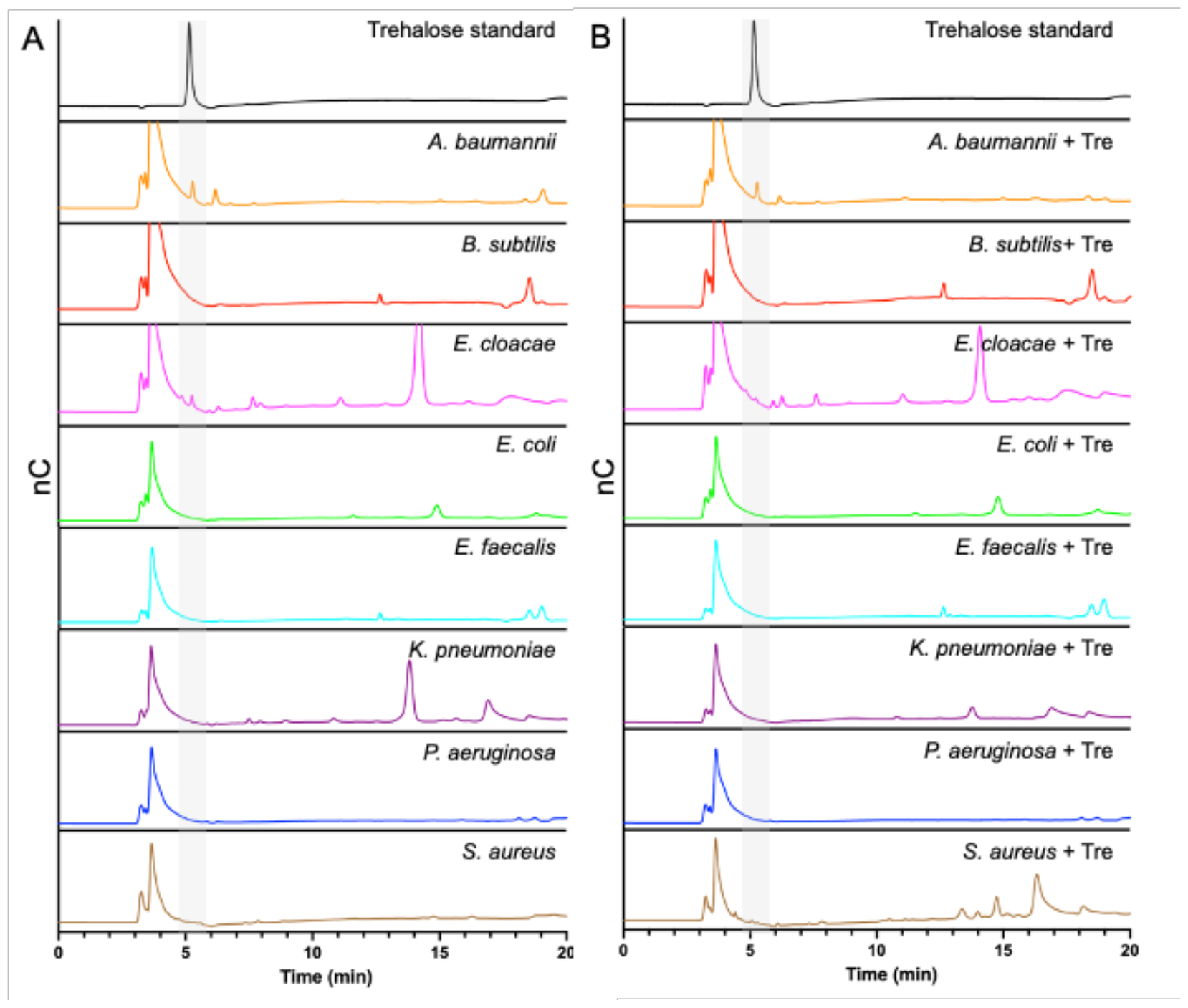
Cytosolic extracts analysis of the ESKAPE pathogens, *E. coli* and *B. subtilis*. The bacterial strains were cultured to mid-log phase in the (A) absence and (B) presence of trehalose (100 μM) and the cytosolic extracts were analysed by HPAEC-PAD.

Despite the lack of trehalose uptake, we investigated whether the ESKAPE pathogens, *E. coli* and *B. subtilis* could take up the fluorinated-trehalose derivatives. Each strain was cultured with **2**-**5** (100 μM) for four-doubling times and the cytosolic extracts analysed by HPAEC. Under these conditions, no uptake of any F-Tre analogue was measurable (Fig. S11). Taken together, these findings have established that the assimilation of fluorinated-trehalose derivatives is highly specific to mycobacteria and rules out uptake in non-mycobacterial species lacking homologous trehalose importers.

### Imaging F-Tre labelled *Mtb* by secondary ion mass spectrometry

As fluorine-modified trehalose analogues are incorporated into cell-surface trehalose-containing *Mtb* glycolipids, we next examined the ability to visualise and image F-Tre labelled *Mtb* cells using focused ion beam (FIB) secondary ion mass spectrometry (SIMS), focused on ^19^F^-^ ions. Elemental scanning for negative ions in the mass range 1-100 detected ^19^F^-^ fluorine ions in all *Mtb* samples exposed to F-Tre and only trace levels in non-treated cells (Fig. S12). We then mapped the distribution of fluorine in the samples, first imaging the *Mtb* cells by scanning electron microscopy (SEM) to show the morphology of the cells and then analysis of the same section by FIB-SIMS. The intensity of the SIMS ^19^F^-^ signal correlated with the scanning electron micrograph and showed a clear pattern of accumulation and localisation of F-Tre in treated *Mtb* cells (Fig. 10). In contrast only background ^19^F^-^ levels were observed in untreated cells. Crucially, our findings establish that F-Tre labelling of live *Mtb* cells affords a prominent ^19^F^-^ SIMS signal and low background demonstrating the feasibility of this approach for the direct detection and imaging of mycobacteria. Whilst robust labelling is observed for each probe, it is conceivable that the incorporation of multiple fluorine atoms on the same glucose ring that combines modifications at either the 2- 3-, or 6-positions could enhance the signal intensity further.

**Fig. 10.**
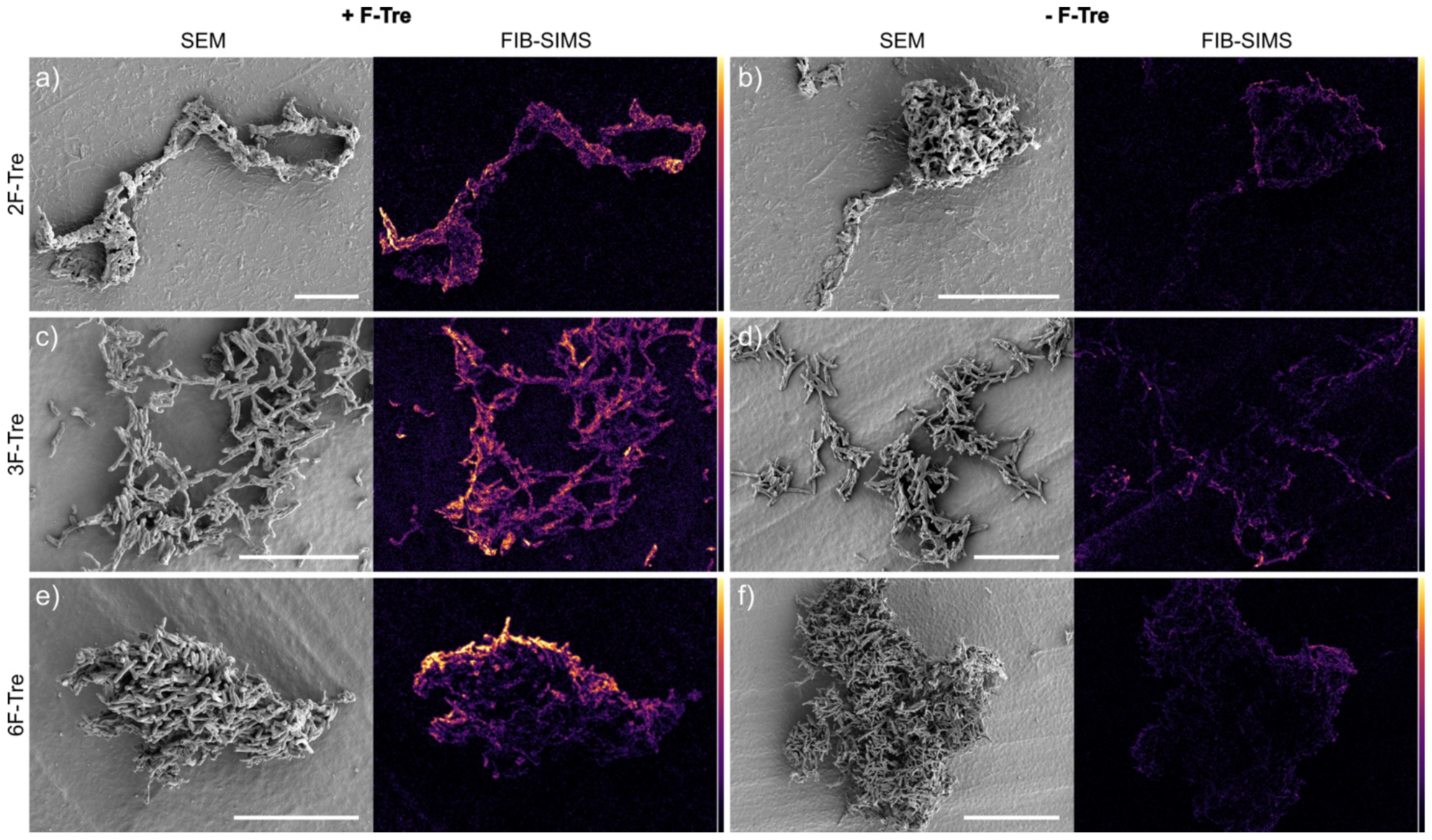
F-Tre labelled *Mtb*. Scanning electron microscope (SEM) images and focused ion beam-secondary ion mass spectrometry (FIB-SIMS) maps of *Mtb* cells treated with (A) 2F-Tre, (C) 3F-Tre (E) 6F-Tre and untreated control cells (B), (D) and (F). Colour scale represents ^19^F^-^ SIMS counts, black = lowest intensity, yellow = highest intensity. White scale bar 10 μm (a), (c), (d) and (e), 20 μm (b) and (d).

## Conclusions

One of the major hurdles in controlling the TB pandemic is the need for improved TB diagnosis. One route to achieve this is to transition from sputum microscopy to molecular testing. Yet molecular probes for the direct detection of TB are lacking. To address this deficiency, we developed native fluorinated trehalose probes for imaging by FIB-SIMS, providing unique opportunities to specifically visualise and detect the *Mtb* pathogen. In conclusion, here we have clearly shown that we can hijack the promiscuity of the mycobacterial specific LpqY-SugABC trehalose transport machinery to incorporate fluorinated reporters into mycobacterial specific cell-surface glycolipids, combined with FIB-SIMS imaging to resolve ^19^F^-^ localisation and directly image *Mtb* cells. Moreover, our findings demonstrate that F-Tre probes allow for pathogen specific detection of *Mtb* with high sensitivity and specificity over other bacterial species. Expanding on this, our studies reveal a discernible dependence on the position of fluorine atom modification for efficient *Mtb* labelling. Specifically, we have uncovered the molecular framework that dictates probe recognition and import allowing us to explain F-Tre selectivity. We have shown that a single fluorine modification at the 2-, 3- and 6-positions of one glucose ring is optimal since substitution at the 4-position disrupts a critical hydrogen bond interaction that significantly weakens interactions with LpqY, impeding 4F-Tre import. The structural insights into F-Tre recognition and uptake analysis provided by this study, combined with FIB-SIMS imaging of *Mtb* cells provides a firm foundation that open opportunities for the rational development of high-contrast molecular probes for *Mtb* pathogen identification and provides an important step towards expanding the existing toolbox of molecular reporters for TB diagnosis.

## Conflicts of interest

There are no conflicts to declare.

## Data availability

The datasets supporting this article have been uploaded as part of the ESI.

## Acknowledgements

This work was supported by a Sir Henry Dale Fellowship to EF jointly funded by the Wellcome Trust and Royal Society (104193/Z/14/Z and 104193/Z/14/B) and a research grant from the Leverhulme Trust (RPG-2019-087. We thank Dr Ben Swarts (Central Michigan University, USA) for providing the TreT construct, Professor Rainer Kalscheuer (HHU Dusseldorf, Germany) for kindly providing the *M. bovis* BCG mutant strains, Dr John Moat for providing the ESKAPE pathogens used in this study, Dr Saskia Bakker for help with sample preparation for FIB-SIMS and Dr James Harrison for critical reading of this manuscript. We acknowledge equipment access, training, and support made available by the Research Technology Facility (managed by Dr Sarah Bennett) of the Warwick Integrative Synthetic Biology centre (WISB), which received funding from EPSRC and BBSRC (BB/M017982/1). J. A. acknowledges support from the Universidad de Sevilla (Acciones Especiales del VI Plan Propio de Investigación y Transferencia).

## Notes

### Competing Interest Statement

The authors have declared no competing interest.

