## Supporting information for "Fluorinated trehalose analogues for cell surface engineering and imaging of *Mycobacterium tuberculosis*"

*To whom correspondence should be addressed:

### Table of Contents

**Supplementary Results** 4
Scheme S1. Synthetic route for 4-fluoro-4-deoxy-trehalose 4
Figure S1: Binding affinities for LpqY 5
Fig. S2. STD NMR study of the binding of *Mtr* LpqY with 2F-Tre, 4F-Tre and 6F-Tre with protein irradiation in the aliphatics spectral region 6
Fig. S3. STD NMR *for Mtr* LpqY with 2F-Tre, 4F-Tre and 6F-Tre with protein irradiation in the aromatics spectral region 7
Table S1. STD NMR data for the binding of *Mtr* LpqY with 2-fluoro-trehalose 8
Table S2. STD NMR data for the binding of *Mtr* LpqY with 3-fluoro-trehalose at a single saturation time 9
Table S3. STD NMR data for the binding of *Mtr* LpqY with 4-fluoro-trehalose 10
Table S4. STD NMR data for the binding of *Mtr* LpqY with 6-fluoro-trehalose 11
Fig. S4. Differential Epitope Mapping by STD NMR of *Mtr* LpqY with 2-fluoro-trehalose, 3-fluoro-trehalose, 4-fluoro-trehalose and 6-fluoro-trehalose 12
Fig S5. Molecular dynamic simulations *Mtr* LpqY with 2-fluoro-trehalose, 3-fluoro-trehalose, 4-fluoro-trehalose and 6-fluoro-trehalose 13
Fig S6. Comparison of the binding orientations of the F-Tre analogues 14
Fig. S7. Ion chromatography traces of F-Tre labelled *Mycobacterium tuberculosis* 15
Table S5. *Mtb* cytosolic concentrations of F-Tre analogue uptake 16
Fig. S8. Ion chromatography traces of F-Tre labelled *Mycobacterium Mtb* 17
Fig. S9. Ion chromatography traces of hydrolysed lipid extracts F-Tre labelled *Mycobacterium Mtb* 18
Fig. S10. F-Glc metabolite analysis of *Mtb* labelled with F-Tre analogues 19
Fig. S11. F-Tre analogue uptake analysis in the ESKAPE pathogens, *Eschericia coli* and *Bacillus subtilis* 20
Fig. S12. SIMS mass spectra of *Mtb* cells treated with 2F-Trehl, 3F-Trehl, 6F-Trehl and non treated control cells, from m/z 1-50 21
**Experimental** 22
General Information and Procedures 22
$^{1}$H NMR, $^{13}$C NMR and MS data 22
Bacterial strains, cell lines, culture conditions and chemicals 22
Expression and purification of TreT 23
Chemoenzymatic synthesis of fluorinated trehalose derivatives 23
2-fluoro-2-deoxy-trehalose (**2**) 23
3-fluoro-3-deoxy-trehalose (**3**) 24
6-fluoro-6-deoxy-trehalose (**5**) 24
Chemical synthesis of 4-fluoro-4-deoxy-trehalose 24
2,3,6,2′,3′,4′,6’,-hepta-*O*-benzoyl-α,α’-D-trehalose (**6**) 24
2,3,6,-tri-*O*-benzoyl-α-D-galactopyranosyl-(1→1)-2′,3′,4′,6′,-tetra-*O*-benzoyl-α-D-glucopyranoside (**8**) 25
4-fluoro-2,3,6,-tri-*O*-benzoyl-α-D-glucopyranosyl-(1→1)-2′,3′,4′,6′,-tetra-*O*-benzoyl-α-D-glucopyranoside (**9**) 25
4-fluoro-4-deoxy-trehalose (**4**) 26
Production and purification of *Mtr* LpqY 26
Microscale Thermophoresis 27
STD-NMR 27
Molecular dynamics 28
Input preparation and equilibration 28
MD simulation 28
Analysis of F-Tre uptake at a single-time point 29
Table S6: High performance anion exchange chromatography KOH elution gradient for F-Tre analysis 29
Time dependent F-Tre uptake 30
Analysis of F-Glc metabolites 30

Table S7: High performance anion exchange chromatography KOH elution gradient for F-Glc analysis 30
Lipid extraction and analysis 31
Preparation of samples for focussed ion beam (FIB) secondary ion mass spectrometry (SIMs) 31
Scanning electron microscopy (SEM) and FIB-SIMS 32
**NMR Spectra** 33
Figure S13. $^{1}$H NMR 2-fluoro-2-deoxy-trehalose (**2**) 33
Figure S14. $^{13}$C NMR 2-fluoro-2-deoxy-trehalose (**2**) 33
Figure S15. $^{19}$F NMR 2-fluoro-2-deoxy-trehalose (**2**) 34
Figure S16. $^{1}$H NMR 3-fluoro-3-deoxy-trehalose (**3**) 34
Figure S17. $^{13}$C NMR 3-fluoro-3-deoxy-trehalose (**3**) 35
Figure S18. $^{19}$F NMR 3-fluoro-3-deoxy-trehalose (**3**) 35
Figure S19. $^{1}$H NMR of 4-fluoro-4-deoxy-trehalose (**4**) 36
Figure S20. $^{13}$C NMR of 4-fluoro-4-deoxy-trehalose (**4**) 36
Figure S21. $^{19}$F NMR of 4-fluoro-4-deoxy-trehalose (**4**) 37
Figure S22. $^{1}$H NMR of 6-fluoro-6-deoxy-trehalose (**5**) 37
Figure S23. $^{13}$C NMR of 6-fluoro-6-deoxy-trehalose (**5**) 38
Figure S24. $^{19}$F NMR of 6-fluoro-6-deoxy-trehalose (**5**) 38
Figure S25. $^{1}$H NMR of 2,3,6,2′,3′,4′,6’,-hepta-*O*-benzoyl-α,α’-D-trehalose (**6**) 39
Figure S26. $^{13}$C NMR of 2,3,6,2′,3′,4′,6’,-hepta-*O*-benzoyl-α,α’-D-trehalose (**6**) 39
Figure S27. $^{1}$H NMR of 2,3,6,-tri-*O*-benzoyl-α-D-galactopyranosyl-(1→1)-2′,3′,4′,6′,-tetra-*O*-benzoyl-α-D-glucopyranoside (**8**) 40
Figure S28. $^{13}$C NMR of 2,3,6,-tri-*O*-benzoyl-α-D-galactopyranosyl-(1→1)-2′,3′,4′,6′,-tetra-*O*-benzoyl-α-D-glucopyranoside (**8**) 40
Figure S29. $^{1}$H NMR of 4-fluoro-2,3,6,-tri-*O*-benzoyl-α-D-galactopyranosyl-(1→1)-2′,3′,4′,6′,-tetra-*O*-benzoyl-α-D-glucopyranoside (**9**) 41
Figure S30. $^{13}$C NMR of 4-fluoro-2,3,6,-tri-*O*-benzoyl-α-D-galactopyranosyl-(1→1)-2′,3′,4′,6′,-tetra-*O*-benzoyl-α-D-glucopyranoside (**9**) 41
Figure S31. $^{19}$F NMR of 4-fluoro-2,3,6,-tri-*O*-benzoyl-α-D-galactopyranosyl-(1→1)-2′,3′,4′,6′,-tetra-*O*-benzoyl-α-D-glucopyranoside (**9**) 42
**References** 43

**Scheme S1. Synthetic route for 4-fluoro-4-deoxy-trehalose.** *Reagents and conditions:* a) 8.5 eq benzoyl chloride, -40 °C, 2hr, room temp , 48 hr, b) 10 eq. pyridine, 2 eq. triflic anhydride, 0 °C, then room temp, 3 hr, c) 7 eq. sodium nitrite, DMF, room temp., 22 hr, d) 2.5 eq. diethyaminosulfur trifluoride (DAST), 40 °C, 72 hr e) 0.2 M methanolic NaOMe, room temp., 14 hr.

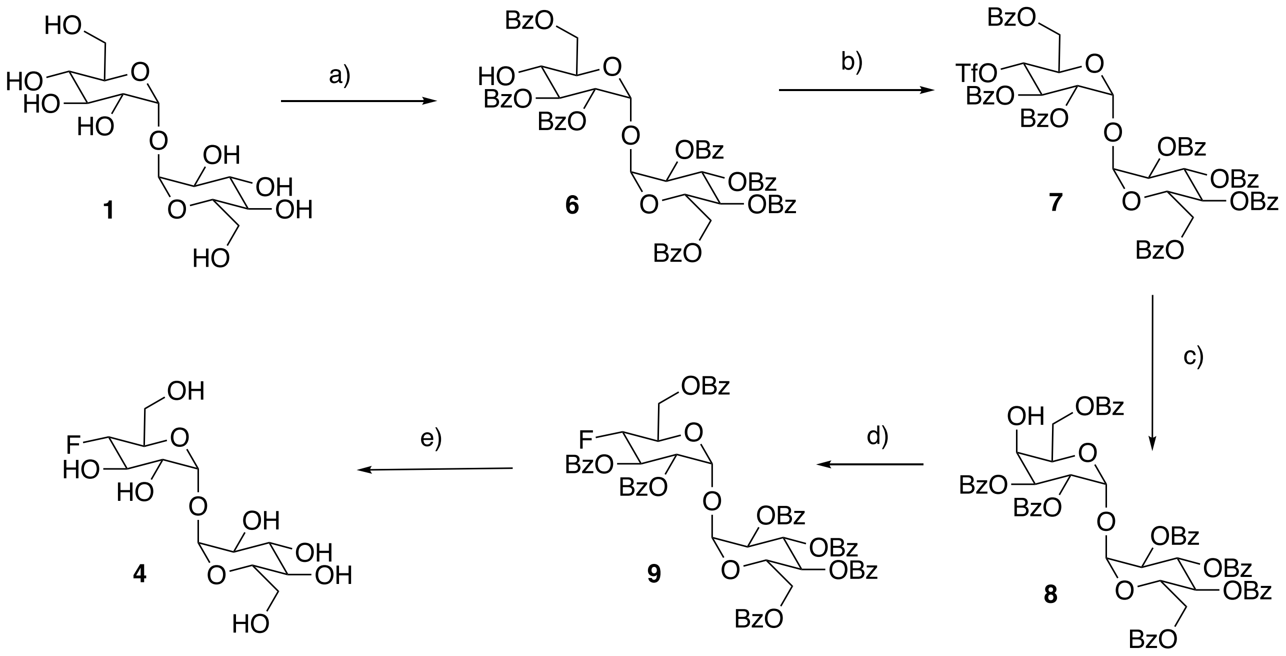

**Figure S1: Binding affinities for LpqY**. Binding of trehalose, 2F-Tre, 3F-Tre, 4F-Tre and 6F-Tre to LpqY measured by microscale thermophoresis (MST). FNorm (%) is the normalized fluorescence signal of the change in MST signal. Error bars represent standard deviations from at least three independent experiments.

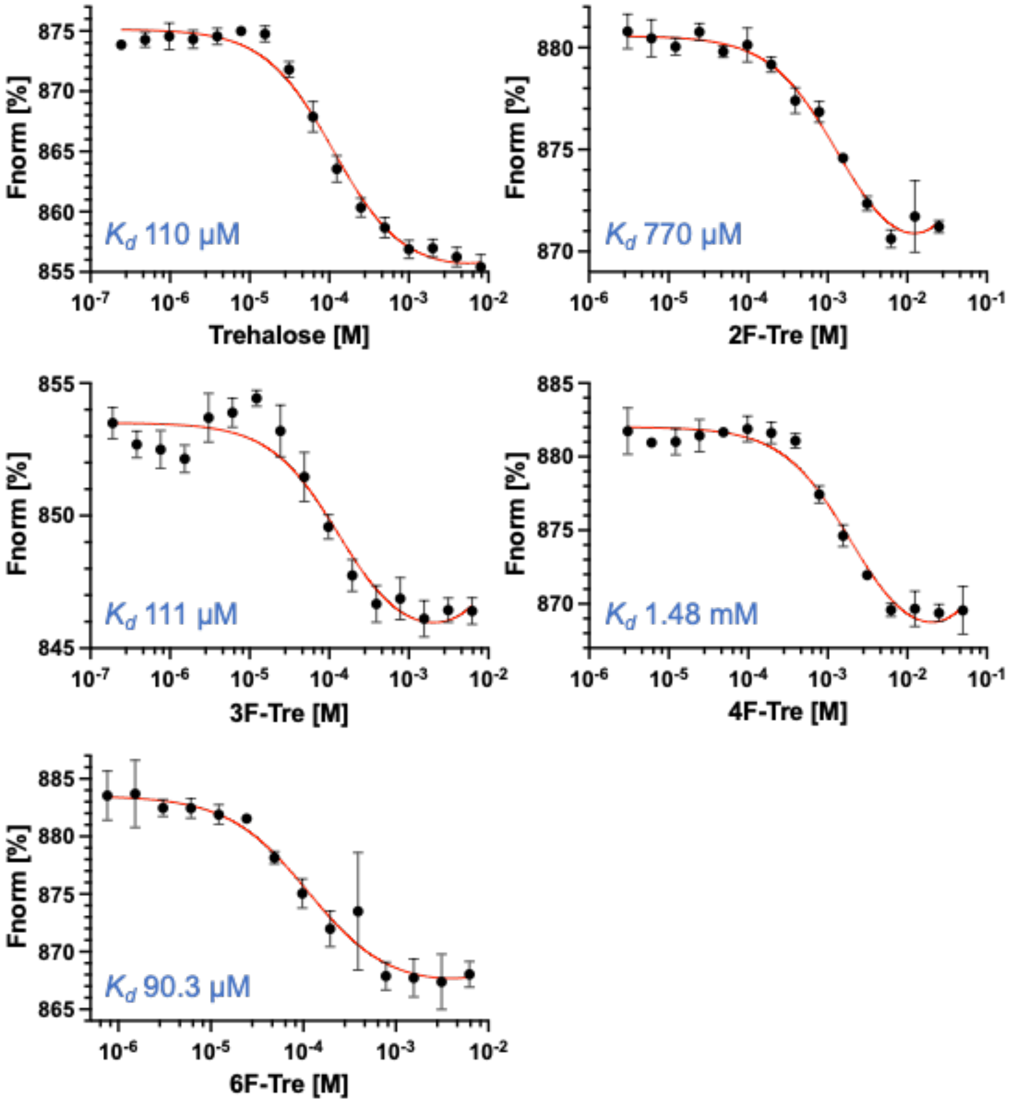

**Fig. S2. STD NMR study of the binding of *Mtr* LpqY with 2F-Tre, 4F-Tre and 6F-Tre with protein irradiation in the aliphatics spectral region.** STD NMR build-up curves for (A) 2F-Tre, (B) 4F-Tre and (C) 6F-Tre, in complex with *Mtr* LpqY. Temperature 5°C for (**A**) and (**B**), and 30°C for (**C**). Saturation frequency set at 0.53 ppm for (**A**) and (**B**), and 0.84 ppm for (**C**).

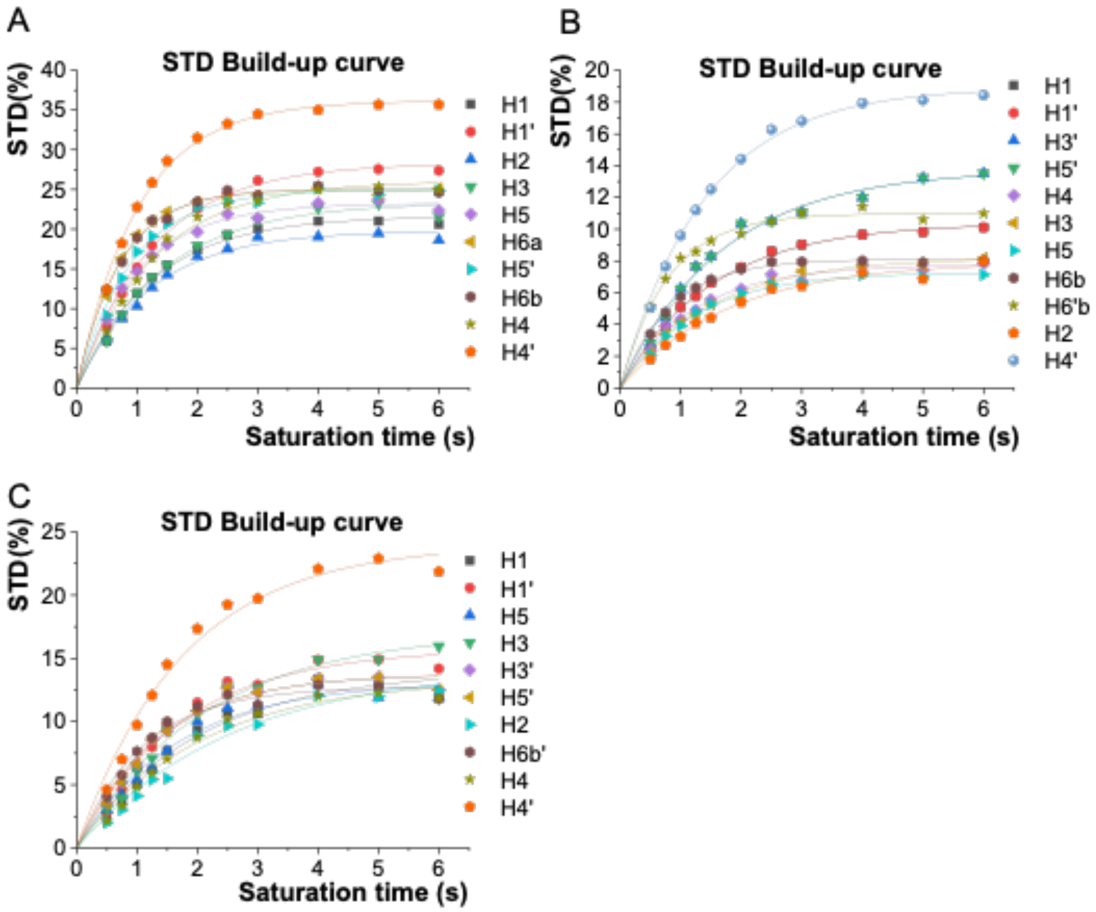

**Fig. S3. STD NMR *for Mtr* LpqY with 2F-Tre, 4F-Tre and 6F-Tre with protein irradiation in the aromatics spectral region.** STD NMR build-up curves for (A) 2F-Tre, (B) 4F-Tre and (C) 6F-Tre in complex with *Mtr* LpqY. Temperature 5°C for (**A**) and (**B**), and 30°C for (**C**). Saturation frequency set at 7.0 ppm for (**A**) and (**B**), and 7.24 ppm for (**C**).

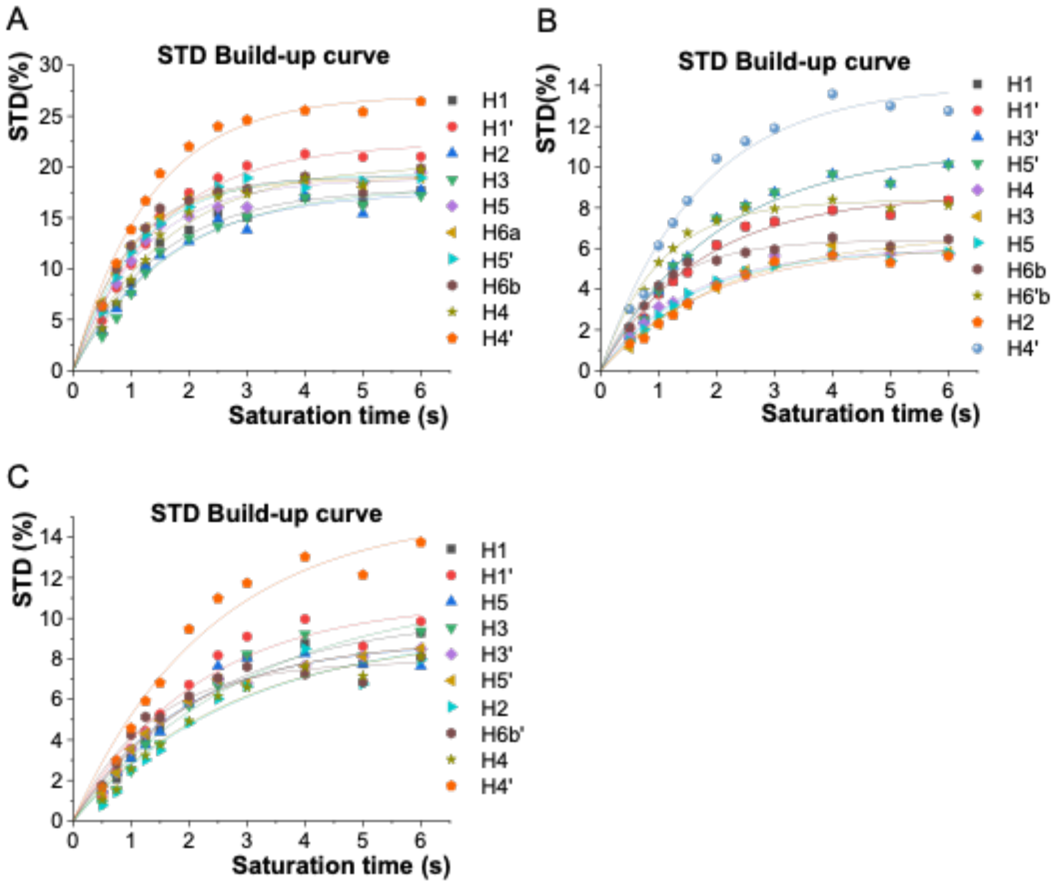

**Table S1. STD NMR data for the binding of *Mtr* LpqY with 2-fluoro-trehalose** with protein irradiation in aliphatics (**A**) or in aromatics (**B**). Temperature 5°C. Saturation frequency set at 0.53 ppm for (**A**) and at 7.00 ppm for (**B**).

**A**

| | $k_{sat}$ ($s^{-1}$) | $STD^{max}$ | $STD_0$ ($s^{-1}$) | $STD_{rel}$ (%) |
|---|---|---|---|---|
| **H1'** | 0,79 | 28,30 | 22,29 | 63 |
| **H4'** | 0,98 | 36,18 | 35,27 | 100 |
| **H5'** | 1,10 | 24,99 | 27,57 | 78 |
| **H6'** | -- | n.d. | -- | -- |
| **H1** | 0,80 | 21,63 | 17,23 | 49 |
| **H2** | 0,82 | 19,79 | 16,26 | 46 |
| **H3** | 0,71 | 23,36 | 16,62 | 47 |
| **H4** | 0,78 | 26,06 | 20,38 | 58 |
| **H5** | 1,00 | 23,22 | 23,14 | 66 |
| **H6a** | 1,41 | 24,93 | 35,10 | 100 |
| **H6b** | 1,38 | 25,05 | 34,50 | 98 |

**B**

| | $k_{sat}$ ($s^{-1}$) | $STD^{max}$ | $STD_0$ ($s^{-1}$) | $STD_{rel}$ (%) |
|---|---|---|---|---|
| **H1'** | 0,68 | 22,31 | 15,21 | 75 |
| **H4'** | 0,75 | 27,08 | 20,28 | 100 |
| **H5'** | 0,91 | 19,25 | 17,42 | 86 |
| **H6'** | -- | n.d. | -- | -- |
| **H1** | 0,68 | 17,89 | 12,22 | 60 |
| **H2** | 0,63 | 17,59 | 11,06 | 55 |
| **H3** | 0,59 | 18,03 | 10,63 | 52 |
| **H4** | 0,63 | 20,24 | 12,83 | 63 |
| **H5** | 0,83 | 18,89 | 15,67 | 77 |
| **H6a** | 1,02 | 18,81 | 19,19 | 95 |
| **H6b** | 1,02 | 19,03 | 19,40 | 96 |

**Table S2. STD NMR data for the binding of *Mtr* LpqY with 3-fluoro-trehalose at a single saturation time.** STD factors and relative STDs obtained at a single saturation time (6 s) for 3-deoxy-3-fluoro-α,α′-trehalose in complex with *Mtr* LpqY with protein irradiation in aliphatics (**A**) or in aromatics (**B**). Temperature 30°C. Saturation frequency set at 0.84 ppm for (**A**) and at 7.24 ppm for (**B**).

**A**

| | STD factor | $STD_{rel}$ (%) |
|---|---|---|
| **H1'** | 2,79 | 79 |
| **H2'** | 2,79 | 79 |
| **H4'** | 3,55 | 100 |
| **H6'** | n.d. | -- |
| **H1** | 2,48 | 70 |
| **H2** | 2,06 | 58 |
| **H4** | 1,53 | 43 |
| **H5** | 1,53 | 43 |
| **H6b** | 1,96 | 55 |

**B**

| | STD factor | $STD_{rel}$ (%) |
|---|---|---|
| **H1'** | 1,66 | 79 |
| **H2'** | 1,78 | 85 |
| **H4'** | 2,10 | 100 |
| **H6'** | n.d. | -- |
| **H1** | 1,58 | 75 |
| **H2** | 1,71 | 81 |
| **H4** | 1,37 | 65 |
| **H5** | 1,02 | 49 |
| **H6b** | 1,31 | 62 |

**Table S3. STD NMR data for the binding of *Mtr* LpqY with 4-fluoro-trehalose** with protein irradiation in aliphatics (**A**) or in aromatics (**B**). Temperature 5°C. Saturation frequency set at 0.53 ppm for (**A**) and at 7.00 ppm for (**B**).

**A**

| | $k_{sat}$ ($s^{-1}$) | $STD^{max}$ | $STD_0$ ($s^{-1}$) | $STD_{rel}$ (%) |
|---|---|---|---|---|
| **H1'/H1** | 0,66 | 10,38 | 6,80 | 49 |
| **H3’/H5'** | 0,59 | 13,75 | 8,17 | 59 |
| **H4'** | 0,71 | 18,92 | 13,48 | 97 |
| **H6’b** | 1,27 | 10,99 | 13,92 | 100 |
| **H2** | 0,56 | 7,95 | 4,42 | 32 |
| **H3** | 0,67 | 8,20 | 5,49 | 39 |
| **H4** | 0,83 | 7,76 | 6,44 | 46 |
| **H5** | 0,82 | 7,29 | 5,95 | 43 |
| **H6b** | 1,20 | 8,11 | 9,72 | 70 |

**B**

| | $k_{sat}$ ($s^{-1}$) | $STD^{max}$ | $STD_0$ ($s^{-1}$) | $STD_{rel}$ (%) |
|---|---|---|---|---|
| **H1’/H1** | 0,58 | 8,61 | 5,01 | 61 |
| **H3’/H5'** | 0,50 | 10,79 | 5,38 | 65 |
| **H4'** | 0,59 | 14,04 | 8,28 | 100 |
| **H6'b** | 0,99 | 8,36 | 8,28 | 100 |
| **H2** | 0,53 | 6,12 | 3,22 | 39 |
| **H3** | 0,46 | 6,72 | 3,09 | 37 |
| **H4** | 0,65 | 5,99 | 3,92 | 47 |
| **H5** | 0,63 | 5,97 | 3,76 | 45 |
| **H6b** | 0,99 | 6,40 | 6,36 | 77 |

**Table S4. STD NMR data for the binding of *Mtr* LpqY with 6-fluoro-trehalose** with protein irradiation in aliphatics (**A**) or in aromatics (**B**). Temperature 30°C. Saturation frequency set at 0.84 ppm for (**A**) and at 7.24 ppm for (**B**).

**A**

| | $k_{sat}$ ($s^{-1}$) | $STD^{max}$ | $STD_0$ ($s^{-1}$) | $STD_{rel}$ (%) |
|---|---|---|---|---|
| **H1'** | 0,58 | 15,77 | 9,16 | 67 |
| **H4'** | 0,57 | 24,02 | 13,72 | 100 |
| **H3'/H5'** | 0,74 | 13,76 | 10,13 | 74 |
| **H6'b** | 0,92 | 12,72 | 11,69 | 85 |
| **H1** | 0,50 | 14,05 | 7,00 | 51 |
| **H2** | 0,39 | 14,32 | 5,59 | 41 |
| **H3** | 0,46 | 17,15 | 7,96 | 58 |
| **H4** | 0,49 | 13,39 | 6,58 | 48 |
| **H5** | 0,60 | 13,17 | 7,90 | 58 |

**B**

| | $k_{sat}$ ($s^{-1}$) | $STD^{max}$ | $STD_0$ ($s^{-1}$) | $STD_{rel}$ (%) |
|---|---|---|---|---|
| **H1'** | 0,46 | 10,85 | 4,98 | 78 |
| **H4'** | 0,42 | 15,21 | 6,39 | 100 |
| **H3'/H5'** | 0,51 | 8,94 | 4,54 | 71 |
| **H6'b** | 0,73 | 7,89 | 5,79 | 91 |
| **H1** | 0,41 | 10,13 | 4,18 | 65 |
| **H2** | 0,34 | 9,50 | 3,27 | 51 |
| **H3** | 0,32 | 11,36 | 3,66 | 57 |
| **H4** | 0,36 | 9,30 | 3,37 | 53 |
| **H5** | 0,52 | 8,85 | 4,63 | 72 |

**Fig. S4. Differential Epitope Mapping by STD NMR of *Mtr* LpqY with 2-fluoro-trehalose, 3-fluoro-trehalose, 4-fluoro-trehalose and 6-fluoro-trehalose.** Differential Epitope Mapping histograms with multifrequency irradiation (0.53 ppm/7.0 ppm) for **A**) 2F-Tre and **C**) 4F-Tre. A different multifrequency irradiation (0.84 ppm/7.24 ppm) was used for **B**) 3F-Tre (analysis carried out at a single saturation time of 6 s) and **D**) 6F-Tre in complex with *Mtr* LpqY. Positive DEEP-STD factors (ΔSTDs) indicate proximity towards aliphatic side chains in the binding site and are shown in orange, whereas negative ΔSTDs indicate proximity towards aromatic side chains in the binding site and are in blue. ΔSTD values were calculated as previously described.[1]

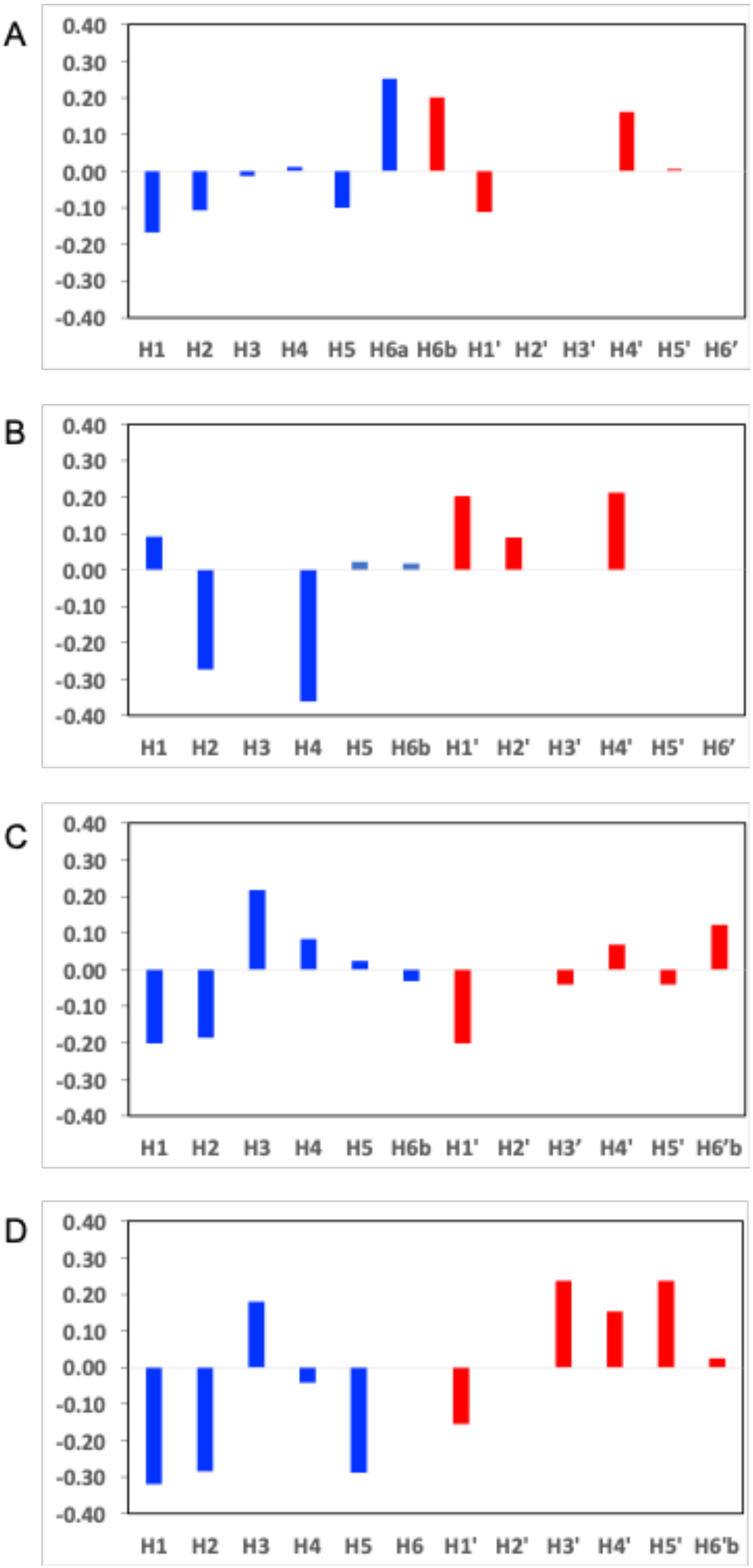

**Fig S5. Molecular dynamic simulations *Mtr* LpqY with 2-fluoro-trehalose, 3-fluoro-trehalose, 4-fluoro-trehalose and 6-fluoro-trehalose**. Evolution of the ligand backbone RMSD (green) with respect to the protein binding site (*i.e.* 5 Å from the ligand) and evolution of the protein backbone RMSD (blue) over 100 ns of MD simulation of the complex between *Mtr* LpqY. (a) 2F-Tre, (b) 3-F-Tre, (c) 4F-Tre, and (d) 6-F-Tre

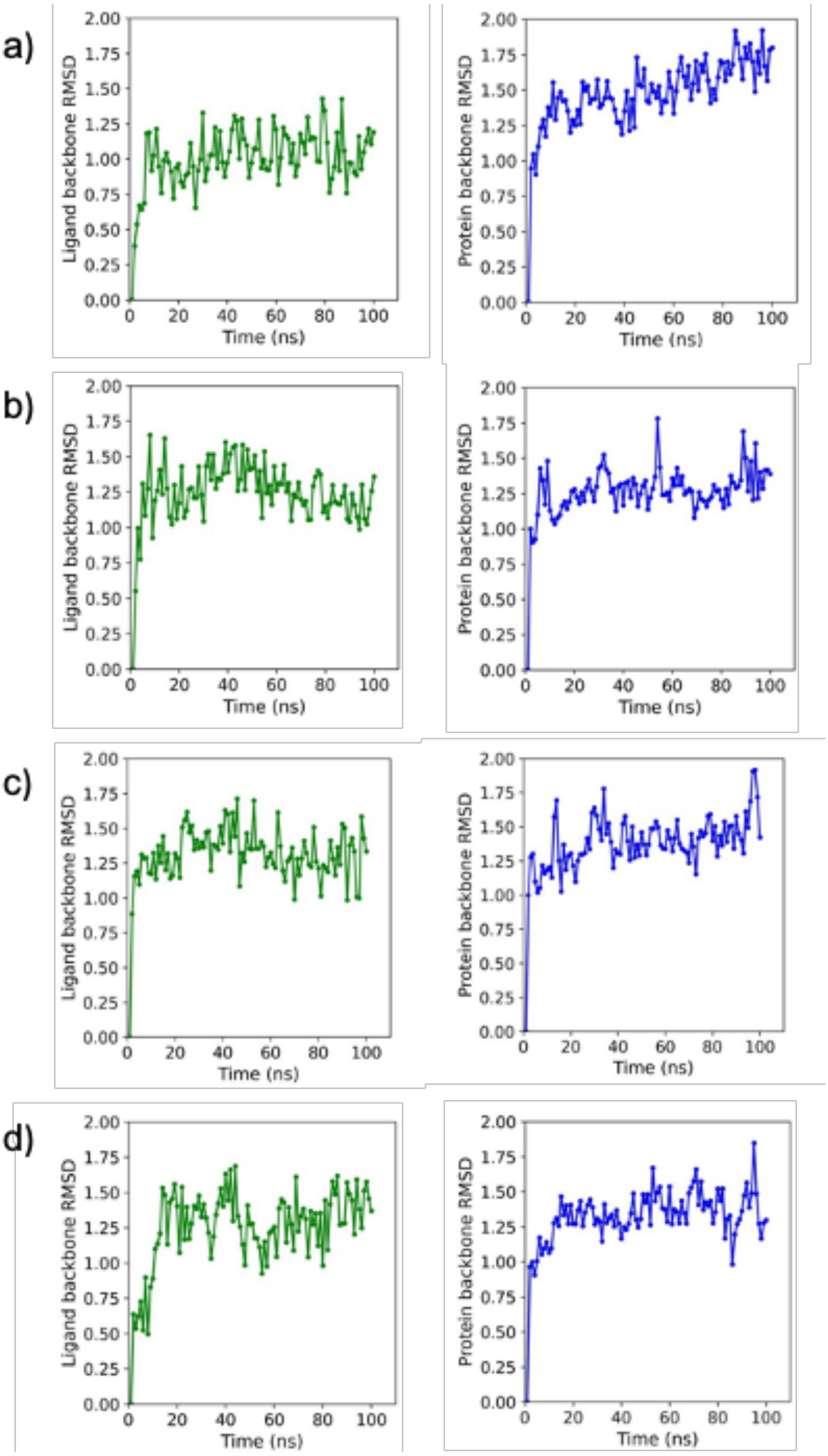

**Fig S6**. **Comparison of the binding orientations of the F-Tre analogues**. Close up superposition showing the binding orientation of the trehalose ligand in stick representation (orange carbon atoms), (PDB 7APE), with the 100ns molecular dynamic snapshots of 2F-Tre (blue carbon atoms), 3F-Tre (magenta carbon atoms), 4F-Tre (grey carbon atoms) and 6F-Tre (green carbon atoms). LpqY is shown in cartoon representation (white, PDB 7APE) and Arg404, which is at the base of the binding cavity, is highlighted. Oxygen, red; nitrogen, blue; fluorine, pale blue

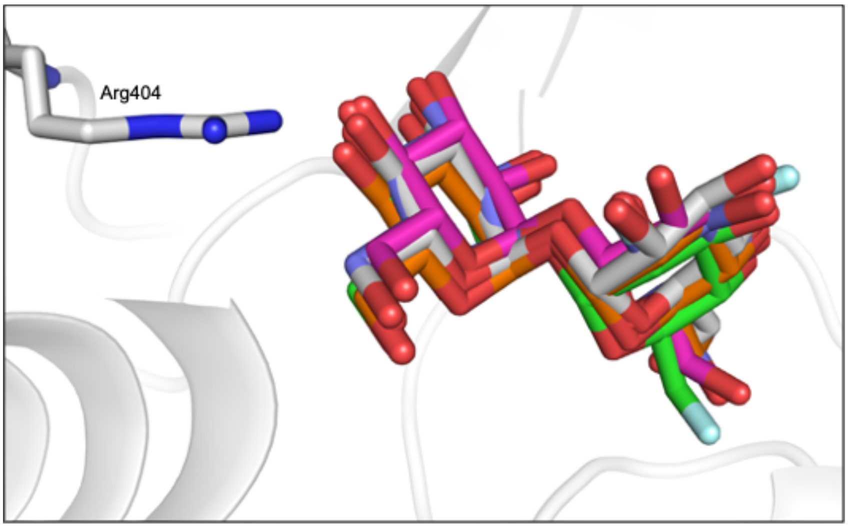

**Fig. S7. Ion chromatography traces of F-Tre labelled *Mycobacterium tuberculosis.*** *Mtb* was cultured in the presence of F-Tre analogues **2-5** (0 – 200 µM) and the cytosolic extracts analysed by high performance anion exchange chromatography with pulsed amperometric detection (HPAEC-PAD). The F-Tre cytosolic concentration was determined from the respective calibration curves for each F-Tre standard.

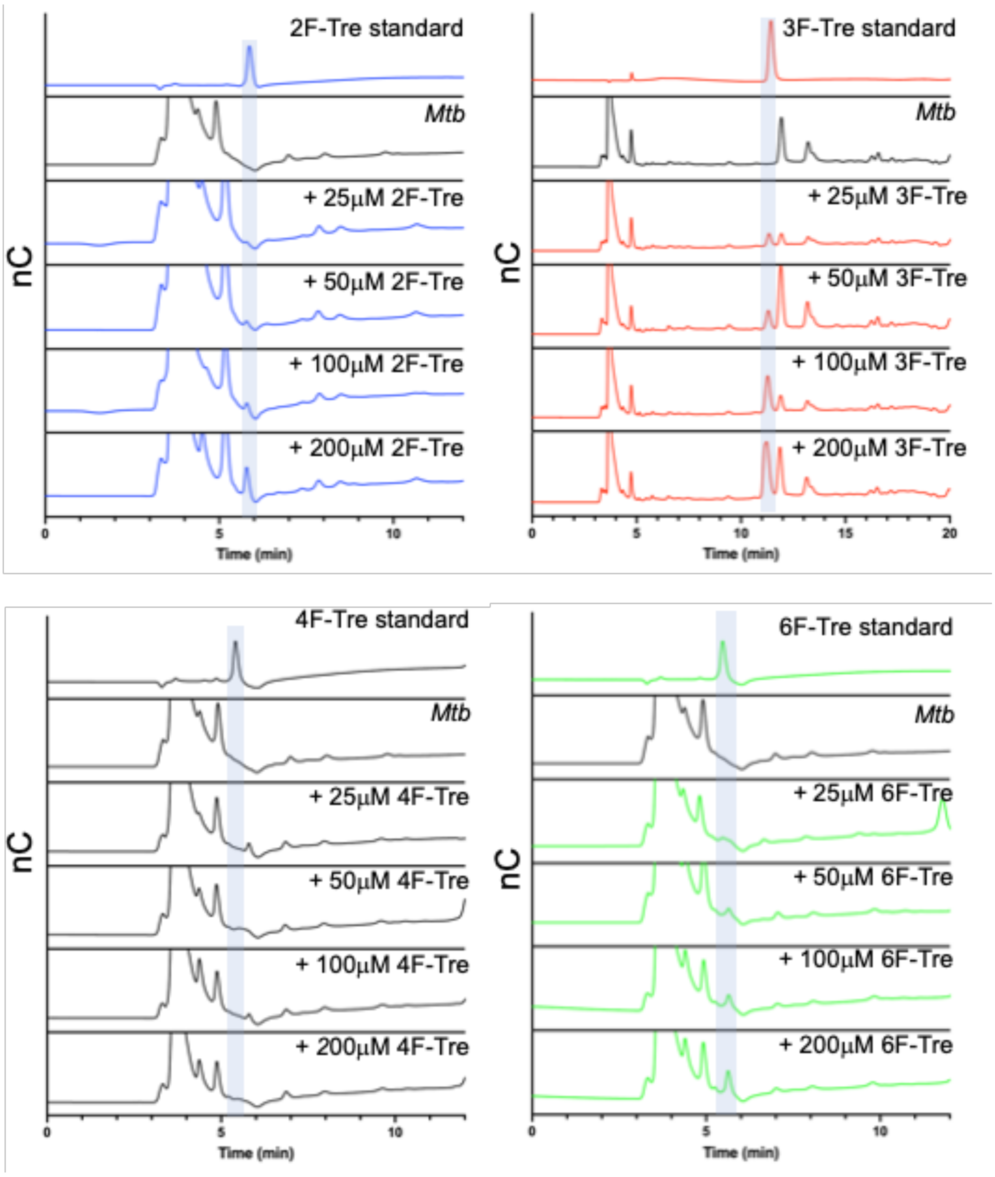

**Table S5.** ***Mtb*** **cytosolic concentrations of F-Tre analogue uptake**

| Time (h) | 2F-Tre (2) (μM) | 3F-Tre (3) (μM) | 4F-Tre (4) (μM) | 6F-Tre (5) (μM) |
|---|---|---|---|---|
| 0 | 0 ± 0 | 0 ± 0 | 0 | 0 ± 0 |
| 0.5 | 636 ± 33 | 250 ± 102 | 0 | 342 ± 16 |
| 1 | 838 ± 52 | 423 ± 256 | 0 | 797 ± 224 |
| 1.5 | 1144 ± 250 | 1030 ± 335 | 0 | 1392 ± 83 |
| 2 | 1717 ± 151 | 1628 ± 25 | 0 | 1804 ± 111 |
| 4 | 2917 ± 105 | 3016 ± 102 | 0 | 2945 ± 234 |
| 6 | 4597 ± 236 | 5872 ± 297 | 0 | 4694 ± 237 |
| 8 | 6639 ± 337 | 7892 ± 72 | 0 | 6843 ± 43 |

Error bars denote the standard deviation from duplicate experiments.

**Fig. S8. Ion chromatography traces of F-Tre labelled *Mycobacterium Mtb.*** *Mtb* was cultured in the presence of F-Tre analogues **2-5** (100 µM) and the cytosolic extracts analysed at the time points by high performance anion exchange chromatography with pulsed amperometric detection (HPAEC-PAD). The F-Tre cytosolic concentration was determined from the respective calibration curves from each F-Tre standard.

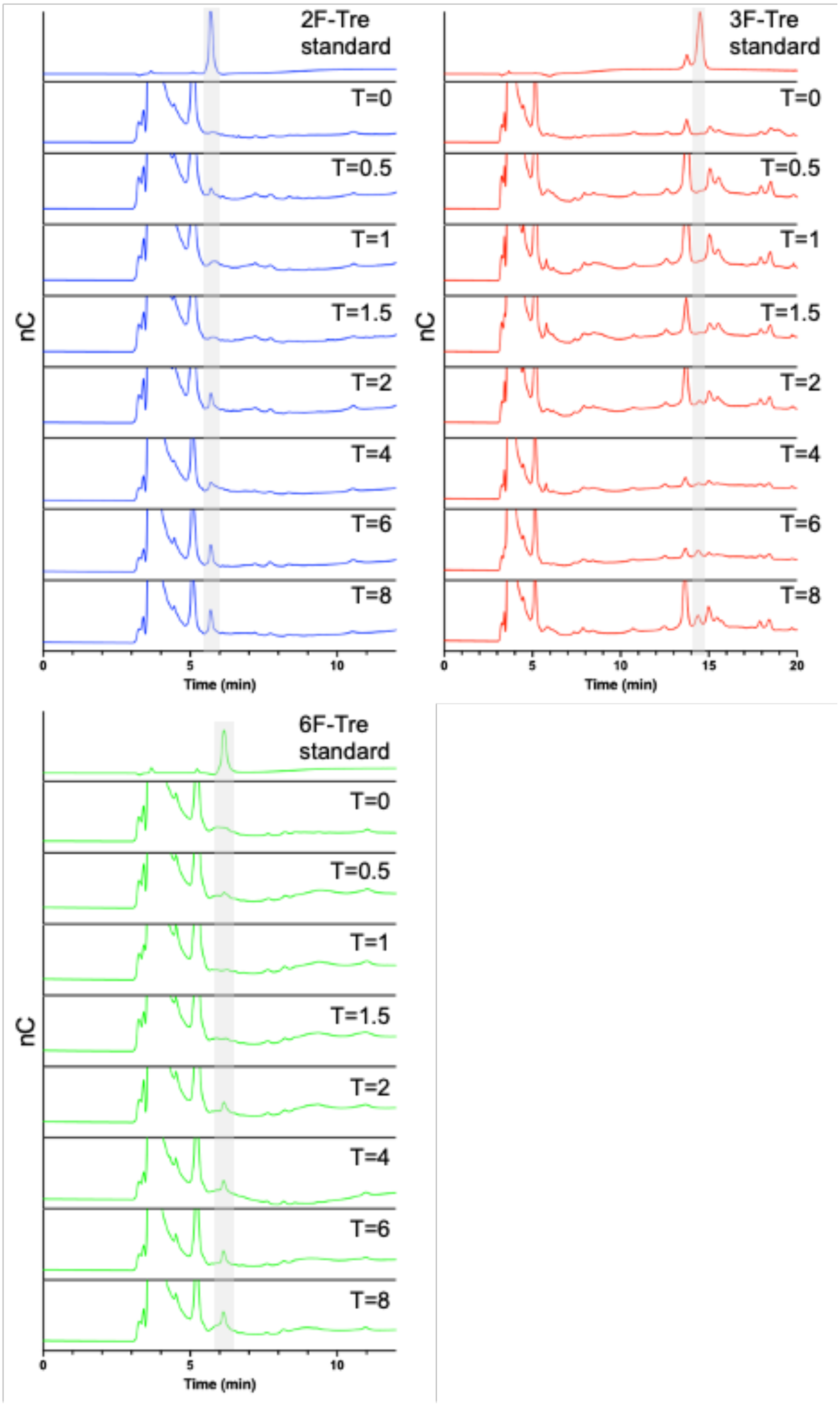

**Fig. S9. Ion chromatography traces of hydrolysed lipid extracts F-Tre labelled *Mycobacterium Mtb.*** To complement our TLC studies which identified new spots in F-Tre labelled *Mtb* cells, free sugars released from the lipid extracts of F-Tre labelled cells were analysed by high performance anion exchange chromatography with pulsed amperometric detection (HPAEC-PAD) A) 2F-Tre, B) 3F-Tre, C) 4F-Tre. 2F-Tre was detected in samples labelled with this analogue. We speculate that 3F-Tre and 6F-Tre are present, however the presence of a peak from *Mtb* control cells that runs at the same time as the 3F-Tre and 6F-Tre standards hindered our ability to confirm 3F-Tre and 6F-Tre in lipid extracts using this method.

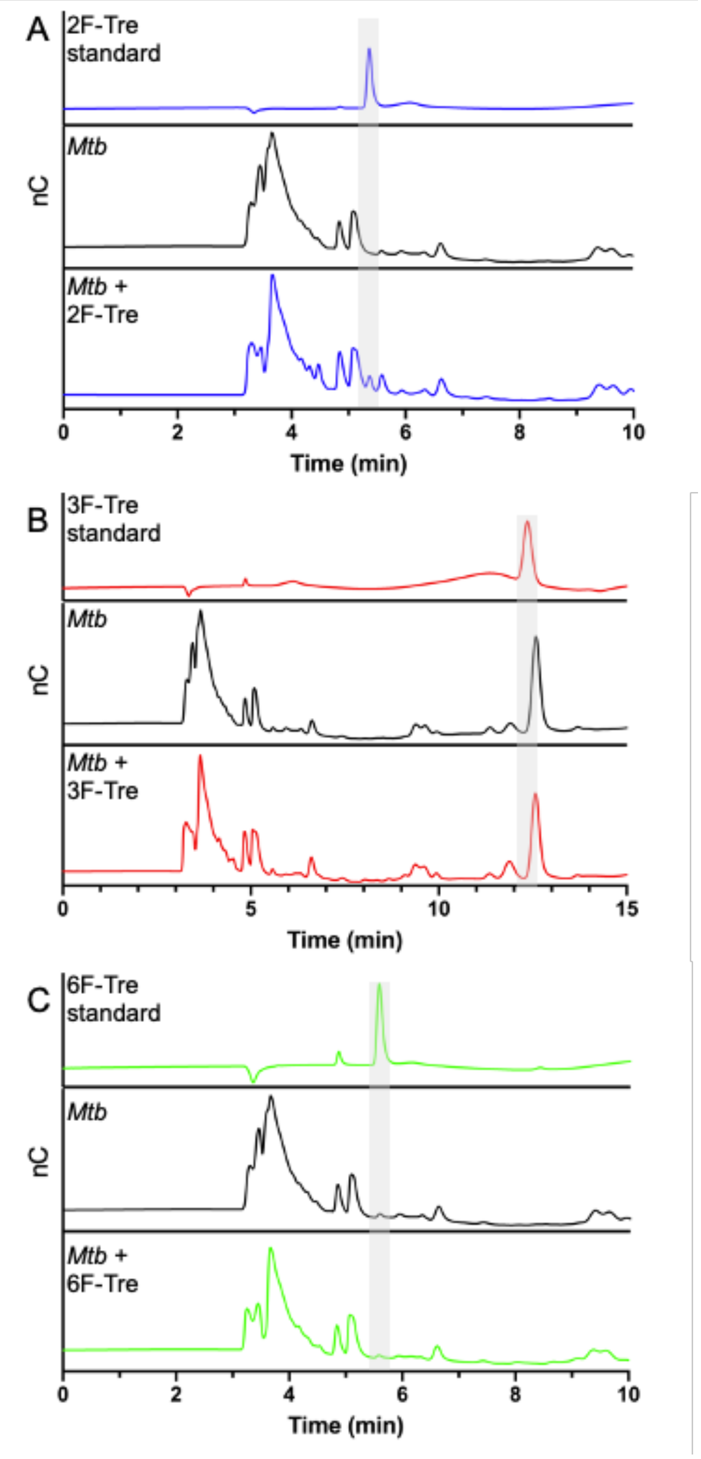

**Fig. S10. F-Glc metabolite analysis of *Mtb* labelled with F-Tre analogues.** *Mtb* was cultured in the presence of (A) 2F-Tre, (B) 3F-Tre or (C) 6F-Tre (100 µM) and the cytosolic extracts analysed by high performance anion exchange chromatography with pulsed amperometric detection (HPAEC-PAD).

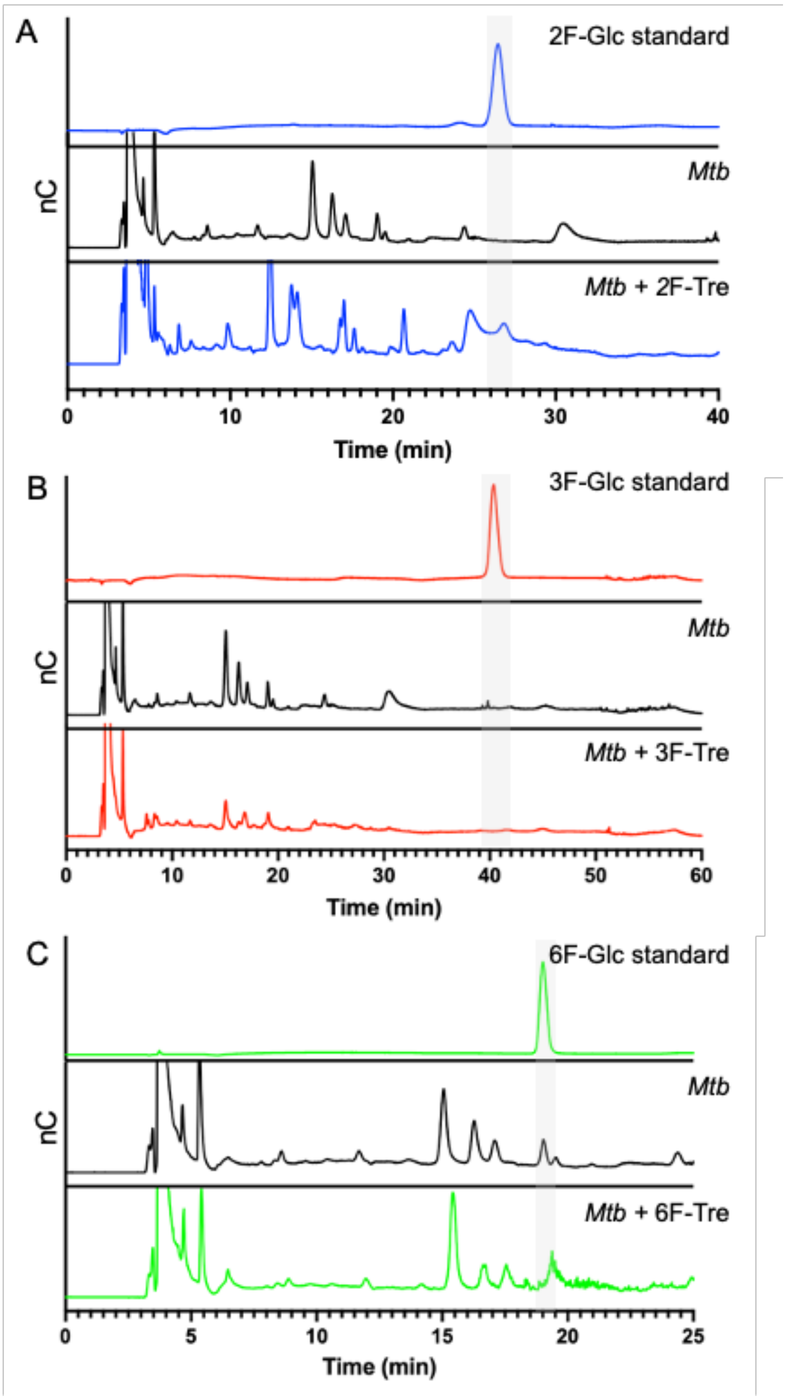

**Fig. S11. F-Tre analogue uptake analysis in the ESKAPE pathogens, *Eschericia coli* and *Bacillus subtilis*.** The bacterial strains were cultured for four doubling times in the presence of (A) 2F-Tre, (B) 3F-Tre, (C) 4F-Tre or (D) 6F-Tre (100 μM) and the cytosolic extracts analysed by high performance anion exchange chromatography with pulsed amperometric detection (HPAEC-PAD). No F-Tre analogue uptake was detected in any strain.

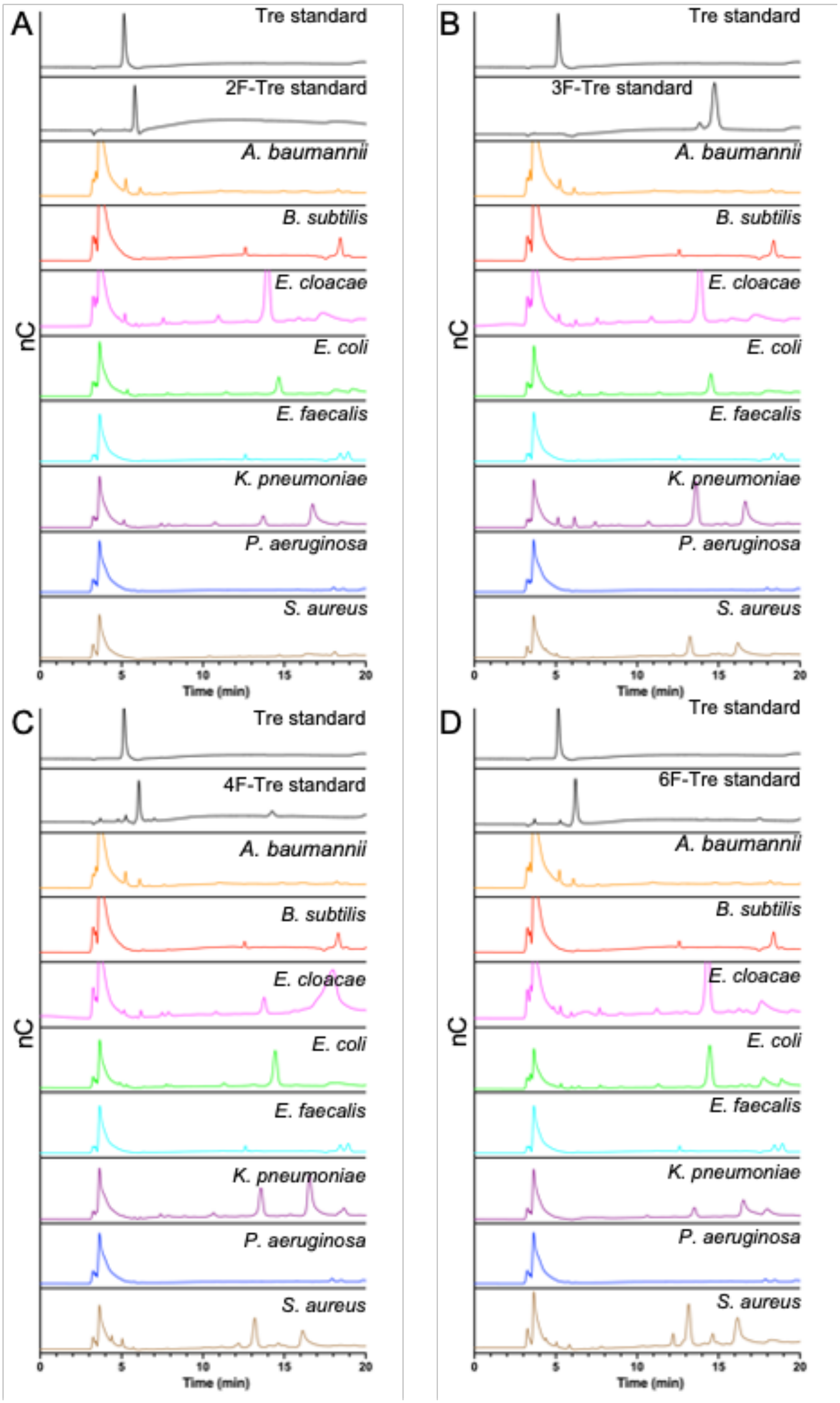

**Fig. S12. SIMS mass spectra of *Mtb* cells treated with 2F-Trehl, 3F-Trehl, 6F-Trehl and non treated control cells, from m/z 1-50.** *Mtb* was grown to $OD_{600}$ 1-1.2 in the presence of 2F-Tre, 3F-Tre or 6F-Tre (100 µM), and a non-treated control. Mass spectra recorded uisng ThermoFisher Scientific Scios Ga source Dualbeam system equipped with a Hiden electrostatic quadrupole secondary ion mass spectrometer (EQS) operating in negative ion mode.

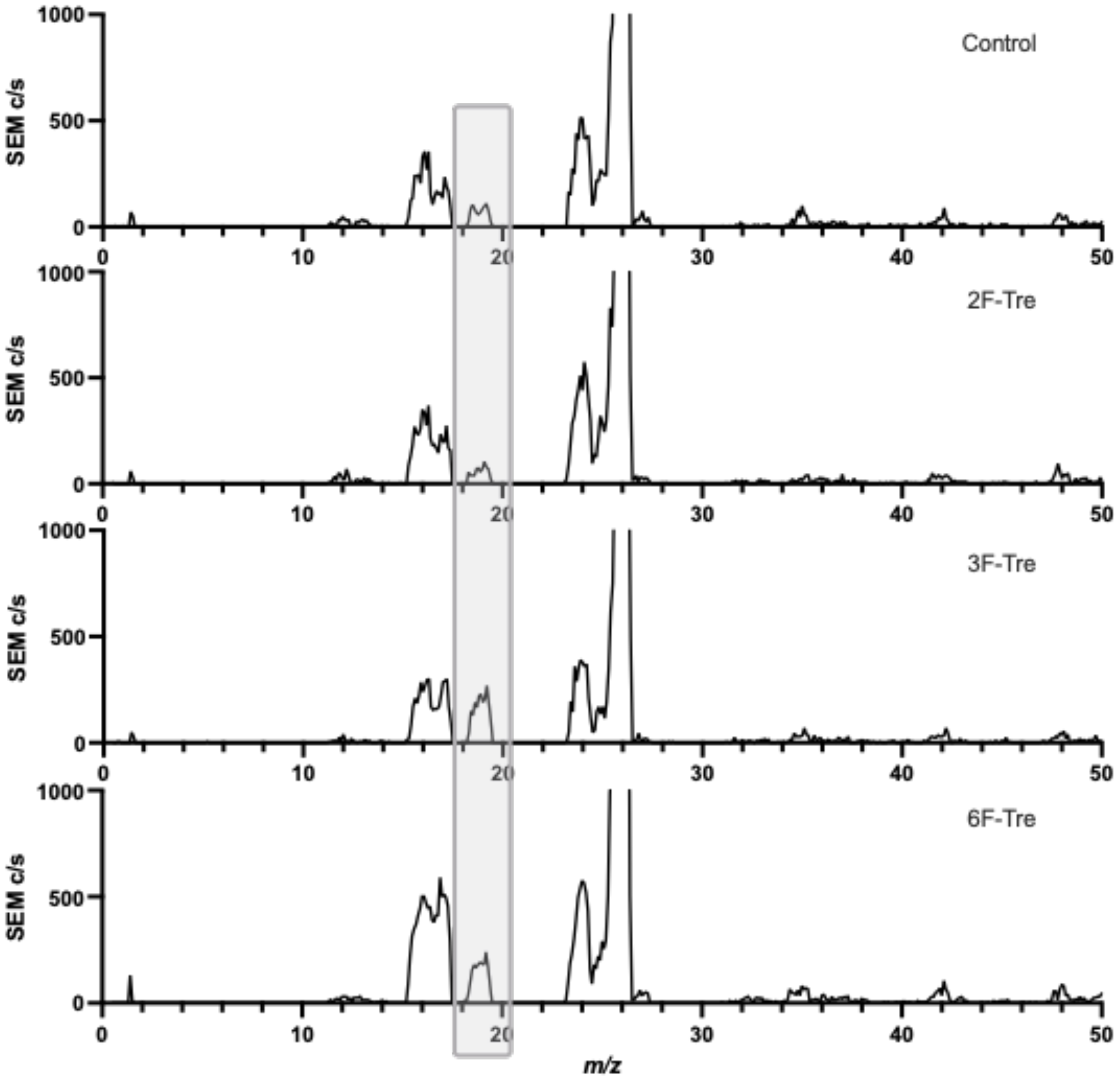

### Experimental

#### General Information and Procedures

Unless stated, the chemicals and solvents, including anhydrous solvents, used in these syntheses were used as supplied and without further purification. 2-fluoro-2-deoxy-glucose, 3-fluoro-3-deoxy-glucose, 6-fluoro-6-deoxy-glucose and UDP-glucose were purchased from Carbosynth. Trehalose was purchased from Acros Organics. Methanol (MeOH), dichloromethane (DCM), pyridine, ethyl acetate (EtOAc), toluene, triethylamine, and magnesium sulfate ($MgSO_4$) were purchased from Fisher Scientific at laboratory reagent grade. Anhydrous *N,N*-dimethylformamide (DMF) >99.8%, anhydrous dichloromethane (DCM), deuterium oxide ($D_2O$) 99.9%, sodium nitrite 98% and triflic anhydride 99% were purchased from Sigma-Aldrich. Deuterochloroform ($CDCl_3$) 99.8% and deuteromethanol (MeOD) 99.8% were purchased from Apollo Scientific. Purified Trehalose Monomycolate (TMM), NR-48784, and purified Trehalose Dimycolate (TDM), NR-14844, were obtained through BEI Resources, NIAID, NIH.

#### $^{1}$H NMR, $^{13}$C NMR and MS data

Proton ($^{1}$H), carbon ($^{13}$C) and fluorine ($^{19}$F) NMR spectra were obtained at 298 K on a Bruker DPX-400 instrument. $^{19}$F NMRs were proton decoupled. NMRs were fully assigned using COSY, HSQC and HMBC. $^{1}$H NMR chemical shifts are quoted in parts per million (ppm), using the residual solvent as the internal standard ($^{1}$H $D_2O$ = 4.79 ppm). Coupling constants (J) are reported in hertz (Hz) with the following abbreviations: s, singlet; d, doublet; t, triplet; q, quartet; quin, quintet; m, multiplet; br, broad. Mass spectra were recorded on a Bruker Esquire 2000 spectrometer using electrospray ionisation (ESI). *M/z* values are reported in Daltons (Da).

#### Bacterial strains, cell lines, culture conditions and chemicals

*Mycobacterium tuberculosis* H37Rv and *Mycobacterium bovis* BCG (ATCC-35734) were routinely grown at 37 °C in Middlebrook 7H9 broth supplemented with 0.2 % glycerol, 0.05 % Tween 80 and 10% albumin-dextrose-catalase (ADC) or on Middlebrook 7H10 plates supplemented with 0.5 % glycerol and 10% oleic acid-albumin-dextrose-catalase (OADC). The gene deletion mutant and complemented strains: *M. bovis BCG* ΔLpqY-SugABC and *M. bovis* BCG ΔLpqY-SugABC:pMV306_LpqY-SugABC, a gift from Professor Rainer Kalscheuer (HHU Dusseldorf, Germany), were cultured with the addition of hygromycin (50 mgL) or hygromycin (50 mg/L) plus kanamycin (20mg/L) respectively. *Acinetobacter baumannii* (ATCC-19606), *Enterobacter cloacae* (ATCC-13045), *Escherichia coli* (ATCC-25922), *Klebsiella pneumoniae* (ATCC-700603), *Pseudomonas aeruginosa* (PA01) and *Staphylococcus aureus* (USA300) were routinely grown in nutrient broth at 37 °C. *Bacillus subtilis* (168CA) and *Enterococcus faecalis* (ATCC-51299) were grown in brain heart infusion (BHI) broth at 37 °C. PBST is phosphate buffered saline supplemented with 0.05% Tween 80.

**Expression and purification of TreT**

The trehalose synthase (TreT) enzyme from *Thermoproteus tenax* was overexpressed and purified as described previously[1]. *Escherichia coli* Top 10 were transformed with the treT_pBADHisA expression plasmid (a gift from Dr B Swarts (Central Michigan University, USA)) and grown at 37°C to an optical density at 600nm (OD600) of 0.6-0.8 in Terrific Broth (Difco) supplemented with 100μg/mL ampicillin. Protein production was induced with 1 mM L-arabinose and the cultures were grown at 37°C overnight with shaking (180 rpm). The cells were harvested (4,000 x g, 45 min, 4 °C) and the pellets were resuspended in PBS and frozen at -80 °C. The frozen pellets were resuspended in 50mM $NaH_2PO_4$, 500 mM NaCl, 20 mM imidazole, pH 8.0 (buffer A), Complete Protease Inhibitor Cocktail (Roche) and lysozyme (60 mg) were added and the cells sonicated on ice (MSE Soniprep 150 plus). Following sonication, the cells were centrifugated (39,000 x g, 30 min, 4 °C), the supernatant was filtered (0.45μm filter, EMD Millipore) and loaded onto a pre-equilibrated HisPur $Ni^{2+}$-affinity resin (Thermo Scientific). The column was washed with buffer A (5 column volumes) and TreT was eluted from the $Ni^{2+}$resin with increasing concentrations of imidazole. Fractions containing the TreT protein were dialysed at 4 °C for 16 h (50mM HEPES, 300mM NaCl, pH 8.0 (buffer B) and purified further using size exclusion chromatography (Superdex 200 16/60 column (GE Healthcare)). Purified TreT fractions were pooled, and the protein concentrated to 1.5-5mg/mL (Amicon, 10kDa MWCO) and stored at 4°C.

**Chemoenzymatic synthesis of fluorinated trehalose derivatives**

The enzymatic reaction contained either 2-fluoro-2-deoxy-D-glucose (30 mM), 3-fluoro-3-deoxy-D-glucose (30 mM) or 6-fluoro-6-deoxy-D-glucose (30 mM), UDP-Glucose (45 mM), $MgCl_2$ (40 mM), and TreT (300 μg/mL) in 50 mM HEPES, 300 mM NaCl, pH 8.0, and was incubated at 70 °C for 2 h, with shaking (300 rpm) and then cooled by placing on ice. A 10 kDa centrifugal filter unit was pre-rinsed with deionised water (2 mL) five times by centrifugation (3,200 x g for 20 mins, room temp). The cooled enzymatic reaction was then added and centrifuged at 3,200 x g for 20 mins, room temp, after which the centrifuge filter was washed 3 times with 2 mL deionised water (3200 x g, 20 mins, room temp). The filtrates were combined and Bio-Rad Bio-Rex RG 501-X8 resin (1 g) added, and the mixture stirred at room temperature for 1 h. The mixture was then filtered and the resin washed with 10 mL deionised water. The filtrates were combined and lyophilised to give the product as a white solid. The reactions were monitored by TLC (5:3:2 n-butanol/ethanol/water), stained with 5% $H_2SO_4$ in ethanol and heated to visualise spots containing sugars.

**2-fluoro-2-deoxy-trehalose (2)**

From 19 mg of 2-fluoro-2-deoxy-glucose obtained 16.0 mg (45%) of 2-fluoro-2-deoxy-trehalose as a white solid. $^1$H NMR (400 MHz, $D_2O$) δ 5.42 (1H, d, $J$ = 4.0 Hz, $H^1$), 5.20 (1H, d, $J$ = 3.5 Hz, $H^{1'}$), 4.50 (1H, ddd, $J$ = 4.0, 9.5 Hz, $J_{H,F}$ = 49.0 Hz, $H^2$), 4.11 (1H, dt, $J$ = 9.5 Hz, $J_{H,F}$ = 13.5 Hz, $H^3$), 3.69 – 3.94 (7H, m, $H^{3'}$, $H^5$, $H^{5'}$, $H^{6ab}$, $H^{6ab'}$), 3.65 (1H, dd, $J$ = 3.5, 10.5 Hz, $H^{2'}$), 3.50 (1H, t, $J$ = 9.5 Hz, $H^4$), 3.44 (1H, t, $J$ = 9.5 Hz, $H^{4'}$). $^{13}$C NMR (100 MHz, $D_2O$) δ 94.0 (C1'), 91.2 (d, $J_{C,F}$ = 21.5 Hz, $C^1$), 89.5 (d, $J_{C,F}$ = 188 Hz, $C^2$), 72.6 ($C^{5'}$), 72.2 ($C^5$), 71.1 (d, $J_{C,F}$ = 17.0 Hz, $C^3$), 70.9 ($C^{2'}$), 69.6 ($C^{4'}$), 69.1 ($C^{3'}$), 69.0 ($C^4$), 60.47, 60.27 ($C^6$ and $C^{6'}$). $^{19}$F NMR (400 MHz, $D_2O$) δ -201.1. *m/z* ($ES^-$): $[M-H]^-$ calcd. for $C_{12}H_{21}O_{10}^-$, 343.1; found 343.1.

**3-fluoro-3-deoxy-trehalose (3)**

From 25 mg of 3-fluoro-3-deoxy-glucose obtained 19.6 mg (42%) of 3-fluoro-3-deoxy-trehalose as a white solid. $^1$H NMR (400 MHz, $D_2O$) δ 5.25 (1H, t, $J$ = 4.0 Hz, $H^1$), 5.19 (1H, d, $J$ = 4.0 Hz, H1'), 4.65 – 4.87 (1H, m, $H^3$), 3.94 (1H, ddd, $J$ = 4.0, 9.5 Hz, $J_{H,F}$ = 13.0 Hz, $H^2$), 3.69 – 3.90 (8H, m, $H^{3'}$, $H^4$, $H^5$, $H^{5'}$, $H^{6ab}$, $H^{6ab'}$), 3.65 (1H, dd, $J$ = 4.0, 10.5 Hz, $H^{2'}$), 3.46 (1H, t, $J$ = 10.0 Hz, $H^{4'}$). $^{13}$C NMR (100 MHz, $D_2O$) δ 94.4 (d, $J_{C,F}$ = 170 Hz, $C^3$), 93.4, 93.4 ($C^1$ and $C^{1'}$), 72.5 ($C^{3'}$), 72.2 ($C^{5'}$), 71.65 (d, $J_{C,F}$ = 7.0 Hz, $C^5$), 70.9 ($C^{2'}$), 69.6 ($C^{4'}$), 69.5 (d, $J$ = 25.0 Hz, $C^2$), 67.85 (d, $J_{C,F}$ = 18.0 Hz, $C^4$), 60.5, 60.0 ($C^6$ and $C^{6'}$). $^{19}$F NMR (400 MHz, $D_2O$) δ -199.7. *m/z* ($ES^-$): $[M-H]^-$ calcd. for $C_{12}H_{21}O_{10}^-$, 343.1; found 343.1.

**6-fluoro-6-deoxy-trehalose (5)**

From 19 mg of 6-fluoro-6-deoxy-glucose obtained 17.3 mg (49%) of 6-fluoro-6-deoxy-trehalose as a white solid. $^1$H NMR (400 MHz, $D_2O$) δ 5.22 (1H, d, $J$ = 4.0 Hz, $H^1$), 5.19 (1H, d, $J$ = 4.0 Hz, $H^{1'}$), 4.58 – 4.78 (2H, m, $H^{6ab}$), 3.98 (1H, dd, $J$ = 10.0 Hz, $J_{H,F}$ = 29.0 Hz, $H^5$), 3.81 – 3.91 (4H, m, $H^3$, $H^{3'}$, $H^{5'}$, $H^{6a'}$), 3.77 (1H, dd, $J$ = 5.5, 12.0 Hz, $H^{6b'}$), 3.66 (2H, td, $J$ = 4.0, 10.5 Hz, $H^2$ and $H^{2'}$), 3.56 (1H, t, $J$ = 9.5 Hz, $H^4$), 3.46 (1H, t, $J$ = 9.5 Hz, $H^{4'}$), $^{13}$C NMR (100 MHz, $D_2O$) δ 93.6, 93.5 ($C^1$ and $C^{1'}$), 82.07 (d, $J_{C,F}$ = 168 Hz, $C^6$), 72.5 ($C^3$), 72.3 ($C^{3'}$), 72.2 ($C^{5'}$), 71.0 (d, $J_{C,F}$ = 10.5 Hz, $C^5$), 70.9, 70.9 ($C^2$ and $C^{2'}$), 69.6 ($C^{4'}$), 68.5 (d, $J_{C,F}$ = 6.5 Hz, $C^4$), 60.5 ($C^{6'}$). $^{19}$F NMR (400 MHz, $D_2O$) δ -235.5. *m/z* ($ES^-$): $[M-H]^-$ calcd. for $C_{12}H_{21}O_{10}^-$, 343.1; found 343.1.

**Chemical synthesis of 4-fluoro-4-deoxy-trehalose (Scheme S1)**

**2,3,6,2′,3′,4′,6',-hepta-*O*-benzoyl-α,α'-D-trehalose (6)**

Trehalose dihydrate (5 g, 13.2 mmol) was suspended in pyridine (75 mL) under nitrogen and cooled to -40 °C. Benzoyl chloride (11.5 mL, 99.1 mmol) was added dropwise, and the reaction was maintained at -40 °C for 2 h before allowing to warm slowly to room temperature and stirred for 16 h. A further portion of benzoyl chloride was added (1.54 mL, 13.2 mmol) and the reaction stirred at room temperature for a further 20 h. The reaction mixture was poured into ice cold 1 M HCl (100 mL) and extracted with EtOAc (3 x 80 mL). The combined organics extracts were washed with sat. $NaHCO_3$ (2 x 80 mL) and brine (80 mL). The organic phase was dried ($MgSO_4$), filtered and concentrated *in vacuo* to give the crude product which was purified by column chromatography (9:1, toluene/EtOAc) to give the desired product as a white foam (1.95g, 14 %). $^1$H NMR (400 MHz, CDCl3) δ 7.81 – 8.13 (14H, m, ArH), 7.22 – 7.62 (21H, m, ArH), 6.28 (1H, t, $J$ = 10.0 Hz, $H^3$), 5.98 (1H, t, $J$ = 9.5 Hz, $H^{3'}$), 5.68 – 5.71 (2H, m, $H^1$, $H^4$), 5.64 (1H, d, $J$ = 4.0 Hz, $H^{1'}$), 5.50 (1H, dd, $J$ = 10.5, 4.0 Hz, $H^2$), 5.46 (1H, dd, $J$ = 10.0, 4.0 Hz, $H^{2'}$), 4.37 (1H, ddd, $J$ = 10.5, 4.5, 3.0 Hz, $H^5$), 4.22 (1H, dd, $J$ = 12.5, 4.0 Hz, $H^{6a'}$), 4.08 (1H, ddd, $J$ = 10.0, 4.0, 2.0 Hz, $H^{5'}$), 3.98 (1H, dd, $J$ = 12.5, 3.0 Hz, $H^{6a}$), 3.85 – 3.91 (2H, m, $H^{6b}$, $H^{6b'}$), 3.82 (1H, t, $J$ = 10.0 Hz, $H^{4'}$); $^{13}$C NMR (100 MHz, CDCl3) δ 167.3, 167.0, 165.9, 165.7, 165.5, 165.0 (C=O), 134.1, 133.9, 133.6, 133.60, 133.5, 133.3, 133.2, 130.3, 130.0, 130.0, 130.0, 129.9, 129.9, 129.8, 129.5, 129.4, 129.2, 129.1, 128.9, 128.8, 128.8, 128.6, 128.5, 128.5, 128.4 (ArC), 93.1, 92.9($C^1$, $C^{1'}$),

73.6, 71.3, 71.2, 70.8, 70.3, 69.3, 68.9, 68.7 ($C^2$, $C^3$, $C^4$, $C^5$, $C^{2'}$, $C^{3'}$, $C^{4'}$, $C^{5'}$), 62.5, 62.0 ($C^6$, $C^{6'}$); *m/z* ($ES^+$): $[M+Na]^+$ calcd. for $C_{61}H_{50}O_{18}Na^+$, 1093.3; found 1093.3.

**2,3,6,-tri-*O*-benzoyl-α-D-galactopyranosyl-(1→1)-2′,3′,4′,6′,-tetra-*O*-benzoyl-α-D-glucopyranoside (8)**

2,3,6,2′,3′,4′,6,-Hepta-*O*-benzoyl-α,α-D-trehalose (1.95 g, 1.82 mmol) was dissolved in DCM (30 mL) under nitrogen and cooled to 0 °C. Pyridine (1.47 mL, 18.2 mmol) and triflic anhydride (613 μL, 3.64 mmol) were added and the reaction allowed to slowly warm to room temperature and stirred for 3 h. The reaction mixture was diluted with DCM (20 mL) and washed with 1 M HCl (50 mL), sat. $NaHCO_3$ (50 mL) and water (50 mL). The organic phase was dried ($MgSO_4$), filtered and concentrated *in vacuo* to give the intermediate triflate as a white solid, which was dissolved in DMF (16 mL) under nitrogen and sodium nitrite (629 mg, 9.11 mmol) was added. The reaction was stirred at room temperature for 16 h. A further portion of sodium nitrite (251 mg, 3.64 mmol) was added, and the reaction stirred for a further 6 h. The reaction was diluted with DCM (60 mL) and washed with water (4 x 60 mL). The organic phase was dried ($MgSO_4$), filtered and concentrated *in vacuo* to give the crude product which was purified by column chromatography (9:1, toluene/EtOAc) to give the desired product as a white solid (670 mg, 34 %). $^1H$ NMR (400 MHz, $CDCl_3$) δ 7.73 – 8.14 (14H, m, Ar<u>H</u>), 7.20 – 7.63 (21H, m, Ar<u>H</u>), 6.25 (1H, t, *J* = 10.0 Hz, $\underline{H}^3$), 5.85 – 5.97 (2H, m, $H^{2'}$, $H^{3'}$), 5.75 (1H, d, *J* = 4.0 Hz, $H^1$), 5.72 (1H, d, *J* = 3.0 Hz, $H^{1'}$), 5.65 (1H, t, *J* = 10.0 Hz, $H^4$), 5.47 (1H, dd, *J* = 10.0, 4.0 Hz, $H^2$), 4.18 – 4.38 (4H, m, $H^4$, $H^5$, $H^{5'}$, $H^{6a'}$), 3.99 – 4.08 (2H, m, $H^{6a}$, $H^{6b'}$), 3.94 (1H, dd, *J* = 12.5, 5.0 Hz, $H^{6b}$); $^{13}C$ NMR (100 MHz, $CDCl_3$) δ 166.2, 166.0, 165.8, 165.7, 165.7, 165.5, 165.1 (<u>C</u>=O), 133.8, 133.7, 133.7, 133.6, 133.4, 133.3, 133.2, 130.0, 130.0, 129.9, 129.9, 129.8, 129.5, 129.4, 129.3, 129.2, 129.0, 128.8, 128.8, 128.8, 128.7, 128.6, 128.5, 128.5, 128.4 (Ar<u>C</u>), 93.1, 92.4 ($C^1$, $C^{1'}$), 71.4, 70.8, 70.4, 69.0, 68.6, 68.6, 68.2, 67.5 ($C^2$, $C^3$, $C^4$, $C^5$, $C^{2'}$, $C^{3'}$, $C^{4'}$, $C^{5'}$), 62.35, 62.18 ($C^6$, $C^{6'}$); *m/z* ($ES^+$): $[M+Na]^+$ calcd. for $C_{61}H_{50}O_{18}Na^+$, 1093.3; found 1093.3.

**4-fluoro-2,3,6,-tri-*O*-benzoyl-α-D-glucopyranosyl-(1→1)-2′,3′,4′,6′,-tetra-*O*-benzoyl-α-D-glucopyranoside (9)**

2,3,6,-tri-*O*-benzoyl-α-D-galactopyranosyl-(1→1)-2′,3′,4′,6′,-tetra-*O*-benzoyl-α-D-glucopyranoside (331 mg, 0.309 mmol) was dissolved in anhydrous dichloromethane (10 mL) under nitrogen at room temperature. Diethyaminosulfur trifluoride (DAST, 102 μL, 0.773 mmol) was then added dropwise. The solution was heated to 40 °C for 72 hrs. The product was diluted with dichloromethane (20 mL), then washed with $NaHCO_3$ (2 x 30 mL) and water (30 mL). The organic layer was dried over anhydrous $MgSO_4$, filtered, and concentrated *in vacuo*. The product was purified by column chromatography on a Biotage Selekt with an Sfär cartridge (10g, silica – 60 μm) (1-20% ethyl acetate in toluene) to give the fluorinated intermediate (208 g, 63%). $^1H$ NMR (400 MHz, CDCl3) δ 7.79 – 8.11 (14H, m, ArH), 7.18 – 7.63 (21H, m, ArH), 6.19 – 6.38 (2H, m, $H^3$, $H^{3'}$), 5.63 – 5.73 (3H, m, $H^1$, $H^{1'}$, $H^{4'}$), 5.51 (1H, dd, *J* = 10.5, 4.0 Hz, $H^2$ or $H^{2'}$), 5.41 (1H, dd, *J* = 10.0, 4.0 Hz, $H^2$ or $H^{2'}$), 4.74 (1H, dt, *J* = 9.4 Hz, $J_{H,F}$ = 50.5 Hz, $H^4$), 4.211 – 4.39 (2H, m, $H^5$, $H^{5'}$), 3.91 – 4.05 (3H, m, $H^{6a}$, $H^{6a'}$,$H^{6b'}$), 3.84 (1H, dd, *J* = 12.5, 4.5 Hz, $H^{6b}$). $^{13}C$ NMR (100 MHz, CDCl3) δ 165.9, 165.5, 165.5, 165.4, 165.4, 165.0 (<u>C</u>=O) 134.1, 133.9, 133.6, 133.5, 133.3, 133.2, 133.2, 129.9, 129.9, 129.8, 129.8, 129.4, 129.18,

129.0, 129.0, 128.9, 128.8, 128.7, 128.6, 128.6, 128.5, 128.5, 128.4, 128.4, 128.3, 128.2 (ArC), 93.0, 92.7 ($C^{1}$, $C^{1'}$), 86.85 (d, $J_{C,F}$ = 189.0 Hz, $C^{4}$), 71.1, 70.7 (d, $J_{C,F}$ = 8.0 Hz), 70.4 (d, $J_{C,F}$ = 20.5 Hz), 70.1, 68.7, 68.7, 67.9 (d, $J_{C,F}$ = 23.0 Hz). ($C^{2}$, $C^{3}$, $C^{5}$, $C^{2'}$, $C^{3'}$, $C^{4'}$, $C^{5'}$), 61.9, 61.6 ($C^{6}$, $C^{6'}$). $^{19}F$ NMR (400 MHz, $D_2O$) δ -197.4 *m/z* ($ES^{+}$): $[M+Na]^{+}$ calcd. for $C_{61}H_{59}FO_{17}Na^{+}$, 1095.3; found 1095.3.

**4-fluoro-4-deoxy-trehalose (4)**

4-fluoro-2,3,6,-tri-*O*-benzoyl-α-D-glucopyranosyl-(1→1)-2′,3′,4′,6′,-tetra-*O*-benzoyl-α-D-glucopyranoside (208 mg, XX mmol) was dissolved in 0.2 M methanolic NaOMe (5 mL) and the reaction was stirred at room temperature for 14 h. Amberlite® IR120 acidic resin (200 mg) was then added and the mixture stirred at room temperature for 30 mins to neutralise the reaction, filtered and concentrated *in vacuo*. The residue was taken up in water (10 mL), washed with petroleum ether (40-60 °C) (3 x 10 mL) and then lyophilised to give the product as a white solid (63.1 mg, 95 %); $^{1}H$ NMR (400 MHz, $D_2O$) δ 5.20 (2H, d, $J$ = 4.0 Hz, $H^{1}$ and $H^{1'}$), 4.38 (1H, dt, $J$ = 9.5 Hz, $J_{H,F}$ = 51.0 Hz, $H^{4}$), 4.16 (1H, dt, $J$ = 9.5 Hz, $J_{H,F}$ = 16.0Hz, $H^{3}$), 3.98 – 4.07 (1H, m, $H^{5}$), 3.73 – 3.92 (6H, m, $H^{3'}$, $H^{5'}$, $H^{6ab}$, $H^{6ab'}$), 3.70 (1H, dd, $J$ = 4.0, 10.0 Hz, $H^{2}$), 3.65 (1H, dd, $J$ = 4.0, 10.0 Hz, $H^{2'}$), 3.46 (1H, t, $J$ = 9.5 Hz, $H^{4'}$). $^{13}C$ NMR (100 MHz, $D_2O$) δ 93.5 ($C^{1'}$), 93.2 ($C^{1}$), 89.12 (d, $J$ = 179.5 Hz, $C^{4}$), 72.5, 72.2 (C3' and C5'), 71.0 ($C^{2'}$), 70.7 (d, $J$ = 17.5 Hz, $C^{3}$), 70.48 (d, $J$ = 8.5 Hz, $C^{2}$), 69.7 ($C^{4'}$), 69.60 (d, $J$ = 23.5 Hz, $C^{5}$), 60.5, 59.8 ($C^{6}$ and $C^{6'}$). $^{19}F$ NMR (400 MHz, $D_2O$) δ -198.3. *m/z* ($ES^{-}$): $[M-H]^{-}$ calcd. for $C_{12}H_{21}O_{10}^{-}$, 343.1; found 343.1.

**Production and purification of *Mtr* LpqY**

*Mtr* LpqY was overexpressed and produced as described previously[2]. In brief, *E. coli* BL21 (DE3) cells containing the *mtr_lpqY_sumo* plasmid were grown at 27 °C to an $OD_{600}$ of 0.4 to 0.6 in Terrific broth medium supplemented with 50 µg/ml kanamycin. Protein production was induced with 1 mM isopropyl-β-thiogalactopyranoside, and the cultures were grown at 16 °C overnight with shaking (180 rpm). The cells were harvested (4,000 *g*, 45 min, 4 °C) and resuspended in lysis buffer (20 mM Tris, 300 mM NaCl, 10% glycerol, pH 7.5 (buffer 1)) and frozen at −80 °C. A complete protease inhibitor tablet (Roche), $MgCl_2$ (5 mM), DNase (2 mg), and lysozyme (20 mg) were added to the resuspended pellet, and sonicated on ice (MSE Soniprep 150 plus). Following centrifugation (39,000*g*, 45 min, 4 °C), the supernatant was filtered (0.45 µm filter) and loaded onto a pre-equilibrated HisPur $Ni^{2}$-NTA affinity resin (Thermo Scientific). The column was washed with buffer 1 and *Mtr* LpqY was eluted with increasing concentrations of imidazole. Fractions containing *Mtr* LpqY were digested with His-tagged SUMO protease (1 h, 30 °C, 300 µg) and dialyzed at 4 °C for 12 h against buffer 1. A second HisPur $Ni^{2}$-NTA affinity resin purification step was undertaken, and fractions containing *Mtr* LpqY were pooled and purified further using size exclusion chromatography (Superdex 200 16/600 column, GE Healthcare) with 50mM HEPES, 300mM NaCl pH 7.5. Purified *Mtr* LpqY was concentrated to 5 to 14 mg/ml (Vivaspin 20; GE Healthcare) and stored at −80 °C.

**Microscale Thermophoresis**

*Mtr* LpqY (2.6 μM) was labelled with the amine reactive RED-NHS dye (3 μM) (second generation, NanoTemper Technologies). Following incubation in the dark for 30 mins with shaking (300 rpm, room temp), excess dye was removed using a Zeba desalting spin column (7KMWCO). The trehalose analogues were prepared in PBS containing 0.05% Tween 20, and the final concentration of the protein in the assay was 500 nM. The samples were loaded into the MonoLith NT.115 standard treated capillaries and incubated for 10 min before analysis using the Monolith NT.115 instrument (NanoTemper Technologies) at 21 °C using the auto-select excitation power (20%) and medium laser power. The binding affinities were calculated using a single-site binding model using the MST Analysis software (version 7.0). All experiments were carried out in triplicate.

**STD-NMR**

All the STD NMR experiments were carried out in PBS $D_2O$ buffer, pH 7.4. The protein concentration was 28 μM and the ligand concentration (2-fluoro-trehalose or 2-deoxy-2-fluoro-α,α′-trehalose, 3-fluoro-trehalose or 3-deoxy-3-fluoro-α,α′-trehalose, 4-fluoro-trehalose or 4-deoxy-4-fluoro-α,α′-trehalose and 6-fluoro-trehalose or 6-deoxy-6-fluoro-α,α′-trehalose) was 1 mM. STD NMR spectra were acquired on a Bruker Avance III 700.25 MHz at 278 K for studies with 2-fluoro-trehalose and 4-fluoro-trehalose, whereas the temperature was 303 K for studies with 3-fluoro-trehalose and 6-fluoro-trehalose. The on- and off-resonance spectra were acquired using a train of 50 ms Gaussian selective saturation pulses using a variable saturation time from 0.5 s to 6 s, and a relaxation delay (D1) of 6 s. The residual protein resonances were filtered using a $T1\rho$-filter of 25 ms. All the spectra were acquired with a spectral width of 9 kHz and 24K data points using 256 scans in saturation times of 0.5, 0.75, 1, 1.25 seconds, 128 scans in 1.5, 2 seconds and 64 scans in 2.5, 3, 4, 5, 6 seconds. The on-resonance spectra were acquired by saturating aliphatic hydrogens, specifically at 0.53 ppm for studies with 2-fluoro-trehalose and 4-fluoro-trehalose, whereas irradiation was at 0.84 ppm for studies with 3-fluoro-trehalose and 6-fluoro-trehalose, but also by saturating of aromatics hydrogens, specifically at 7.0 ppm for studies with 2-fluoro-trehalose and 4-fluoro-trehalose, whereas it was 7.24 ppm for studies with 3-fluoro-trehalose and 6-fluoro-trehalose, where average chemical shifts used came from those predicted from shiftX2[1] [3] for the aliphatic and aromatic residues present in the binding site of Mtr LpqY, whereas the off-resonance spectra were in all cases acquired by saturating at 40 ppm. To get accurate structural information from the STD NMR data and to minimize any $T_1$ relaxation bias, the STD build-up curves were fitted to the equation $STD(t_{sat}) = STDmax*(1-exp(-k_{sat}*t_{sat}))$, calculating the initial growth rate $STD_0$ factor as STDmax*ksat and then normalizing all of them to the highest value. DEEP-STD factors were obtained as previously described [4] with all saturation times (0.5, 0.75, 1, 1.25, 1.5, 2, 2.5, 3, 4, 5 and 6 s) for 2-fluoro-trehalose, 4-fluoro-trehalose and 6-fluoro-trehalose and at a single saturation time (6 s) for 3-fluoro-trehalose on aliphatic or aromatic regions (0.53 or 7 ppm for 2-fluoro-trehalose and 4-fluoro-trehalose, whereas 0.84 or 7.24 ppm 3-fluoro-trehalose and 6-fluoro-trehalose).

### Molecular dynamics

#### *Input preparation and equilibration*

The initial coordinates of the four *Mtr* LpqY–fluorinated trehalose complexes were built from the coordinates of the model of *Mtr* LpqY bound to 6-azido-trehalose[2] by manually modifying the substituents of the trehalose ligand with *Pymol*. The MD simulation setup and equilibration were performed with the BioExcel Building Blocks (*BioBB*) library[5]. The ligands were parametrized and minimized using the *acpype* and *babel* modules, respectively, of BioBB (*biobb_chemistry.acpype* and *biobb_chemistry.babel*). The minimization of the ligands was performed with the steepest descent method and the GAFF force field. The topology of the complexes were generated with the *biobb_amber.leap* module, using the ff14SB force field[6] for the protein and GAFF[7] for the ligand. Subsequently, they were minimized with the *biobb_amber.sander* module using first positional restraints of 50 kcal/mol·Å$^2$ on the protein heavy atoms and, secondly, positional restraints of 500 kcal/mol·Å$^2$ on the ligand to avoid potential changes in ligand orientation due to protein repulsion. Then, each protein-ligand complex was immersed in a TIP3P[8] truncated octahedron water box with a distance from the protein to the box edge of 9.0 Å and Periodic Boundary Conditions, followed by the addition of a 150 mM concentration of NaCl. This gave rise to MD simulation systems of ~ 37,000 atoms. Each solvated system was minimized using the steepest descent protocol and applying positional restraints of 15 kcal/mol·Å$^2$ to the ligand, followed by heating up to 300 K over 2500 steps applying the Langevin thermostat[9] with a collision frequency of 1 ps$^{-1}$ and positional restraints on the ligand of 10 kcal/mol·Å$^2$ (the *biobb_amber.sander* module was used). Next, each system was subjected to NVT followed by NPT equilibration of 100 ps each. A non-bonded interactions cutoff of 10.0 Å, the SHAKE algorithm for constraining the length of bonds involving hydrogen atoms, the Langevin thermostat[9] with a collision frequency of 5 ps$^{-1}$, and smooth positional restraints on the ligand (5 and 2.5 kcal/mol·Å$^2$ for NVT and NPT, respectively) were employed. During the NPT equilibration, a pressure of 1 bar was kept constant using isotropic position scaling with a pressure relaxation time of 2 ps.

#### *MD simulations*

A 100-ns of MD production run was carried out for each complex on a AMD-Ryzen 4xGPU 3070 Computing Cluster using the *pmemd.cuda* module of AMBER 20[10]. The production dynamics was performed at a constant temperature of 300 K, by applying the Langevin thermostat[9] with a collision frequency of 1 ps$^{-1}$, and a constant pressure of 1 bar (using isotropic position scaling with a pressure relaxation time of 1 ps). A non-bonded interactions cutoff of 9.0 Å, periodic boundary conditions (PBC)[11], and the Particle Mesh Ewald method[12] (PME) to account for the long range electrostatic effect were employed. The SHAKE algorithm[13, 14] was also employed, thus allowing 2 fs between time steps. Trajectory coordinates were saved every nanosecond. The analysis of the MD trajectories was performed using the *cpptraj* module (version *4.25.6*) of AMBER 20.[10] The evolution of protein and ligand RMSD over the simulation time was calculated against the first frame of the trajectory. For the protein RMSD, only the backbone atoms were considered for the fit and the calculation. For the ligand, we first aligned the trajectory against the first frame using the protein backbone atoms within 5 Å of the ligand as fitting selection, and subsequently, the RMSD of the ligand backbone was calculated in-place (no superposition) allowing the orientational changes and dynamics of the ligand in the binding site to be determined.

**Analysis of F-Tre uptake at a single-time point**

*M. tuberculosis, M. bovis* BCG, *A. baumannii, B. subtilis, E. cloacae, E. coli, E. faecalis, K. pneumoniae, P. aeruginosa or S. aureus* were cultured in the presence of 2F-Tre, 3F-Tre, 4F-Tre or 6F-Tre (25 μM, 50 μM, 100 μM or 200 μM final concentration) in 5 mL culture volumes, with a starting $OD_{600}$ of 0.05. Controls with either the equivalent volume of water or the equivalent volume of trehalose added were also prepared. Cultures were grown until the optical density at 600 nm ($OD_{600}$) was between 1.0 and 1.2 and the cells then harvested by centrifugation (2,916 *x g*, 22 °C for 5 min for mycobacteria or 10 min for non-mycobacterial species) and washed three times (3 x 5 mL PBST). The pellets were then resuspended in 1 mL $H_2O$ and lysed by mechanical disruption using 0.1 mm zirconia/silica beads (BioSpec Products) on a FastPrep (MP Biomedicals) ribolyser (4 x 45 secs cycles with 90 secs on ice in between). Samples were then centrifuged (16,200 x *g*, 10 min, 22°C) and the supernatant collected, lyophilised, resuspended in 1 mL 18 MΩ $H_2O$ and filtered through a 10-kDa molecular weight cut-off centrifuge filter (Amicon) and the filtrate analysed. HPAEC-PAD was performed on a Dionex ICS5000+ system with a CarboPac PA-20 analytical column (3 mm x 150 mm) and PA-20 guard column (3 mm x 30 mm) kept at 20 °C. Pulsed amperometry with standard quadrupole waveform was used for detection. The system was equipped with an autosampler that was set up to inject 10 μL sample volumes. Multistep gradient elution was performed using a KOH eluent generation cartridge with the gradient conditons shown in Table S6. Authentic standards of trehalose, 2F-Tre, 3F-Tre, 4F-Tre and 6F-Tre were prepared at 100 μM. Chromeleon 7 software (Dionex) was used for data processing.

**Table S6: High performance anion exchange chromatography KOH elution gradient for F-Tre analysis**

| Time (mins) | KOH conc. (mM) | Flow rate (mL/min) |
|---|---|---|
| 0 | 5 | 0.15 |
| 3 | 5 | 0.15 |
| 4 | 5 | 0.40 |
| 18 | 30 | 0.40 |
| 20 | 30 | 0.40 |
| 22 | 5 | 0.15 |

To quantify uptake, the peak area of of 2F-Tre, 3F-Tre and 6F-Tre standards at varying concentrations (0.5 - 50 μM) were measured (Chromeleon 7 software). The peak area was plotted against concentration and simple linear regression was plotted. To determine the concentration of F-Tre analogues in cytosolic samples the area of the peaks of interest was measured (Chromeleon 7 software) and the concentration determined from the calibration plot.

**Time dependent F-Tre uptake**

*M. tuberculosis* was grown to an $OD_{600}$ of 0.8, then 2F-Tre, 3F-Tre, 4F-Tre or 6F-Tre were added to a final concentration of 100 μM. Controls with the equivalent volume of water or equivalent concentration of trehalose were also prepared. 5 mL aliquots were taken at T=0, 0.5, 1, 1.5, 2, 3, 4, 6, 8 h and the cells were harvested by centrifugation (2,916 *x g*, 5 mins, 22 °C) and washed (5 mL PBST) three times. The pellets were then resuspended in 1 mL $H_2O$ and lysed by mechanical disruption using 0.1 mm zirconia/silica beads (BioSpec Products) on a FastPrep (MP Biomedicals) ribolyser (4 x 45 secs cycles with 90 secs on ice in between). Samples were then centrifuged (16,200 x *g*, 10 min, 22°C) and the supernatant collected, lyophilised, resuspended in 1 mL 18 MΩ $H_2O$ and filtered through a 10-kDa molecular weight cut-off centrifuge filter (Amicon). The filtrate was analysed by HPAEC-PAD and quantified as described above.

**Analysis of F-Glc metabolites**

*M. tuberculosis, was* cultured in the presence of 2F-Tre, 3F-Tre, or 6F-Tre (100 μM final concentration) in 5 mL culture volumes, with a starting $OD_{600}$ of 0.05. Controls with either the equivalent volume of water were also prepared. Cultures were grown until the $OD_{600}$ was between 1.0 and 1.2 and the cells then harvested by centrifugation (2,916 *x g*, 22 °C for 5 min) and washed three times (3 x 5 mL PBST). The pellets were resuspended in 1 mL $H_2O$ and lysed by mechanical disruption using 0.1 mm zirconia/silica beads (BioSpec Products) on a FastPrep (MP Biomedicals) ribolyser (4 x 45 secs cycles with 90 secs on ice in between). Samples were then centrifuged (16,200 x *g*, 10 min, 22°C) and the supernatant collected, lyophilised, resuspended in 1 mL 18 MΩ $H_2O$ and filtered through a 10-kDa molecular weight cut-off centrifuge filter (Amicon) and the filtrate analysed. HPAEC-PAD was performed on a Dionex ICS5000+ system with a CarboPac PA-20 analytical column (3 mm x 150 mm) and PA-20 guard column (3 mm x 30 mm) kept at 20 °C. Pulsed amperometry with standard quadrupole waveform was used for detection. The system was equipped with an autosampler that was set up to inject 10 μL sample volumes. Multistep gradient elution was performed using a KOH eluent generation cartridge with the gradient conditions shown in Table S7. Authentic standards of 2F-Glc, 3F-Glc, and 6F-Glc were prepared at 100 μM. Chromeleon 7 software (Dionex) was used for data processing.

**Table S7: High performance anion exchange chromatography KOH elution gradient for F-Glc analysis**

| Time (mins) | KOH conc. (mM) | Flow rate (mL/min) |
|---|---|---|
| 0 | 5 | 0.15 |
| 3 | 5 | 0.15 |
| 4 | 5 | 0.40 |
| 18 | 30 | 0.40 |
| 50 | 30 | 0.40 |
| 52 | 5 | 0.15 |
| 60 | 5 | 0.15 |

**Lipid extraction and analysis**

*M. tuberculosis* and *M. bovis* BCG were cultured in the presence of 2F-Tre, 3F-Tre, 4F-Tre or 6F-Tre (100 μM final concentration) in 25 mL volumes. Controls with equivalent volume of water added, and the equivalent concentration of trehalose were also prepared. Cultures were grown until the $OD_{600}$ was between 1.0 and 1.2 then cells were harvested by centrifugation (2,916 *x g*, 5 mins, 22 °C) and washed (5 mL PBST) three times. The pellets were resuspended in 2 mL in MeOH–0.3% aqueous NaCl (10:1) and 2 mL petroleum ether (60-80 °C) and the samples shaken at 800 rpm at room temperature overnight. The samples were then centrifuged (2187 x *g*, 5 mins, 22 °C) and the top layer collected. A further 2 mL petroleum ether (60-80 °C) was added to the remaining bottom layer and the samples shaken at 800 rpm for 1 hour. The samples were then centrifuged (2187 x *g*, 5 mins, 22 °C) and the top layer collected and combined with the previous fraction and dried to yield the apolar lipids. 2.3 mL of chloroform/methanol/0.3% NaCl (9:10:3) was then added to the remaining lower layer and the samples shaken at 800 rpm at room temp overnight. The samples were then centrifuged (2187 x *g*, 5 mins, 22 °C) and the supernatant collected. The pellet was then resuspended in 750 μL of chloroform/methanol/0.3% NaCl (5:10:4) and the samples shaken at 800 rpm for a further hour. The samples were then centrifuged (2187 x *g*, 5 mins, 22 °C) and the supernatant collected. The pellet was then resuspended in 750 μL of chloroform/methanol/0.3% NaCl (5:10:4) and the samples shaken at 800 rpm for a further hour. The samples were then centrifuged (2187 x *g*, 5 mins, 22 °C) and the supernatant collected. 1.3 mL chloroform and 1.3 mL 0.3% NaCl was added to the combined supernatants and then shaken at 800 rpm at room temp for 5 mins before centrifuging (2187 x *g*, 5 mins, 22 °C). The lower phase was collected and dried to yield the polar lipids. Lipids were analysed by TLC (8:2:0.2 chloroform/methanol/$NH_4OH$) and compared to TMM and TDM standards. Plates were visualised with 5% $H_2SO_4$ in ethanol followed by heating, to detect carbohydrate compounds.

Following TLC analysis the carbohydrate head groups were cleaved and analysed by HPAEC-PAD. The isolated lipid samples were resuspended in anhydrous dichloromethane (0.2 mL), treated with NaOMe/MeOH (2.0 M, 0.2 mL), and stirred vigorously for 16 h at 60 °C. The reaction was neutralized with Amberlite $H^+$ resin to pH 7.0, filtered, and evaporated to dryness. The residue was then resuspended in chloroform (1.0 mL) and extracted three times with 18 MΩ $H_2O$ (1 mL). The combined aqueous layers were lyophilised and resuspended in 300 μL 18 MΩ $H_2O$ and analysed by HPAEC- PAD as above.

**Preparation of samples for focussed ion beam (FIB) secondary ion mass spectrometry (SIMs)**

*M. tuberculosis* was cultured in the presence of 2F-Tre, 3F-Trel, 4F-Tre or 6F-Tre (100 μM final concentration) in 5 mL volumes. Controls with equivalent volume of water added, and the equivalent concentration of trehalose were also prepared. F-Tre was added at an $OD_{600}$ of 0.05 and the cultures were grown until the $OD_{600}$ was between 1.0 and 1.2. The cells were harvested by centrifugation (2,916 *x g*, 5 min, 22 °C) and washed (5 mL PBST) three times, resuspended in PBST (5 mL) and 0.5 mL of the sample was centrifuged (15,871 x g,

5 mins, 22 °C) and the supernatant discarded. The pellet was resuspended in glutaraldehyde (2.5% in PBS, 1 mL) and incubated at 4 °C for 90 mins. The samples were then centrifuged (15,871 x g, 5 mins, 22 °C) and the supernatant removed. The pellet was washed 3 times with PBST (15,871 x g, 5 mins, 22 °C) and twice with deionised water (15,871 x g, 5 mins, 22 °C) and stored at 4 °C prior to imaging.

**Scanning electron microscopy (SEM) and FIB-SIMS**

Glutaraldehyde treated pellets were resuspended in 1 mL $H_2O$ and 5 µL was then spotted onto a copper TEM grid. The sample was left to settle for 2 mins, the excess liquid blotted off with filter paper and the grid was then plunge frozen in liquid ethane (cooled with liquid $N_2$) and lyophilised. SEM imaging was performed using a ThermoFisher Scientific Scios Dualbeam Secondary electron (SE). SEM images were acquired using accelerating voltage 2kV and beam current 100pA. FIB-SIMS was performed using a ThermoFisher Scientific Scios Ga source Dualbeam system equipped with a Hiden electrostatic quadrupole secondary ion mass spectrometer (EQS). To minimise charging effects, samples were first coated with Au/Pd using a Cressington 206HR sputter coater. SIMS mass spectra were acquired by scanning an area of approx. 5 µm x 5 µm, with a dwell time of 100 ns and beam conditions 30 kV, 1 nA. Mass spectra were acquired with the EQS operating in negative ion mode, using a mass dwell time of 200 ms and step size 0.1 amu. FIB-SIMS mapping was performed using the FIB $Ga^+$ beam operating at 30 kV with beam current 100 pA. Operating in negative ion mode, the EQS was set to map counts of m/z = 19 ($19F^-$). FIB-SIMS mapping was performed with a pixel dwell time 1 ms with image size 800 pixels for areas approximately 50 µm x 50 µm. Image intensity of the FIB-SIMS maps correspond to counts s-1 of m/z 19 ($19F^-$), providing qualitative distribution of fluorine. Pairs of ($19F^-$) FIB-SIMS maps for positive and control samples were acquired on the same day using the same ion beam conditions. For each positive/control pair of FIB-SIMS maps, the control image colourmap was scaled so that image intensity/colours correspond to the same image intensity scale as the positive sample. FIB-SIMS map images were smoothed using a gaussian blur 3x3 smoothing filter.

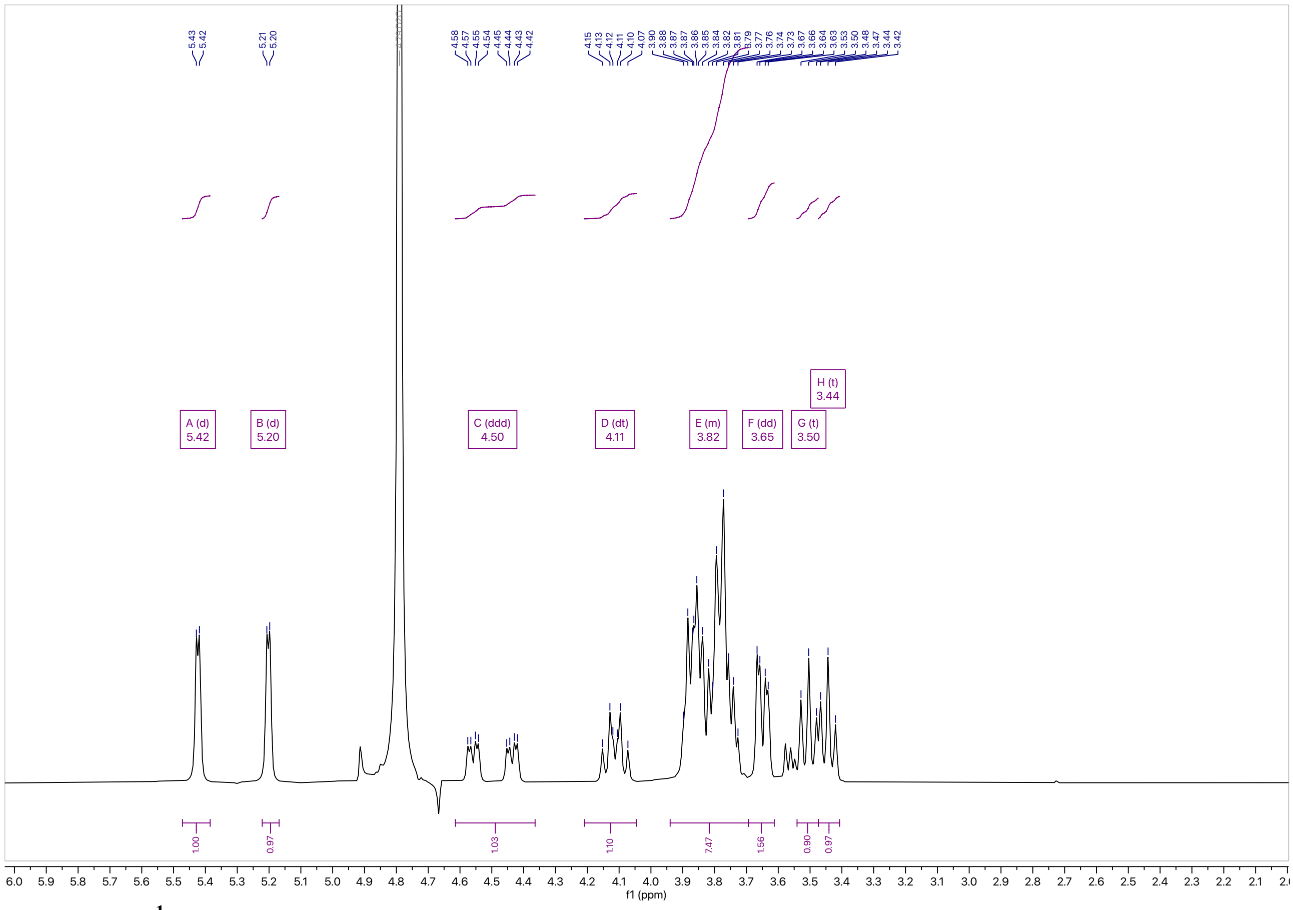

**Figure S13. $^{1}$H NMR 2-fluoro-2-deoxy-trehalose (2)**

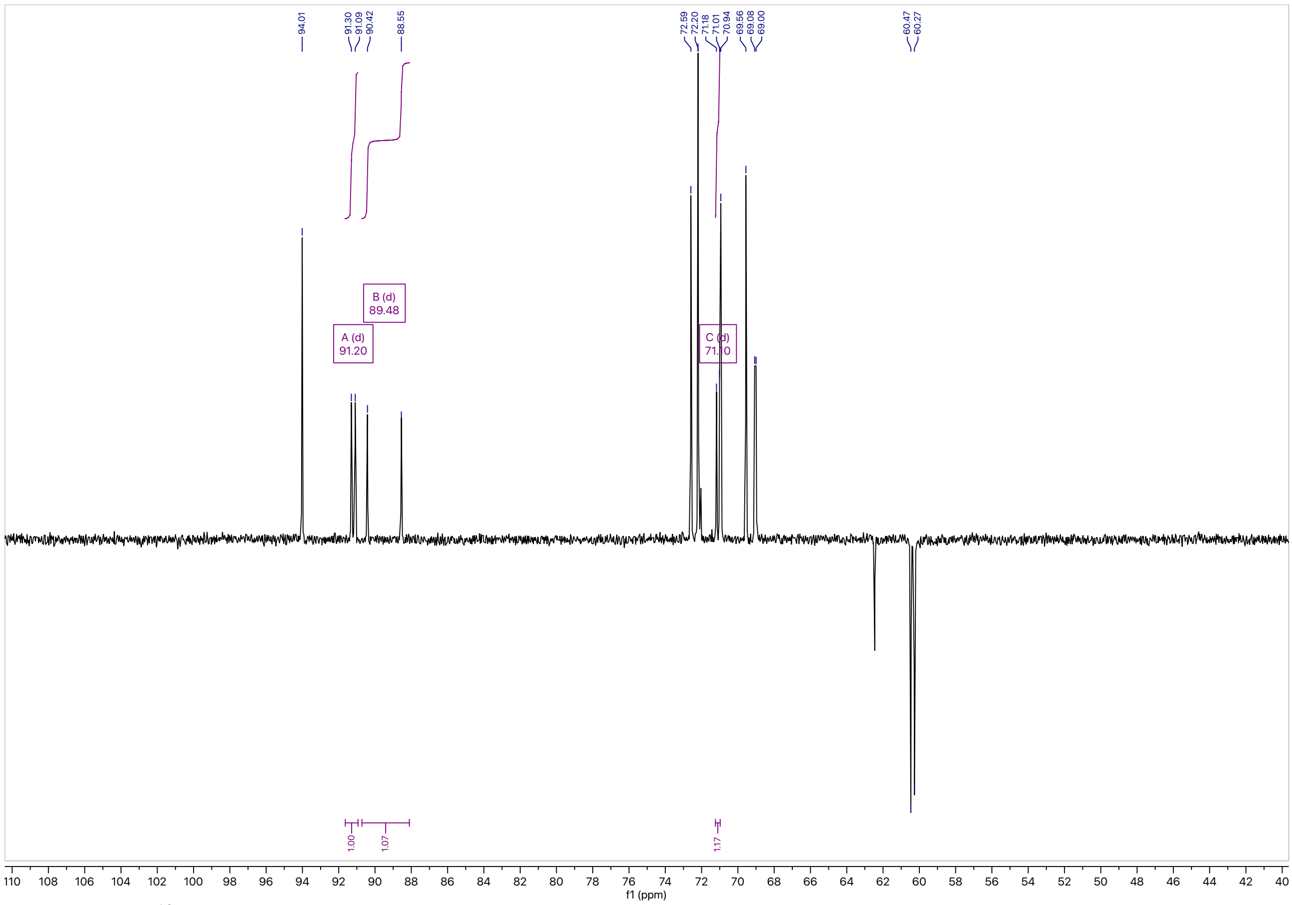

**Figure S14. $^{13}$C NMR 2-fluoro-2-deoxy-trehalose (2)**

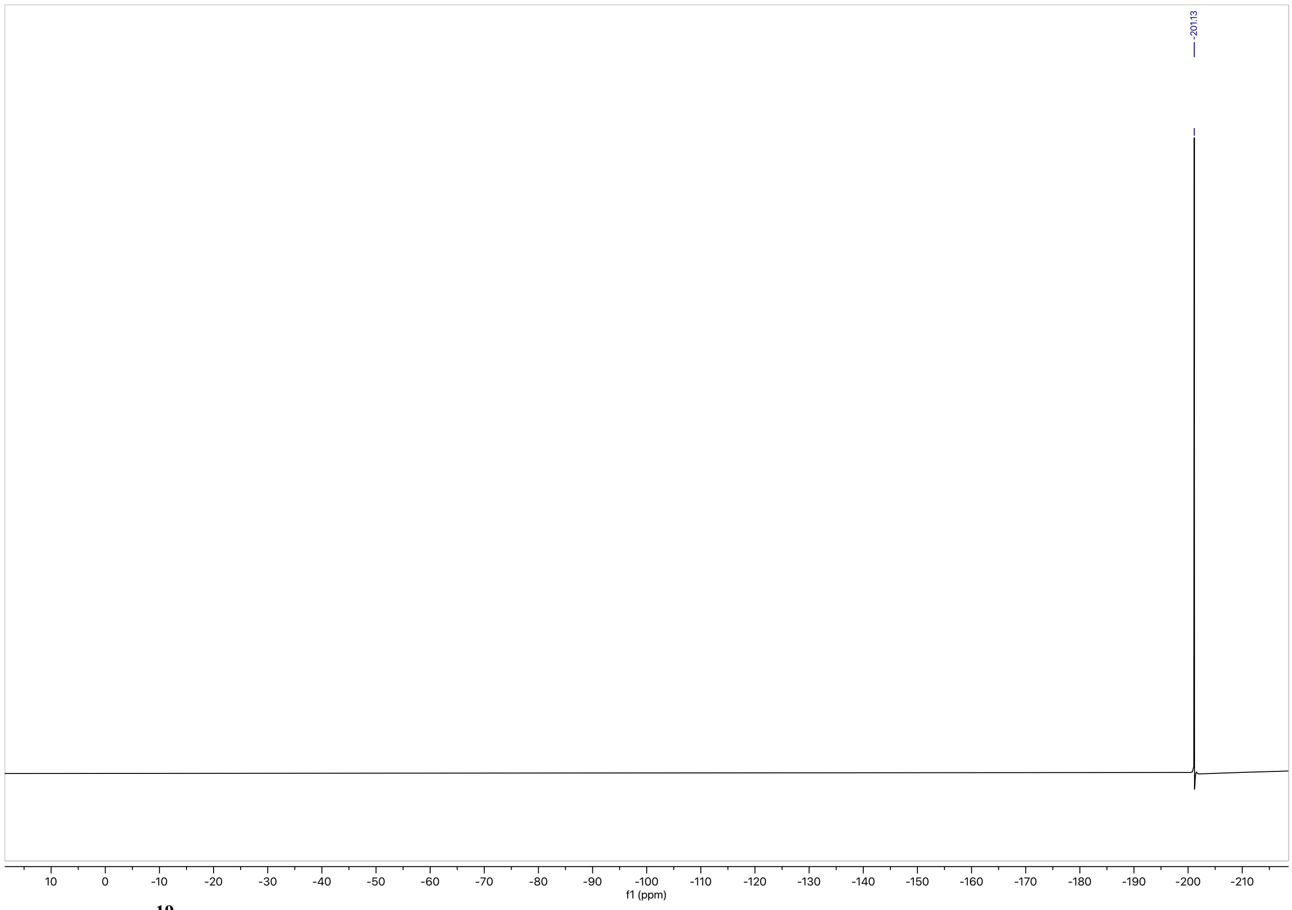


**Figure S15.** $^{19}$F NMR 2-fluoro-2-deoxy-trehalose (2)

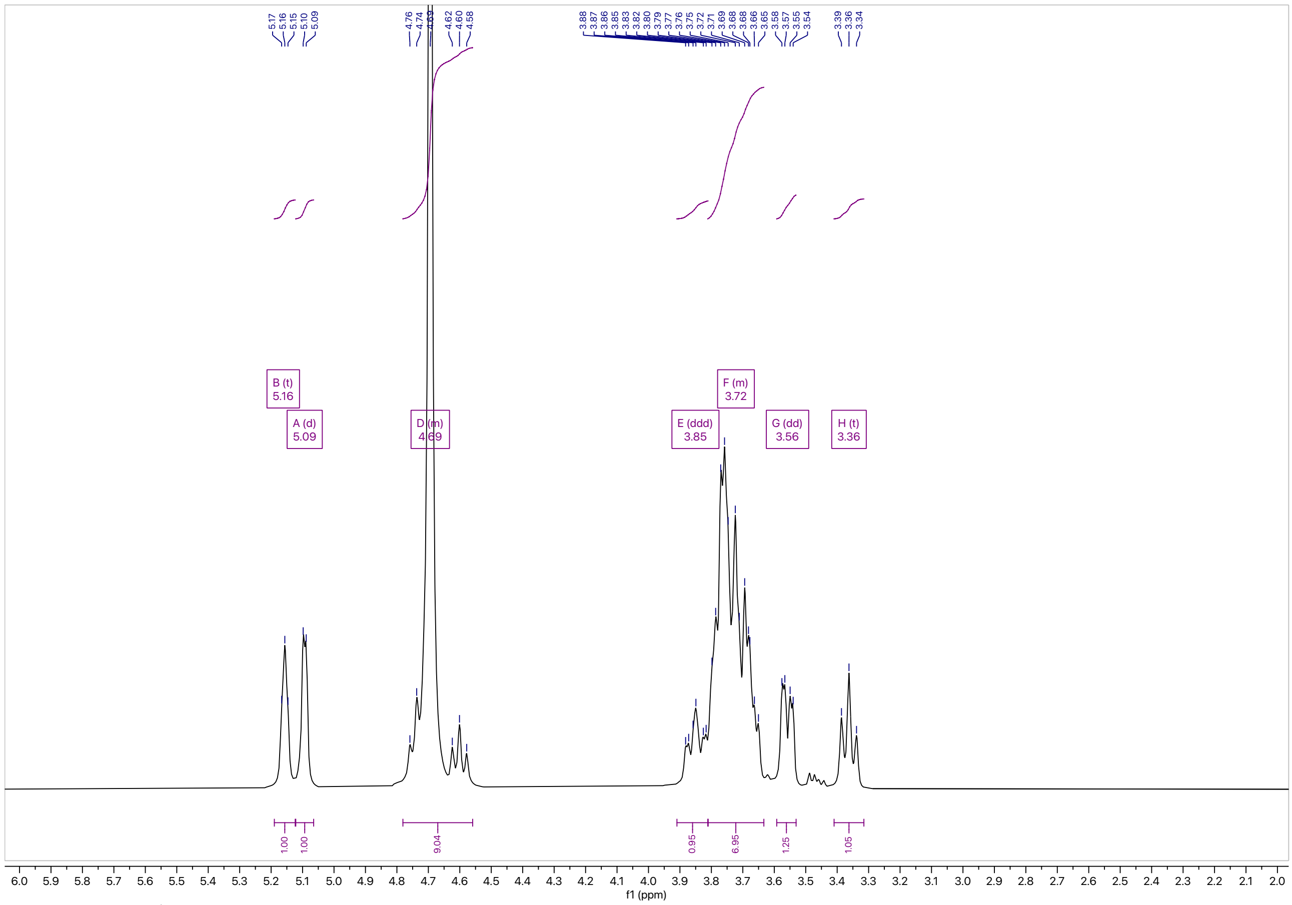


**Figure S16.** $^{1}$H NMR 3-fluoro-3-deoxy-trehalose (3)

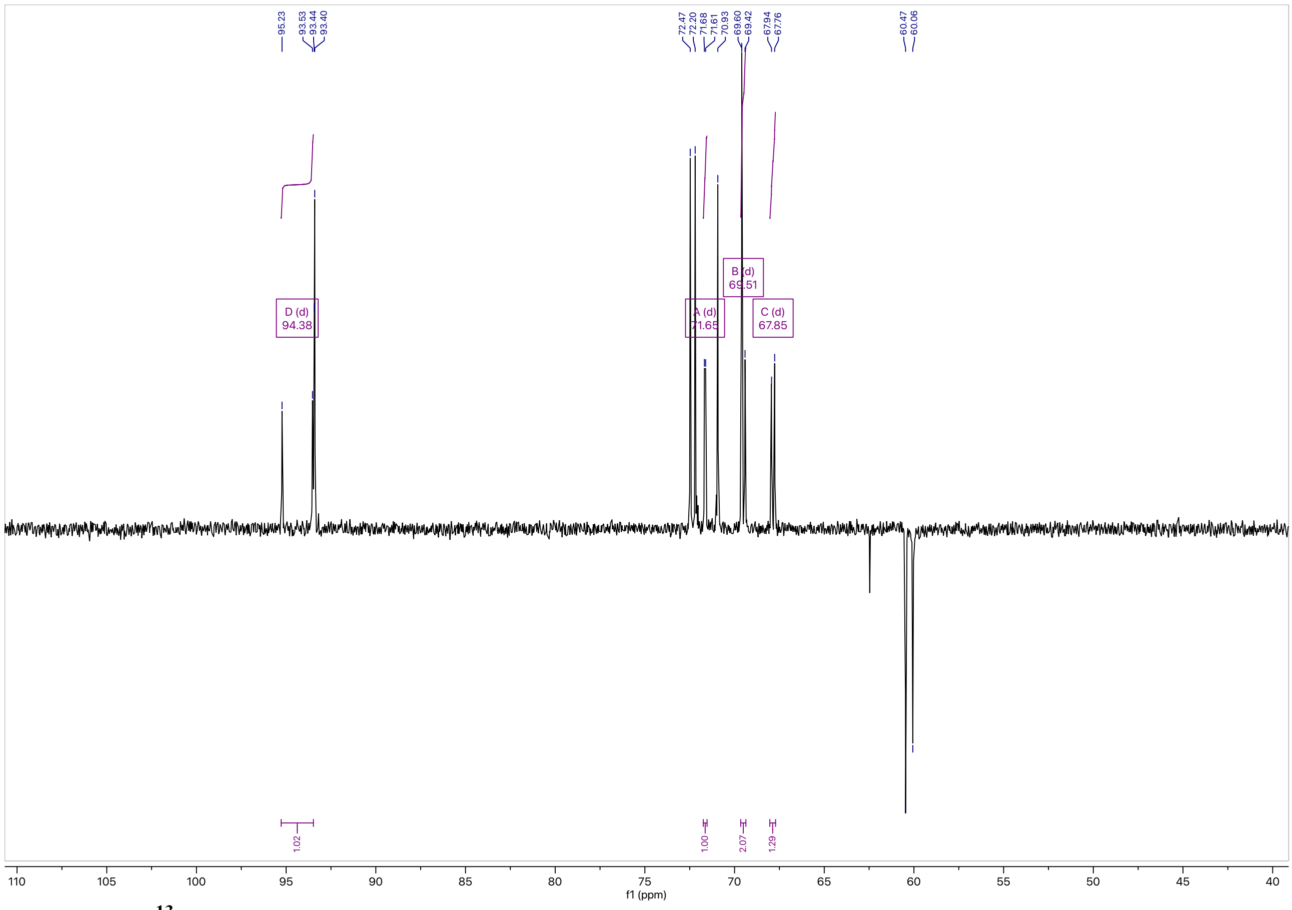


**Figure S17. $^{13}$C NMR 3-fluoro-3-deoxy-trehalose(3)**

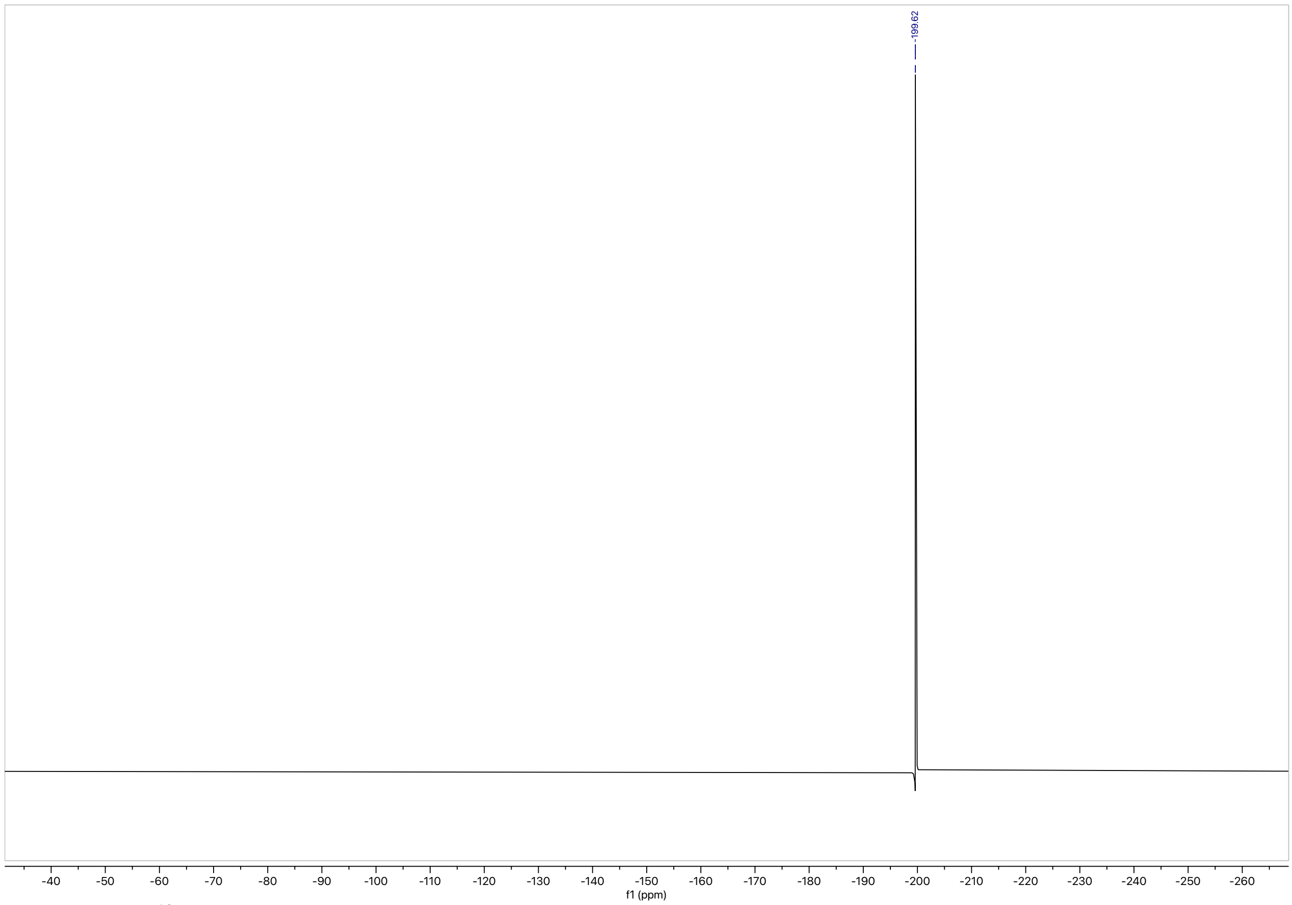


**Figure S18. $^{19}$F NMR 3-fluoro-3-deoxy-trehalose (3)**

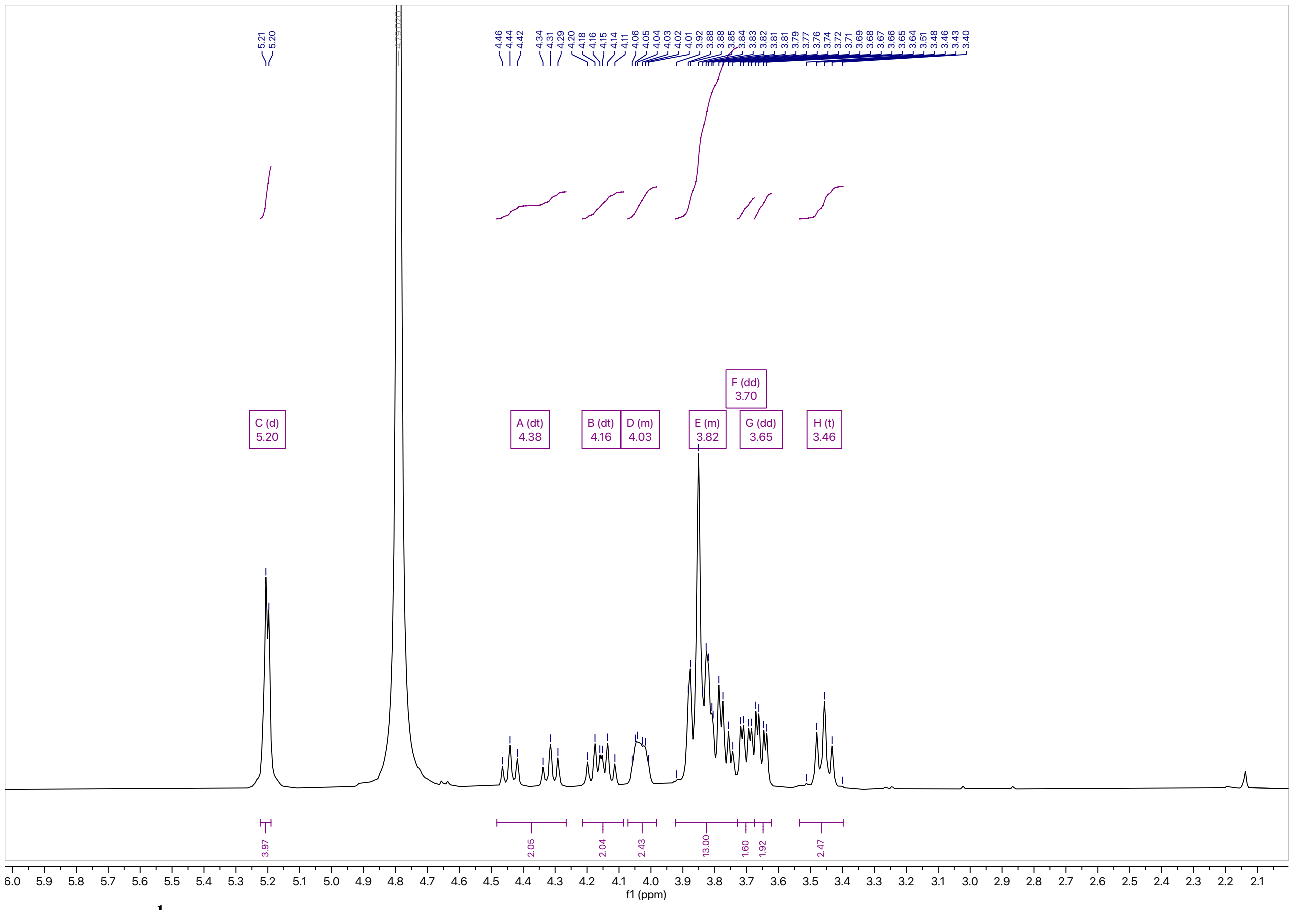

**Figure S19. [1]H NMR of 4-fluoro-4-deoxy-trehalose (4)**

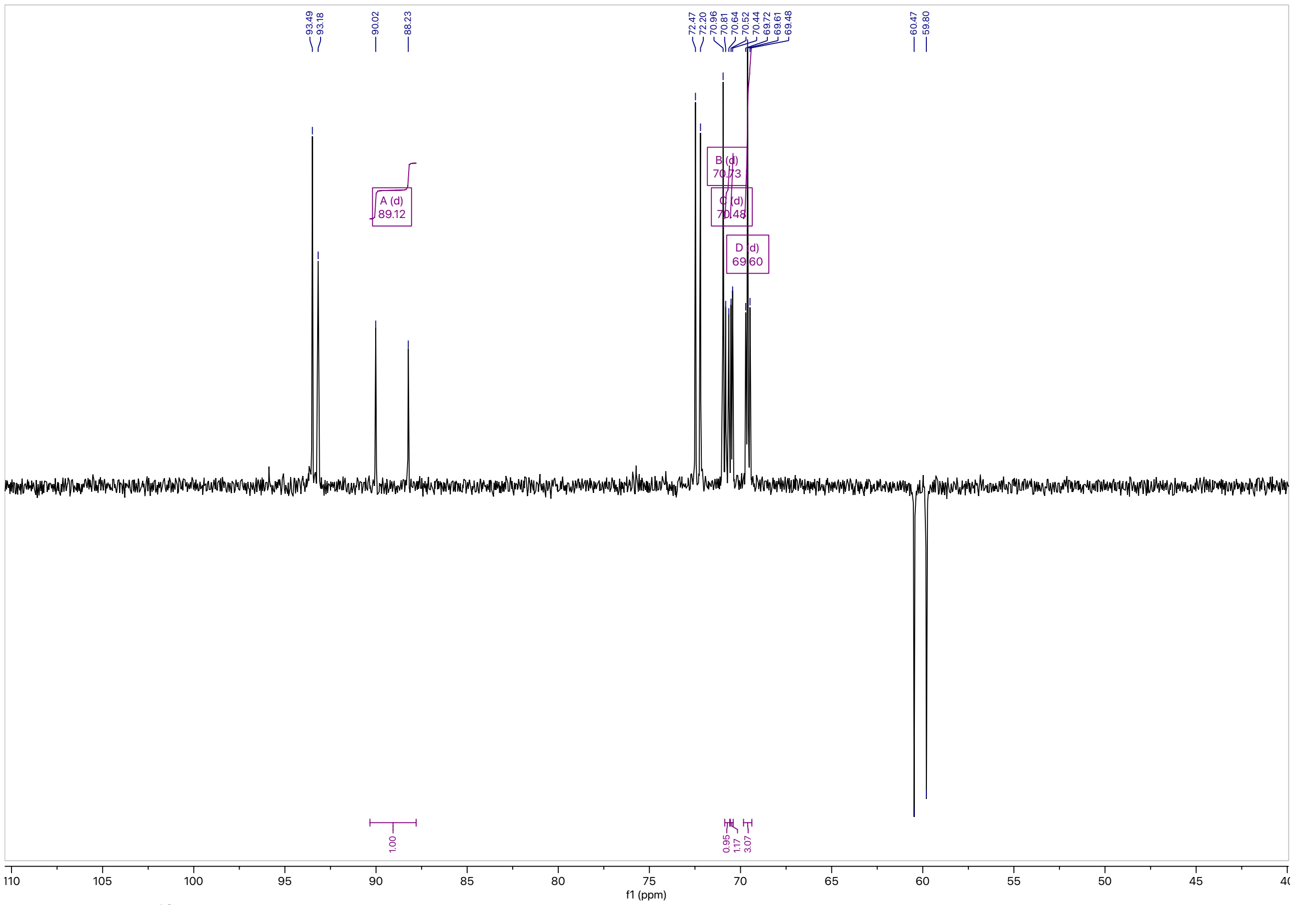

**Figure S20. [13]C NMR of 4-fluoro-4-deoxy-trehalose (4)**

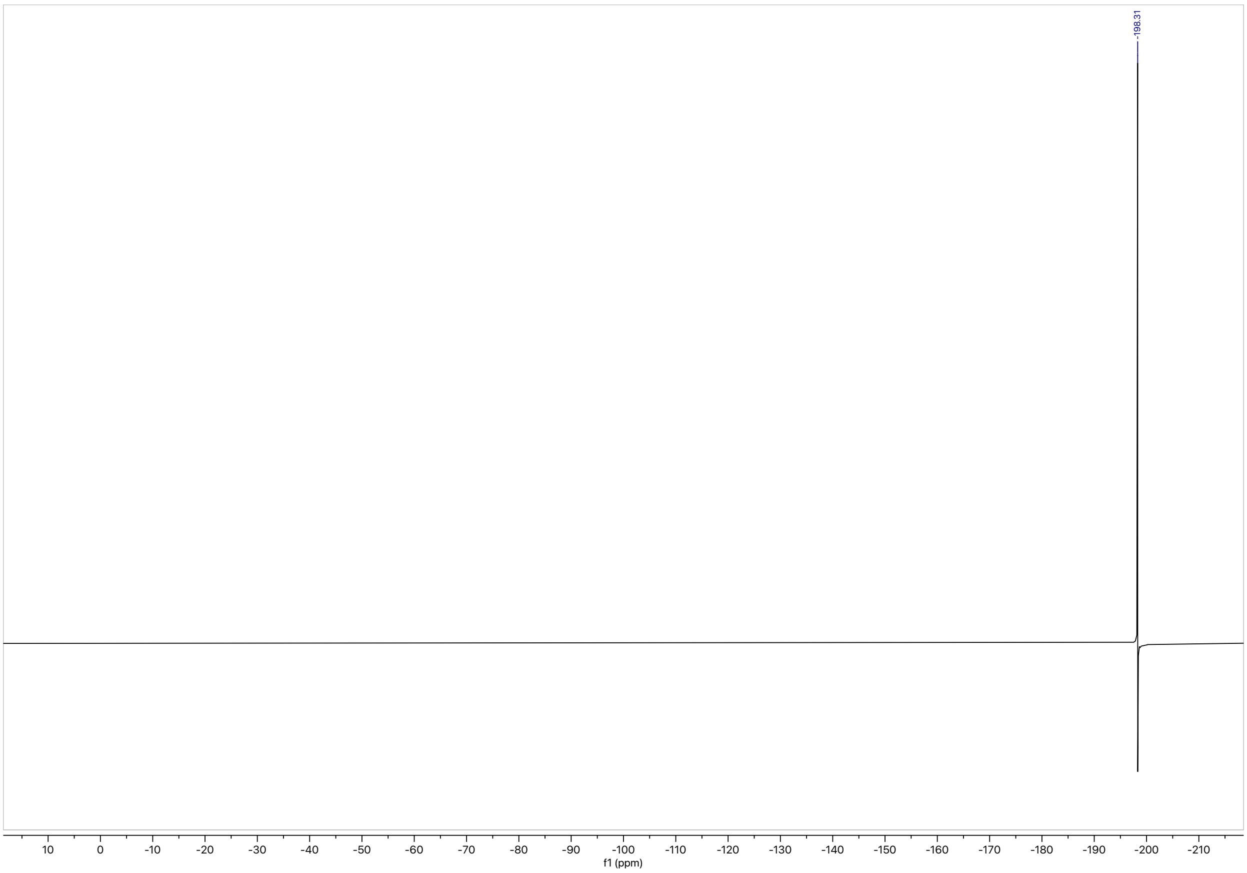


**Figure S21.** $^{19}$F NMR of 4-fluoro-4-deoxy-trehalose (4)

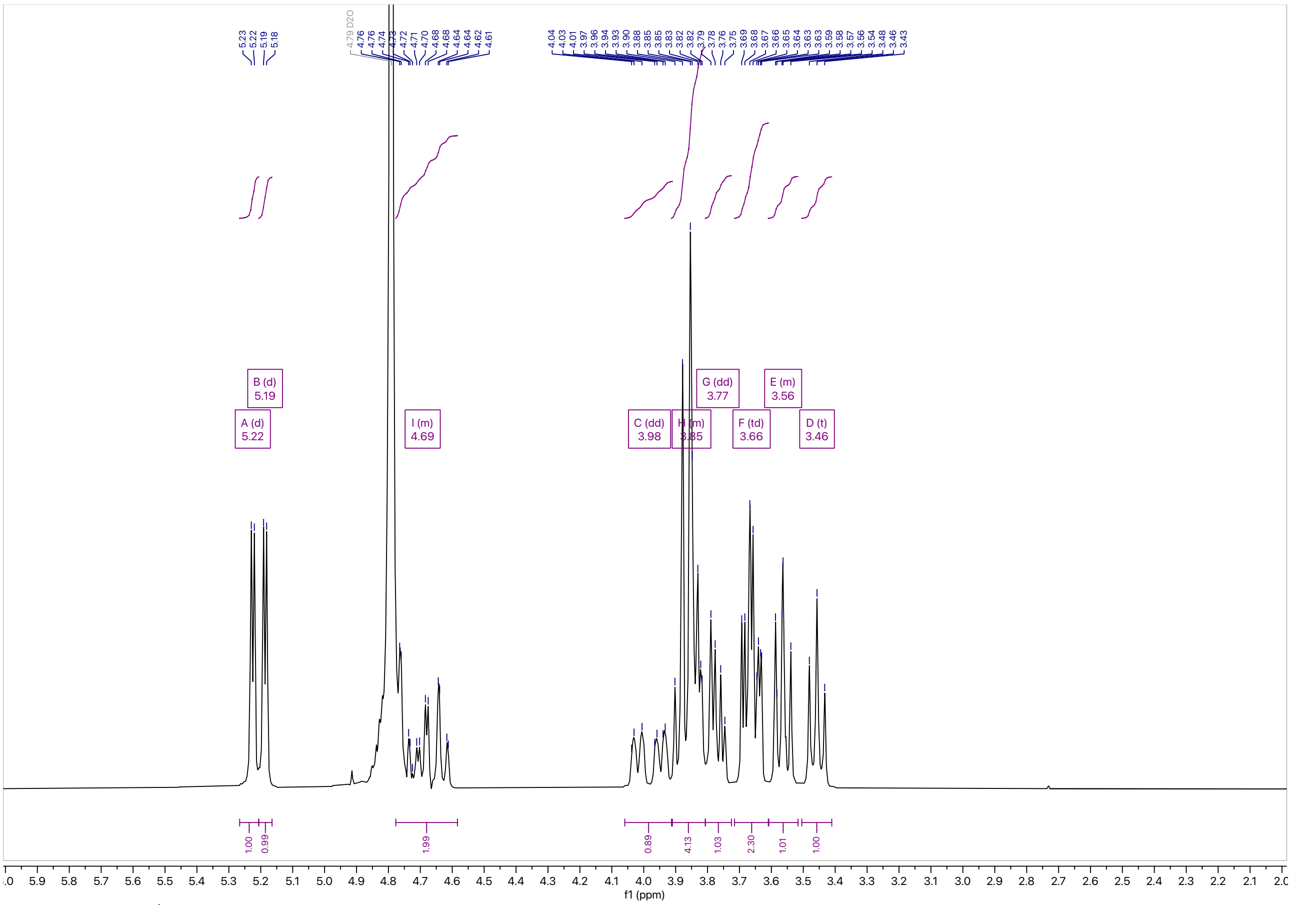


**Figure S22.** $^{1}$H NMR of 6-fluoro-6-deoxy-trehalose (5)

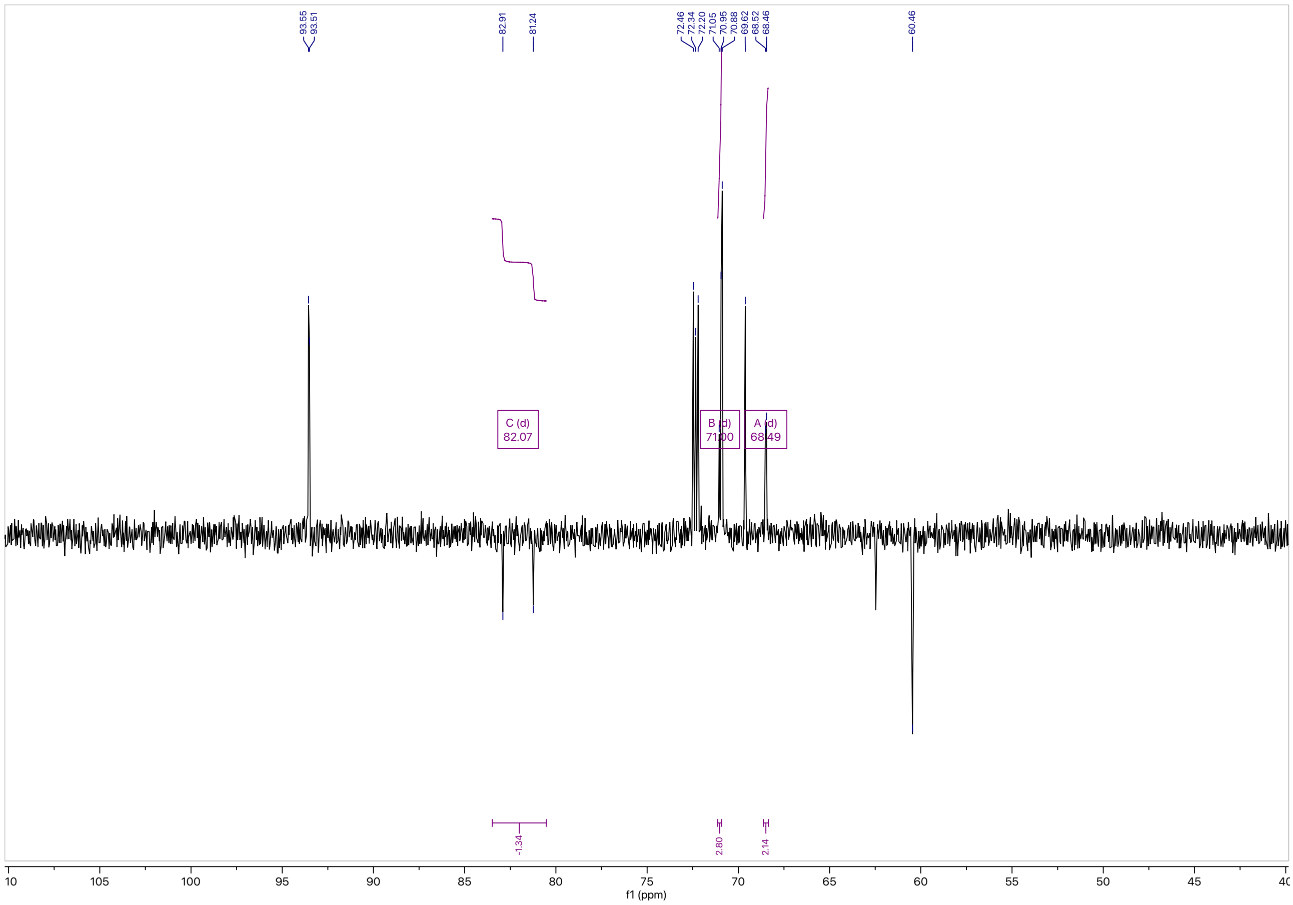


**Figure S23. $^{13}$C NMR of 6-fluoro-6-deoxy-trehalose (5)**

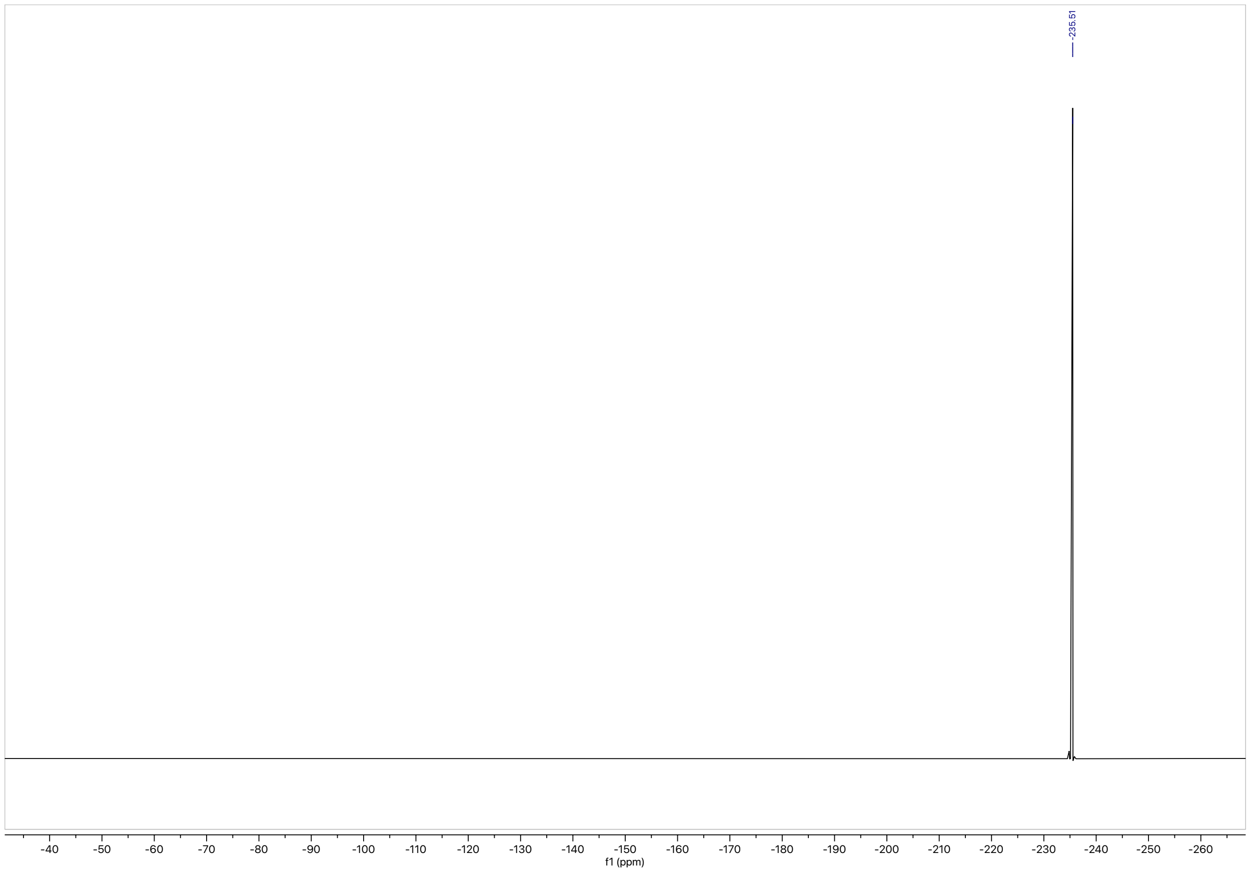


**Figure S24. $^{19}$F NMR of 6-fluoro-6-deoxy-trehalose (5)**

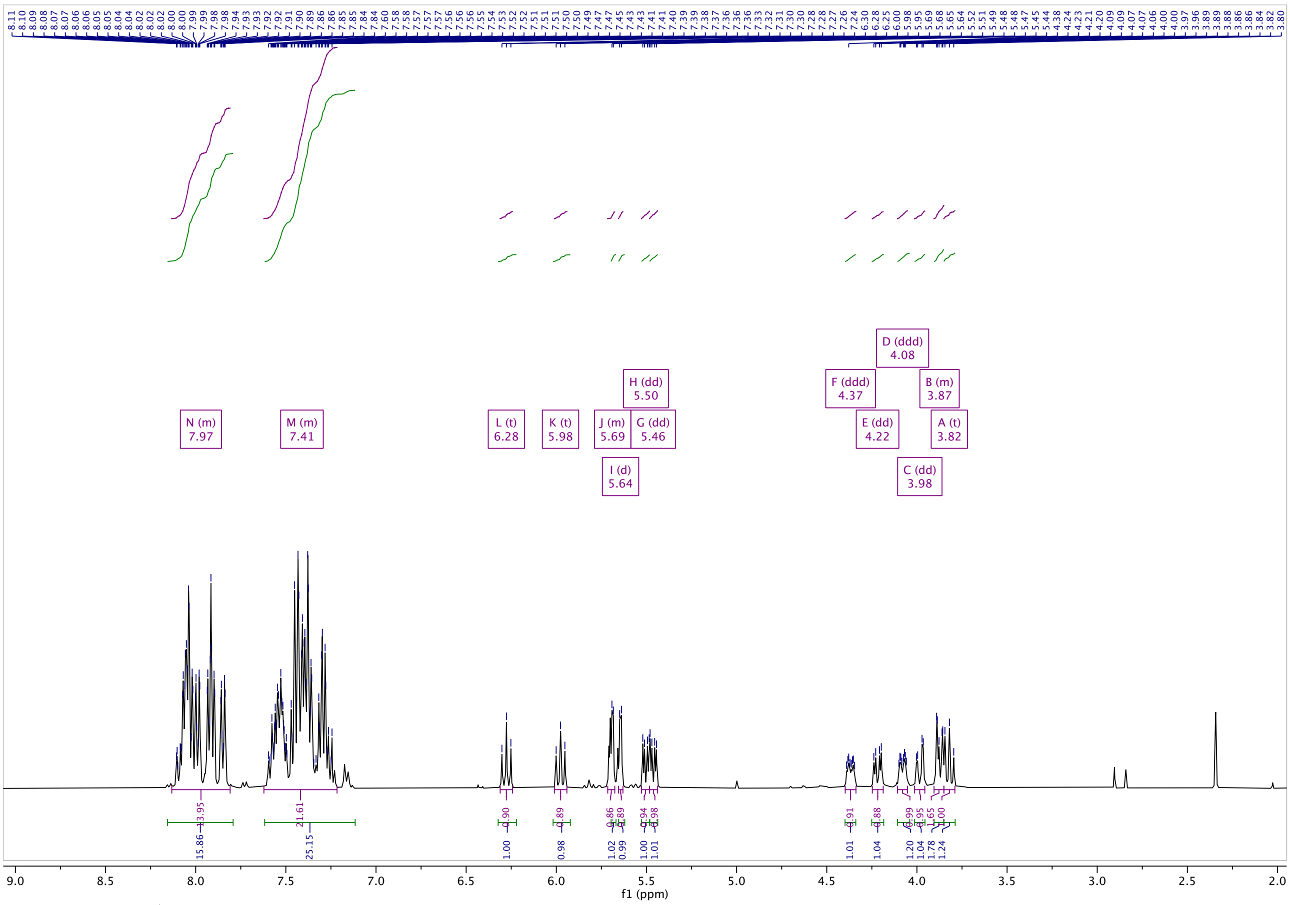


**Figure S25.** $^{1}$**H NMR of 2,3,6,2′,3′,4′,6’,-hepta-*O*-benzoyl-α,α’-D-trehalose (6)**

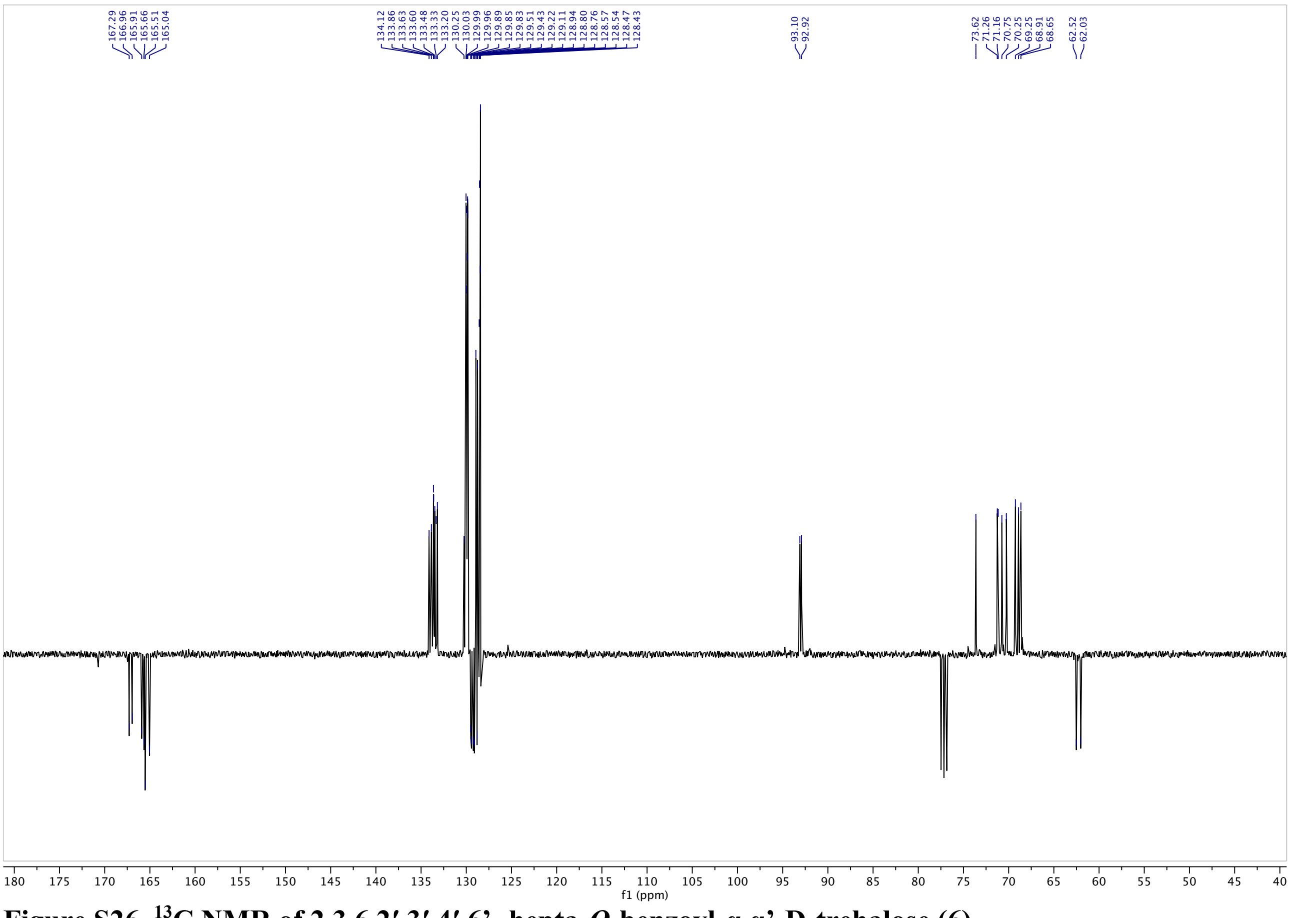


**Figure S26.** $^{13}$**C NMR of 2,3,6,2′,3′,4′,6’,-hepta-*O*-benzoyl-α,α’-D-trehalose (6)**

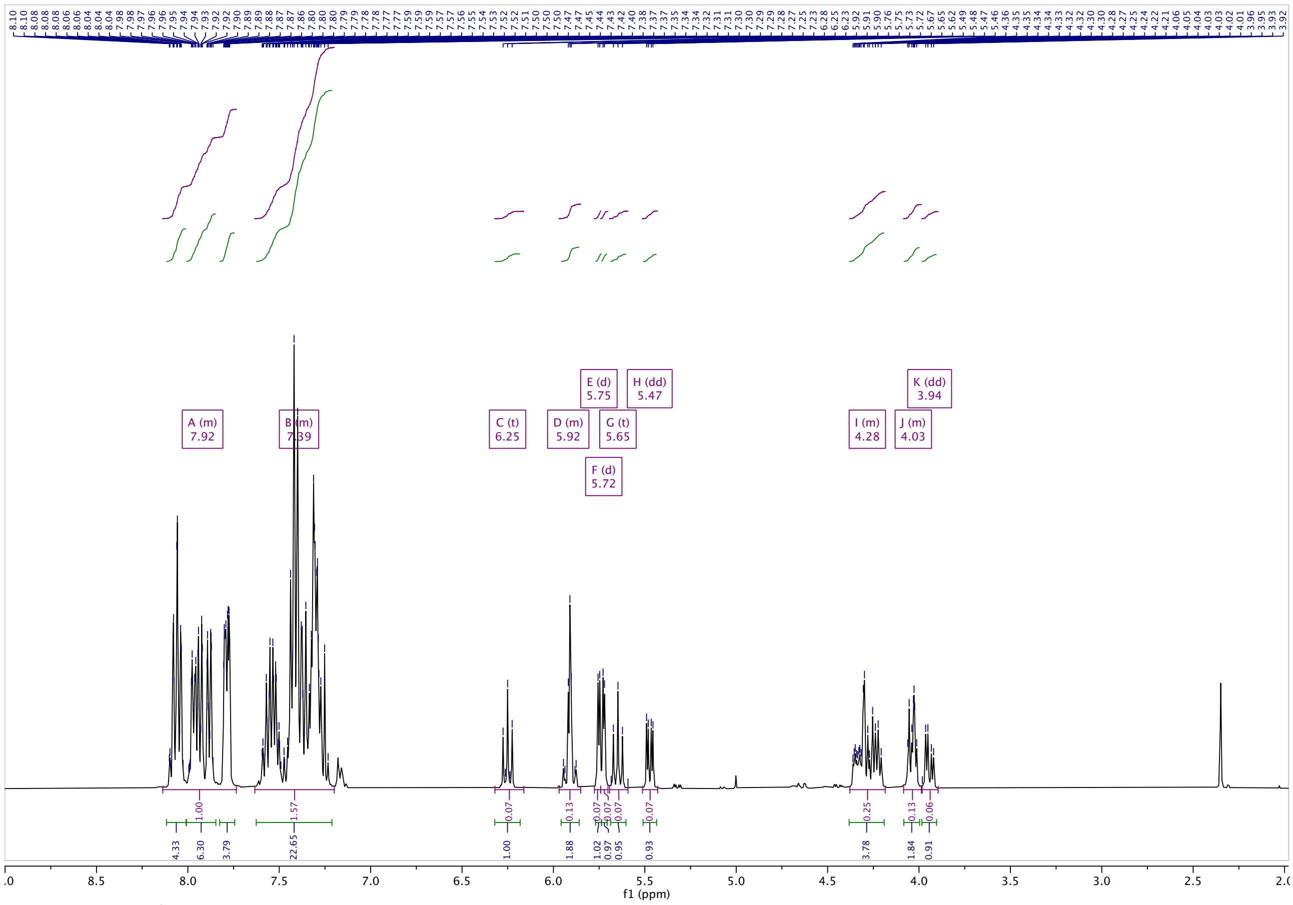


**Figure S27.** $^{1}$H NMR of 2,3,6,-tri-*O*-benzoyl-α-D-galactopyranosyl-(1→1)-2′,3′,4′,6′,-tetra-*O*-benzoyl-α-D-glucopyranoside (8)

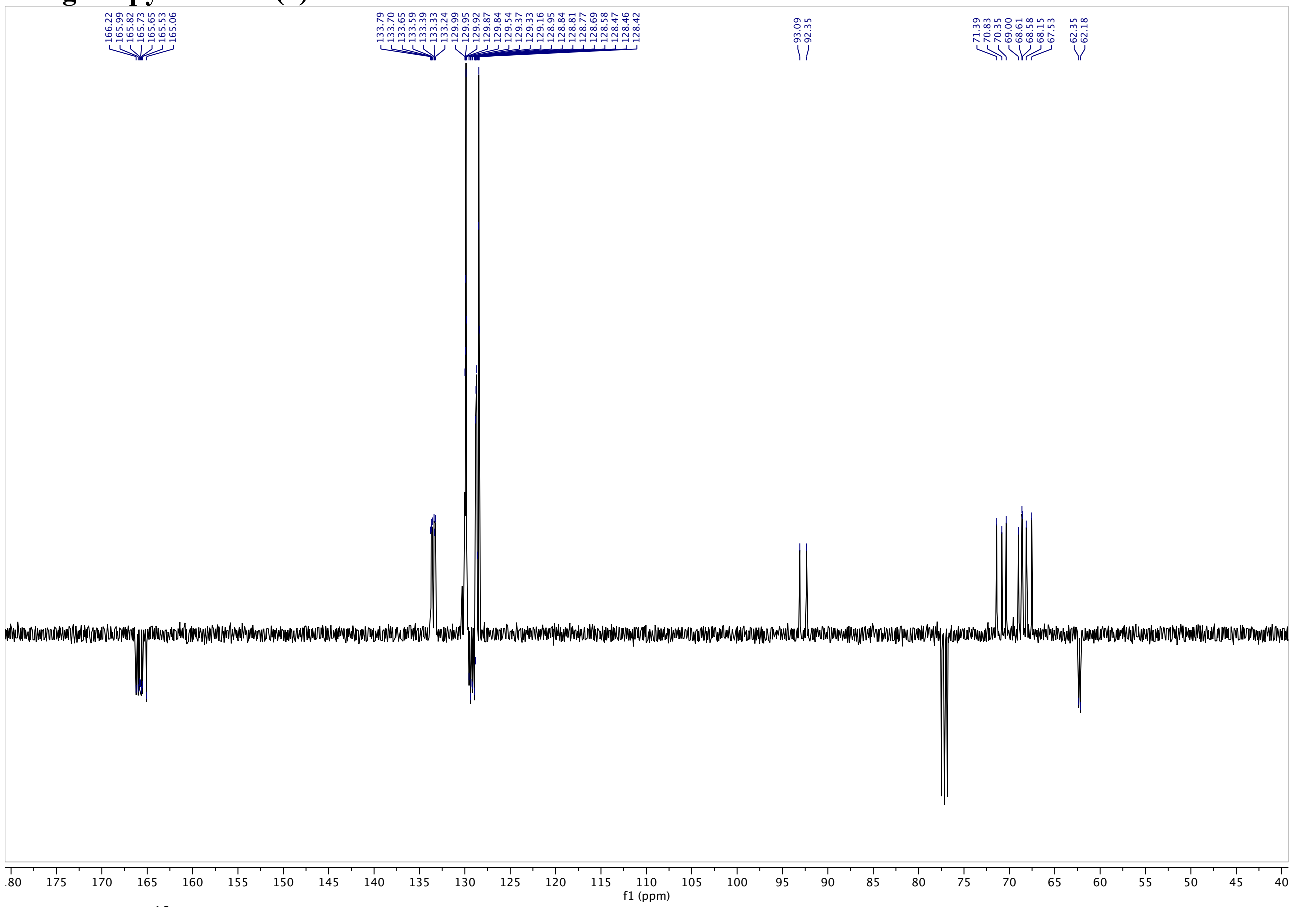


**Figure S28.** $^{13}$C NMR of 2,3,6,-tri-*O*-benzoyl-α-D-galactopyranosyl-(1→1)-2′,3′,4′,6′,-tetra-*O*-benzoyl-α-D-glucopyranoside (8)

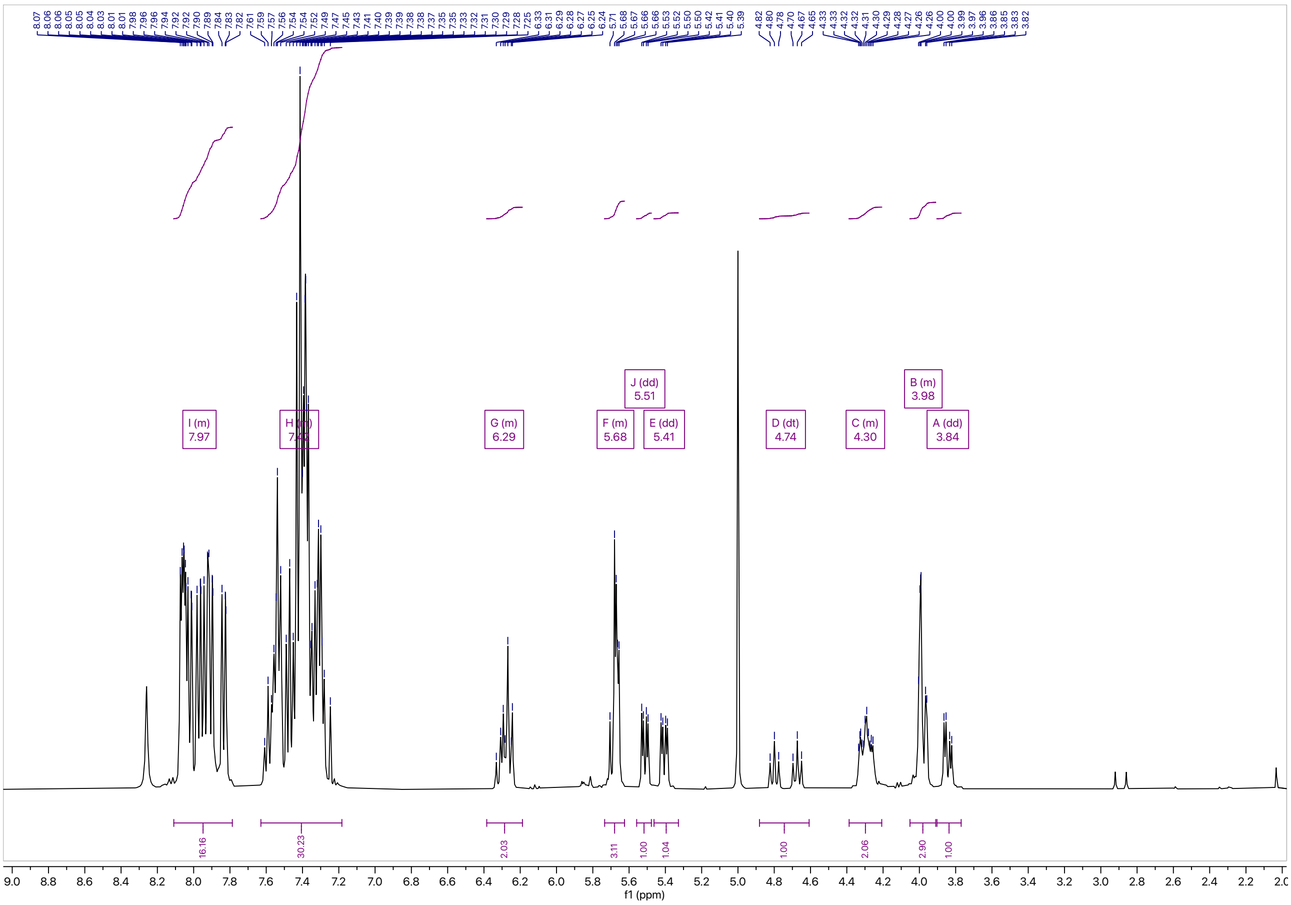


**Figure S29.** **[1]H NMR of 4-fluoro-2,3,6,-tri-*O*-benzoyl-α-D-galactopyranosyl-(1→1)-2′,3′,4′,6′,-tetra-*O*-benzoyl-α-D-glucopyranoside (9)**

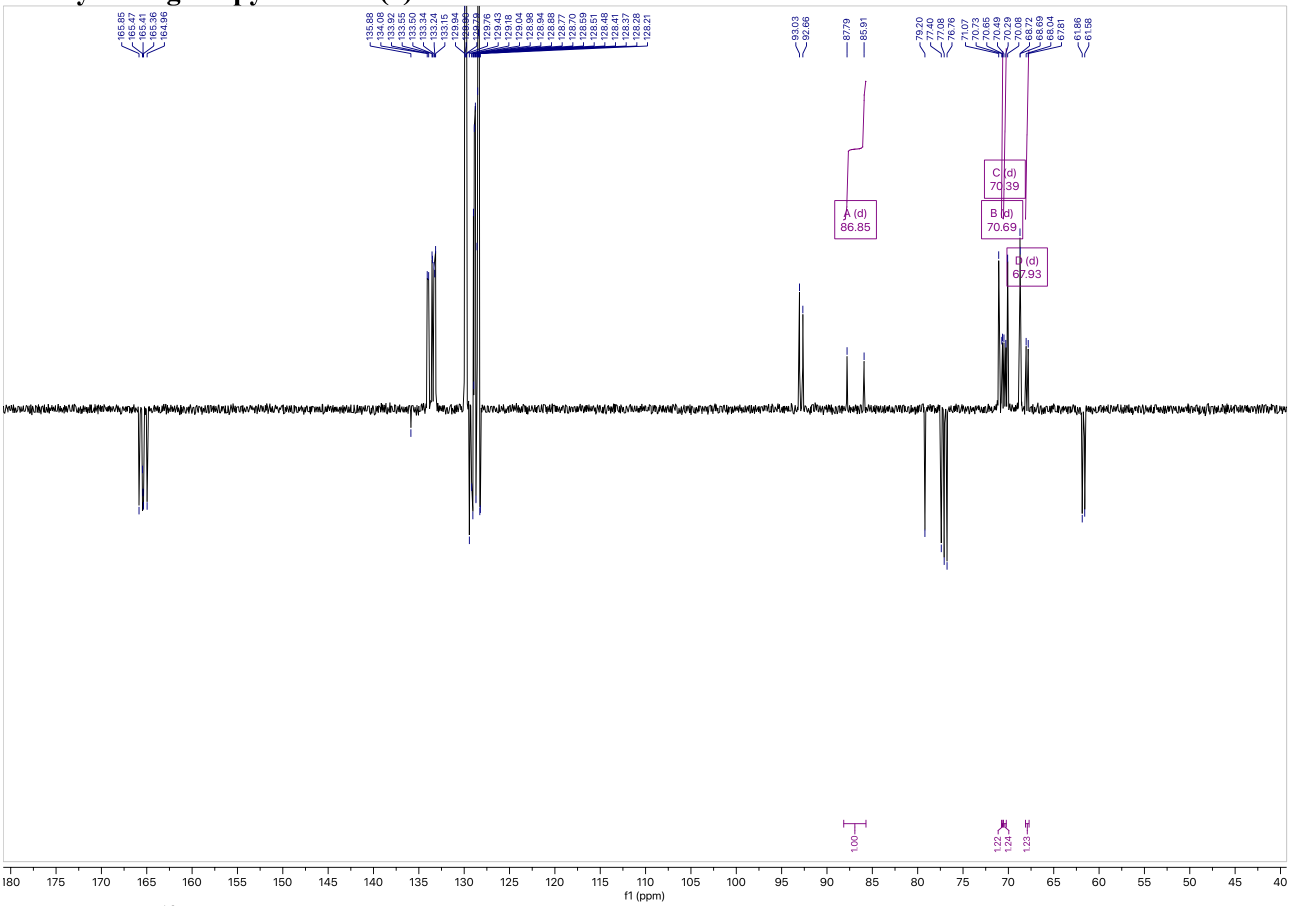


**Figure S30.** **[13]C NMR of 4-fluoro-2,3,6,-tri-*O*-benzoyl-α-D-galactopyranosyl-(1→1)-2′,3′,4′,6′,-tetra-*O*-benzoyl-α-D-glucopyranoside (9)**

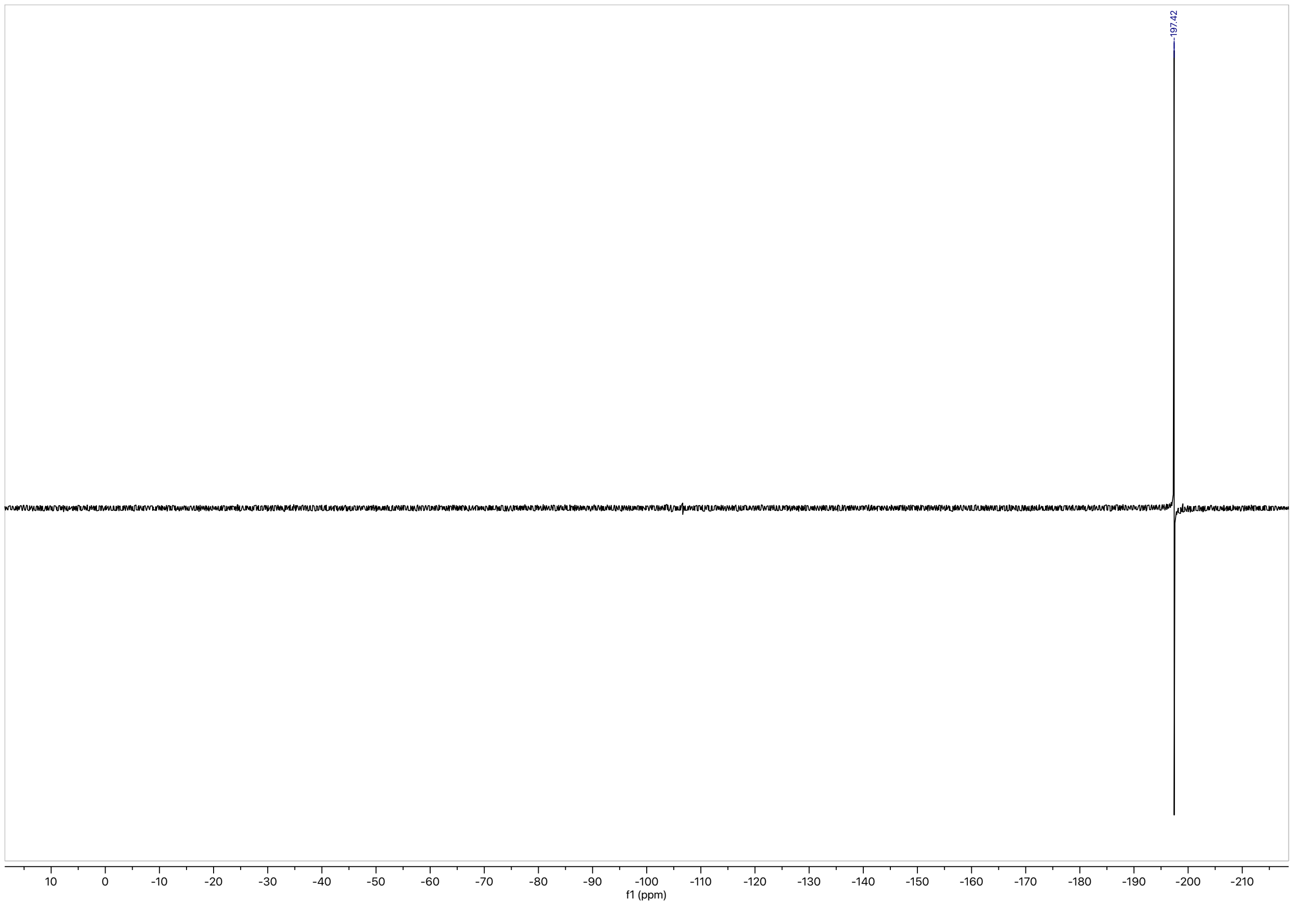


**Figure S31. $^{19}$F NMR of 4-fluoro-2,3,6,-tri-*O*-benzoyl-α-D-galactopyranosyl-(1→1)-2′,3′,4′,6′,-tetra-*O*-benzoyl-α-D-glucopyranoside (9)**
